# Immune and environment-driven gene expression during invasion: An eco-immunological application of RNA-Seq

**DOI:** 10.1101/583617

**Authors:** D Selechnik, MF Richardson, R Shine, GP Brown, LA Rollins

## Abstract

Host-pathogen dynamics change rapidly during a biological invasion and are predicted to impose strong selection on immune function. The invader may experience an abrupt reduction in pathogen-mediated selection (‘enemy release’), thereby favoring decreased investment into ‘costly’ immune responses, and the extent of this reduction may depend on factors such as propagule size. Across plants and animals, there is mixed support for this prediction. Pathogens are not the only form of selection imposed on invaders; differences in abiotic environmental conditions between native and introduced ranges are also expected to drive rapid evolution. Here, we assess the expression patterns of immune and environmentally-associated genes in the cane toad (*Rhinella marina*) across its invasive Australian range. Transcripts encoding mediators of costly immune responses (inflammation, cytotoxicity) showed a curvilinear relationship with invasion history, with highest expression in toads from oldest and newest colonized areas. This pattern is surprising given theoretical expectations of density dynamics in invasive species, and may be because density influences both intraspecific competition and parasite transmission, generating conflicting effects on the strength of immune responses. Alternatively, this expression pattern may be the result of other evolutionary forces, such as spatial sorting and genetic drift, working simultaneously with natural selection. Our findings do not support predictions about immune function based on the enemy release hypothesis, and suggest instead that the effects of enemy release are difficult to isolate in wild populations. Additionally, expression patterns of genes underlying putatively environmentally-associated traits are consistent with previous genetic studies, providing further support that Australian cane toads have adapted to novel abiotic challenges.

## Introduction

Invasive species pose a massive threat to biodiversity (Bax et al., 2003; Clavero et al., 2009). The potential for pathogens to limit the impact of invaders, or to exacerbate that impact, makes it critical to understand the effects of host-pathogen dynamics on invader immunity. The enemy release hypothesis (ERH) predicts that the processes of introduction and range expansion decrease rates of infections with co-evolved pathogens and parasites in invasive hosts due to the former’s inability to spread effectively and persist in novel environmental conditions (Colautti et al., 2004). Because of this, Lee & Klasing (2004) predict that invaders may down-regulate powerful immune responses such as systemic inflammation due to a decreased need (Cornet et al., 2016; Lee & Klasing, 2004; Martin et al., 2010). Such immune responses are also costly due to energetic expenditure (the reduction of nutrients available for partitioning across tissues due to their use in mounting immune responses (Klasing & Leshchinsky, 1999)) and to the potential for collateral damage (tissue injury due to the effects of the immune response (Martin et al., 2010)). Reduced energetic investment into these immune responses may enhance invasion success (McKean & Lazzaro, 2011). Nonetheless, loss of immunocompetence (the ability to mount a normal immune response after exposure to an antigen (Janeway et al., 2001)) could render invaders susceptible to infection by novel pathogens and parasites in their introduced range (Cornet et al., 2016; Lee & Klasing, 2004). Thus, invaders are predicted to exhibit lower investment in costly (but not all) immune responses than are seen in their native ranges (Cornet et al., 2016; Lee & Klasing, 2004).

The predicted consequences of enemy release, as well as other adaptive trait variation in invaders (Colautti & Barrett, 2013; Oduor et al., 2016; White et al., 2013), result from natural selection. However, some traits follow patterns that are not explained by selection (Berthouly-Salazar et al., 2012; Lowe et al., 2015). This may be because the dispersive tendencies of invaders give rise to additional evolutionary forces: 1) As an invasive population expands, genetic drift may reduce genetic diversity across the range (Rollins et al., 2009) and modify phenotypic traits. 2) An expanding invasion front is dominated by individuals with the highest rates of dispersal simply because the fastest arrive at new areas first and can only breed with each other (spatial sorting) (Shine et al., 2011). Thus, a geographic separation of phenotypes occurs; traits that enhance individual fitness are favored in established populations, and traits that enhance dispersal rate are common in expanding populations (Hudson et al., 2016; Shine et al., 2011). 3) Admixture between individuals from different introductions or sources, as well as hybridization, may also drive change in some invasive populations (Mader et al., 2016).

We examined the effects of range expansion on expression of immune and environmentally-associated genes in the invasive Australian cane toad (*Rhinella marina*) using RNA-Seq data from whole spleen tissue from individuals collected from long-established areas in Queensland (QLD, the ‘range core’), geographically ‘intermediate’ areas in the Northern Territory (NT), and the leading edge of the range expansion in Western Australia (WA, the ‘invasion front’) (Figure 1). The invasive range includes highly varied environments; climatic conditions in the range core are similar to those in the native range (Central and South America), but intermediate areas and the invasion front receive much less annual rainfall (2000-3000 mm in QLD, 400-1000 mm in NT and WA) and have higher annual mean temperatures (21-24°C in QLD, 24-27°C in NT and WA) (Bureau of Meteorology, 2018). Toads cluster genetically based on these environmental patterns: toads from the range core are genetically distinct from those from intermediate areas and the invasion front (Selechnik et al., 2019). Furthermore, loci putatively under selection are involved in tolerance of temperature extremes and dehydration (Selechnik et al., 2019). Traits such as locomotor performance at high temperatures also follow this pattern (Kosmala et al., 2018), but others do not. For example, behavioral propensity for exploration increases with distance from the introduction site (Gruber et al., 2017). Traits such as leg length (Hudson et al., 2016), spleen size, and fat body mass (Brown et al., 2015b) follow a U-shaped (curvilinear) pattern across the range, in which they are smallest in toads from intermediate areas of the range and larger at either end.

**Figure 1.**
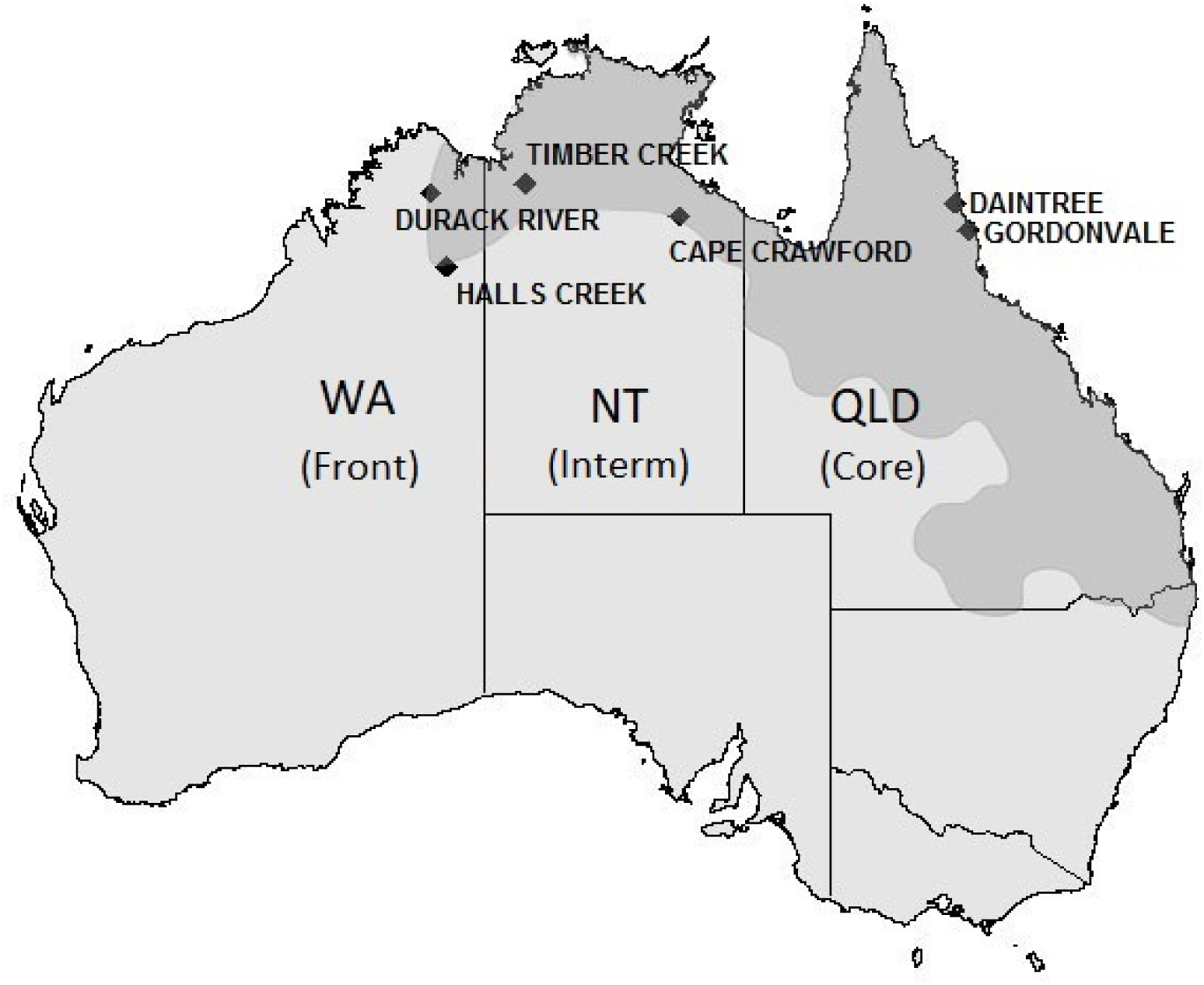
Geographic distribution of the cane toad in Australia (dark grey region). Since arriving in Queensland in 1935, cane toads have further expanded their range through New South Wales, the Northern Territory, and into Western Australia. Black diamonds indicate our toad collection sites (from east to west): QLD (Gordonvale and Daintree, N=5 each), NT (Cape Crawford and Timber Creek, N=4 each) and WA (Caroline Pool and Durack River, N=5 each). Map adapted from Tingley et al (2017) (Tingley et al., 2017).

Consistent with the ERH, many species of bacteria, protozoan, and metazoan parasites of cane toads seem to have been left behind in the native range (Selechnik et al., 2017a). A major parasite (lungworm *Rhabdias pseudosphaerocephala*) from the native range that infects toad populations in the Australian range core is absent from toads at the invasion front (Phillips et al., 2010); this lungworm has been reported to increase mortality by 17% in metamorph cane toads (Kelehear et al., 2009) and 6% in adult cane toads (Finnerty et al., 2017). Conversely, the Australian soil bacterium *Brucella* (*Ochrobactrum*) *anthropi* causes spinal spondylosis in toads primarily at the invasion front (Brown et al., 2007), which may represent a novel infection that forces invaders to remain immunocompetent. Furthermore, Rhimavirus A has only been detected in transcriptomes of toads from areas relatively close to the invasion front (Russo et al., 2018). Invasion history has complex effects on toad immunity (Brown et al., 2015c; Brown & Shine, 2014; Selechnik et al., 2017b).

Here, we aimed to characterize patterns of gene expression in toads across their expanding Australian range through differential expression (DE) analysis of transcriptome data. In terms of the ERH: release from pathogens depends on a decline in pathogen transmission, as is likely when host densities are lower. The densities of many invasive populations follow a ‘travelling wave,’ in which population density is low at recently colonized areas (e.g. the invasion front), high in areas that have been colonized for several years (e.g. intermediate areas), and low at long-colonized areas (e.g. the range core) (Hilker et al., 2005; Simberloff & Gibbons, 2004). Although absolute population densities of cane toads across Australia are unknown, toads appear to follow this trend as well (Brown et al., 2013; Freeland et al., 1986), as does at least one of their major parasites (*Rhabdias pseudosphaerocephala*). This parasitic lungworm is absent from toads at the invasion front, and is most prevalent in toads from intermediate areas (Brown et al., 2015b; Phillips et al., 2010). Therefore, we predicted that the expression of immune genes may follow a curvilinear pattern, in which those encoding mediators of costly immune responses (e.g. inflammation) may be more highly expressed in toads from intermediate areas than in toads from the core or front. In terms of environmentally-associated genes, we predicted that expression patterns would depend on the functional roles of the genes; for example, genes involved in aridity tolerance may be differentially expressed between toads from the (moist) range core and toads throughout the rest of the (more arid) range.

## Materials and Methods

### Sample collection and RNA extraction

In April and May of 2014 and 2015, we collected adult female cane toads from six locations along an invasion transect (Figure 1): Gordonvale, QLD (N=5, range core, 17.0972S 145.7792E); Daintree, QLD (N=5, range core, 16.25S 145.3167E); Cape Crawford, NT (N=4, intermediate, 16.6667S 135.8E); Timber Creek, NT (N=4, intermediate, 15.6453S 130.4744E); Halls Creek, WA (N=5, invasion front, 18.2265S 127.759E); and Durack River, WA (N=5, invasion front, 15.9419S 127.2202E). We euthanized toads using 150 mg/kg sodium pentobarbital, decapitated them as soon as they became unresponsive, and excised their spleens immediately following decapitation. We selected spleen tissue for our investigation because mature immune cells travel to secondary lymphoid tissue (spleen, lymph nodes, and MALT) for activation through pathogenic encounter (Janeway et al., 2001). Thus, the cellular compositions of these tissue should reflect the host’s immune functioning. Each spleen was initially preserved in RNAlater (QIAGEN, USA), kept at 4°C for less than one week, and then drained and transferred to a −80°C freezer for long-term storage.

Prior to RNA extraction, we flash-froze all spleens individually in liquid nitrogen and ground them with a mortar and pestle to lyse preserved tissue. We carried out RNA extractions using the RNeasy Lipid Tissue Mini Kit (QIAGEN, USA) following the manufacturer’s instructions, with an additional genomic DNA removal step using on-column RNase-free DNase treatment (QIAGEN, USA). We quantified the total RNA extracted using a Qubit RNA HS assay on a Qubit 3.0 fluorometer (Life Technologies, USA). Extracts were then stored at −80°C until sequencing was performed.

### Sequencing

Prior to sequencing, we added 4 µL of either mix 1 or mix 2 of External RNA Controls Consortium (ERCC; Thermo Fisher Science) spike-in solutions diluted 1:100 to 2 µg of RNA to examine the technical performance of sequencing (Table S1). Macrogen (Macrogen Inc., ROK) constructed mRNA libraries using the TruSeq mRNA v2 sample kit (Illumina Inc., USA), which included a 300bp selection step. All samples from the core and the front were sequenced in the same batch (but not pooled), and evenly distributed, across two lanes of Illumina HiSeq 2500 (Illumina Inc., USA); samples from intermediate areas were sequenced in a separate batch on a single lane (also not pooled). Capture of mRNA was performed using the oligo dT method, and size selection parameter choices were made according to the HiSeq2500 manufacturer’s protocol. Each individual sample was ligated with a unique barcode. Overall, this generated 678 million paired-end 2 x 125 bp reads. Raw sequence reads are available as FASTQ files in the NCBI short read archive (SRA) under the BioProject Accession PRJNA395127.

### Data pre-processing, alignment, and expression quantification

First, we examined per base raw sequence read quality (Phred scores) and GC content, and checked for the presence of adapter sequences for each sample using FastQC v0.11.5 (Andrews, 2010). We then processed raw reads (FASTQ files) from each sample with Trimmomatic v0.35 (Bolger et al., 2014), using the following parameters: ILLUMINACLIP:TruSeq3-PE.fa:2:30:10:4 SLIDINGWINDOW:5:20 AVGQUAL:20 MINLEN:36. This removed any adaptor sequences, trimmed any set of 5 contiguous bases with an average Phred score below 20, and removed any read with an average Phred score below 20 or sequence length below 36 bp.

As a reference for alignment, we used the annotated *R. marina* transcriptome (Richardson et al., 2018), which was constructed from brain, spleen, muscle, liver, ovary, testes, and tadpole tissue. We conducted per sample alignments of our trimmed FASTQ files to this reference using STAR v2.5.0a (Patro et al., 2017) in basic two-pass mode with default parameters, a runRNGseed of 777, and specifying BAM alignment outputs. We used the BAM outputs to quantify transcript expression using Salmon v0.8.1 (Patro et al., 2017) in alignment mode with libtype=IU, thus producing count files.

### Count filtering and log-ratio transformations

Most methods for analyzing RNA-Seq expression data assume that raw read counts represent absolute abundances (Quinn et al., 2017a). However, RNA-Seq count data are not absolute and instead represent relative abundances as a type of compositional count data (Quinn et al., 2017b; Quinn et al., 2017a). Using methods that assume absolute values is invalid for compositional data (without first including a transformation) because the total number of reads (library size) generated from each sample varies based on factors such as sequencing performance, making comparisons of the actual count values between samples difficult (Fernandes et al., 2014; Quinn et al., 2017b). As such, relationships within RNA-Seq count data make more sense as ratios, either when compared to a reference or to another feature within the dataset. Hence, we analyzed our count data (from Salmon) taking the compositional nature into account using the log-ratio transformation (Aitchison & Egozcue, 2005; Erb & Notredame, 2016; Lovell et al., 2015; Quinn et al., 2018b; Quinn et al., 2017a). Our total number of expressed transcripts across all toads was 22,930. To filter out transcripts with low expression, we removed transcripts that did not have at least 10 counts in 10 samples. This reduced our list of expressed transcripts to 18,945. We then used the R (Team, 2016) package ALDEx2 v1.6.0 (Fernandes et al., 2013) to perform an inter-quantile log-ratio (iqlr) transformation of the transcripts’ counts as the denominator for the geometric mean calculation (rather than centered log-ratio transformation) because it removes the bias of transcripts with very high and low expression that may skew the geometric mean (Quinn et al., 2017a). To circumvent issues associated with other normalization methods, we used ALDEx2 to model the count values over a multinomial distribution by using 128 Monte Carlo samples to estimate the Dirichlet distribution for each sample (Fernandes et al., 2013). The Dirichlet modelling and iqlr transformation enabled us to perform valid significance tests among samples of different groups for DE analysis. This approach has been shown to be consistent with (but more conservative, i.e. fewer false positives, than) those of traditional DE analyses (Quinn et al., 2018a).

### Technical and diagnostic performance

Because the samples from intermediate areas were sequenced on a different run of the sequencing machine than the core and front samples, we needed to rule out a batch effect, in which samples from intermediate areas may have had disproportionately higher or lower numbers of reads for each transcript due to technical variation in sequencing performance during different runs. This could result in erroneous DE calls. Thus, we used the previously added ERCC control mixes (Ambion) to assess whether or not there was a batch effect due to sequencing run. Each set contains four groups of sequences with different ratios between the two mixes, representing ‘known’ differences in abundance (mix 1 versus mix 2 fold changes: 4:1, 1:1, 1:1.5, 1:2). We used the R (Team, 2016) package erccdashboard v1.10.0 (Munro et al., 2014) to analyze the counts of these sequences and generate Receiver Operator Characteristic (ROC) curves and the Area Under the Curve (AUC) statistic, lower limit of DE detection estimates (LODR), and expression ratio variability and bias measures based on these sequence abundance ratios (Figure S1). In the ROC curves, the AUC for two sets of true-positive ERCC sequences (4:1 and 1:2) was 1.0, indicating perfect diagnostic performance, and the AUC for the third (1:1.5) was 0.89, indicating good diagnostic performance (Figure S1B). The MA plot shows that the measured ratios in our ERCC sequences converge around the r_m_ corrected ratios, indicating low variability (Figure S1C). Finally, the LODR plot indicated that DE p-values were lower for ERCC sequences with wider ratios (i.e. 4:1 has the lowest p-values, then 1:2, then 1:1.5), which is expected because the most pronounced fold-change differences should yield the highest DE significance (Figure S1D). From these results, we inferred that the observed relative abundances between mixes of each set of ERCC sequences were close to the known relative abundances, and thus batch effects do not appear to have occurred.

We also examined the counts of the invariant (1:1) group across all samples. Seven invariant ERCC transcripts remained after count filtering (same as used for the DE testing); we generated a boxplot of their counts, normalized by library size (Figure S2). The consistency of the boxplot distributions of the seven invariant ERCC sequences further indicated that there was no batch effect. Because the erccdashboard package indicated that the sets of true positive ERCC sequences (4:1, 1:2, 1:1.5) existed in observed ratios close to the known ratios, and because the invariant sequences (1:1) exhibited consistency across samples, we proceeded with downstream analyses.

### Differential gene expression in discrete phases of the invasion

After applying a log-ratio transformation to the count data, we were able to implement statistical tests that would otherwise be invalid for relative data. We grouped populations by phase (Daintree and Gordonvale in QLD/the core, Cape Crawford and Timber Creek in NT/intermediate areas, and Durack River and Halls Creek in WA/the front) and used these as groups for DE analysis. We fitted our log-transformed count data to a non-parametric generalized linear model (glm) in ALDEx2. We took a ‘one vs. all’ approach, in which we compared samples from each state to samples from the other two states collectively (e.g. core vs. intermediate + front, intermediate vs. core + front, front vs. core + intermediate) using the Kruskal Wallis test. This test design allowed us to identify transcripts that were up- and down-regulated in toads from each state relative to those throughout the rest of the range. We only retained transcripts with Benjamini-Hochberg (FDR) corrected *p*-values < 0.05 (Fernandes et al., 2013). We detected 1,151 differentially expressed transcripts across all samples. We calculated the effect sizes of the differences between groups for each transcript, with positive values indicating up-regulation, and negative values indicating down-regulation. We further investigated all transcripts with effect sizes greater than 1.5 or less than −1.5.

### Spatial gene expression patterns across the range

The DE testing performed in ALDEx2 generates differences between discrete groups; however, our data are sampled across a continuous variable: space. So, to visualize expression patterns across the toad’s Australian range, we performed soft (fuzzy C-means) clustering on our log-transformed count data (with samples grouped by collection site, and sites ordered from east to west) using the R package Mfuzz v2.34.0 (Kumar & Futschik, 2007). The fuzzy C-means algorithm groups transcripts together based on similar expression patterns (using a fuzzifier parameter, *m*) across conditions to identify prominent, recurring patterns (clusters). Each transcript within a cluster is assigned a membership value, indicating how closely its expression pattern aligns with that of the cluster to which it belongs. To prevent random data from being clustered together, we used the mestimate command in the Mfuzz package to determine the optimal fuzzifier parameter value using a relation proposed for fuzzy c-means clustering (Schwämmle & Jensen, 2010). We then used the cselection and Dmin commands to determine the optimal number of clusters, *c*, to generate. The results of both tools suggested using four clusters (*c*=4); however, these tools need to be used with caution because automatic determination of the optimal value of *c* is difficult, and it is advised to review the data before choosing (Kumar & Futschik, 2007). For this reason, we manually performed repeated clustering for a range of *c* (*c*=3, 4, 5, 6, 7, 8) using the fuzzy c-means algorithm to visualize the differences in clusters of expression patterns across space, and to compare the internal cores (identities and membership values of transcripts within each cluster) across *c* values. Although internal cores were consistent across all *c* values, we determined that several uniquely shaped expression patterns were collapsed at *c*=4, and that these expression patterns only became separate at *c*=6. At *c* > 6, redundant patterns began to emerge. For this reason, we selected *c*=6 as the final value with which to perform soft clustering. We required a minimum membership value of 0.7 for all transcripts to their respective clusters.

### Environmentally-influenced gene expression

Genes affected by natural selection may have expression levels that are associated with environmental variables. We downloaded climatic data from the Bioclim database (Hijmans et al., 2005) using the raster package (Hijmans, 2015) in R. Because different areas of Australia vary in aridity, we downloaded data on rainfall during the driest quarter and maximum temperature in the warmest month; these data are averages of annual statistics over the period of 1970 to 2000. We then used the lfmm v2.0 package (Frichot & Francois, 2015) in R to perform a latent factor mixed model (LFMM) to test the association between the log-transformed count values of every expressed transcript and these two environmental variables. We applied a Benjamini-Hochberg correction to all p-values from the LFMM.

### Coordination in gene expression

To identify genes with coordinated (co-associated) expression, we calculated proportionality (ρ) between all pairs of transcripts in our dataset using the propr package (Quinn et al., 2017a). A full description of this analysis is available in the Supplementary Information.

### Annotation and gene ontology enrichment

We performed gene ontology (GO) enrichment analysis to identify the most ‘common’ (recurring) biological functions in which our transcripts were involved. This allowed us to determine whether certain functional categories from the GO database were overrepresented in our DE, fuzzy clustering (spatial expression), and proportionality (coordination and differential coordination) datasets more than would be expected by chance. We used the Bioconductor tool (Huber et al., 2015) GOseq v 1.26.0 (Young et al., 2010) because it accounts for bias introduced by variation in transcript lengths. We assessed three sets of GO categories (Biological Process, Molecular Function, and Cellular Component) for enrichment using the *Wallenius approximation* (while controlling for transcript length) to test for over-representation, and then Benjamini-Hochberg corrected the resulting *p*-values. To visualize the results, we plotted significantly enriched GO categories with REVIGO (Supek et al., 2011), which performs SimRel (Sæbø et al., 2015) semantic clustering of similar GO functions with annotations sourced from the UniProt database (Consortium, 2017). We then used GO terms to filter through the output lists from our DE, fuzzy clustering, and proportionality datasets to identify transcripts of genes involved in immune function.

### Identification of immune genes

To identify additional transcripts that have known functions in the immune system (outside of those with the largest DE effect size or cluster membership), we cross-matched the output transcript lists from all of our analyses with several lists of known immune genes within the database InnateDB (Breuer et al., 2013): Immunology Database and Analysis Portal (ImmPort), the Immunogenetic Related Information Source (IRIS), the MAPK/NFKB Network, and the Immunome Database. These databases consist entirely of human-mouse-bovine genes, but the immune systems of mammals and amphibians are broadly similar (Colombo et al., 2015; Robert & Ohta, 2009). We further investigated all transcripts within our datasets that matched a gene within the InnateDB gene lists.

### Isolation by distance

To assess the effect of geographic distance on divergence in gene expression (thereby testing for isolation by distance), we performed a Mantel test using the ade4 package (Thioulouse & Dray, 2007). A full description of this analysis is available in the Supplementary Information.

## Results

### Identification of differentially expressed genes and their expression patterns

Overall, our DE analysis revealed 1,151 transcripts that were differentially expressed between invasion phases. These consisted of 131 transcripts with unique regulation in toads from the range core, 904 transcripts with unique regulation in toads from intermediate areas, and 122 transcripts with unique regulation in toads from the invasion front (list of transcripts in each phase in Appendix I). Soft clustering analysis revealed six prominent expression patterns (clusters) of differentially expressed genes across the invasion (Figure 2): these clusters correspond to up-and down-regulation of each invasion phase (core, intermediate, front). Only 340 of the 1,151 differentially expressed transcripts had sufficiently high membership values to fit within these six clusters (list of transcripts in each cluster in Appendix I). The first cluster depicts low expression at the core and equally high expression throughout the rest of the range. The other five clusters all depict curvilinear patterns (in which expression is either highest or lowest in toads from intermediate areas), but vary in the expression levels in toads on the ends of the range. Gene ontology enrichment of each cluster from Figure 2 is shown in Figure 3, and functional characterization of each cluster based on individual transcript investigation, are shown in Table 1.

**Figure 2.**
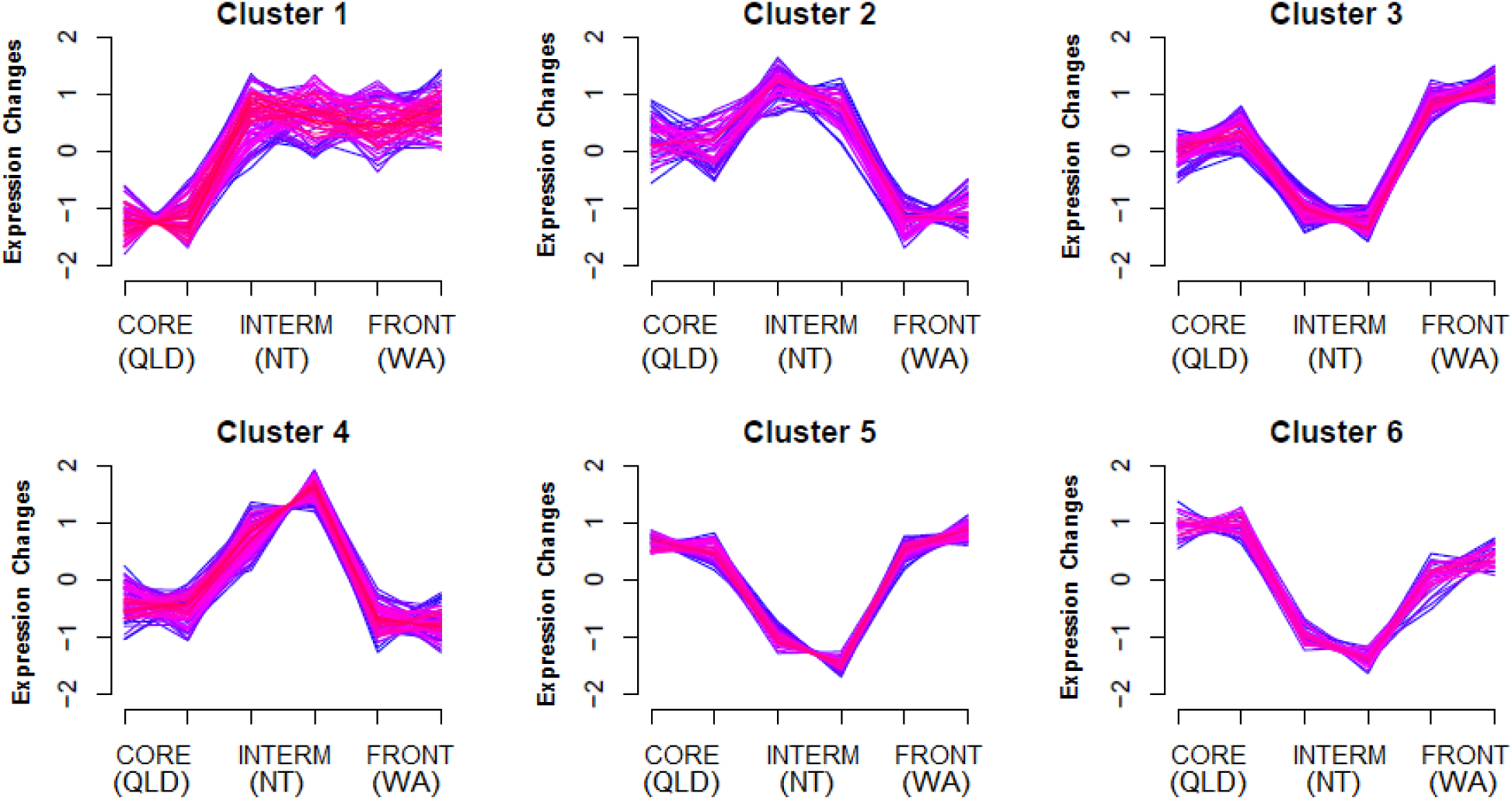
Six unique patterns of gene expression in spleen tissue from invasive cane toads (*Rhinella marina*). We collected samples in populations from areas spanning the invaded range in Australia (QLD = Queensland, NT = Northern Territory, WA = Western Australia). Color indicates membership values of genes to clusters (purple = 0.7-0.8; pink = 0.8-0.9; red = 0.9-1). Tick marks on the x-axis indicate sites across the toad’s Australian range in which spleens were collected (Figure 1).

**Figure 3.**
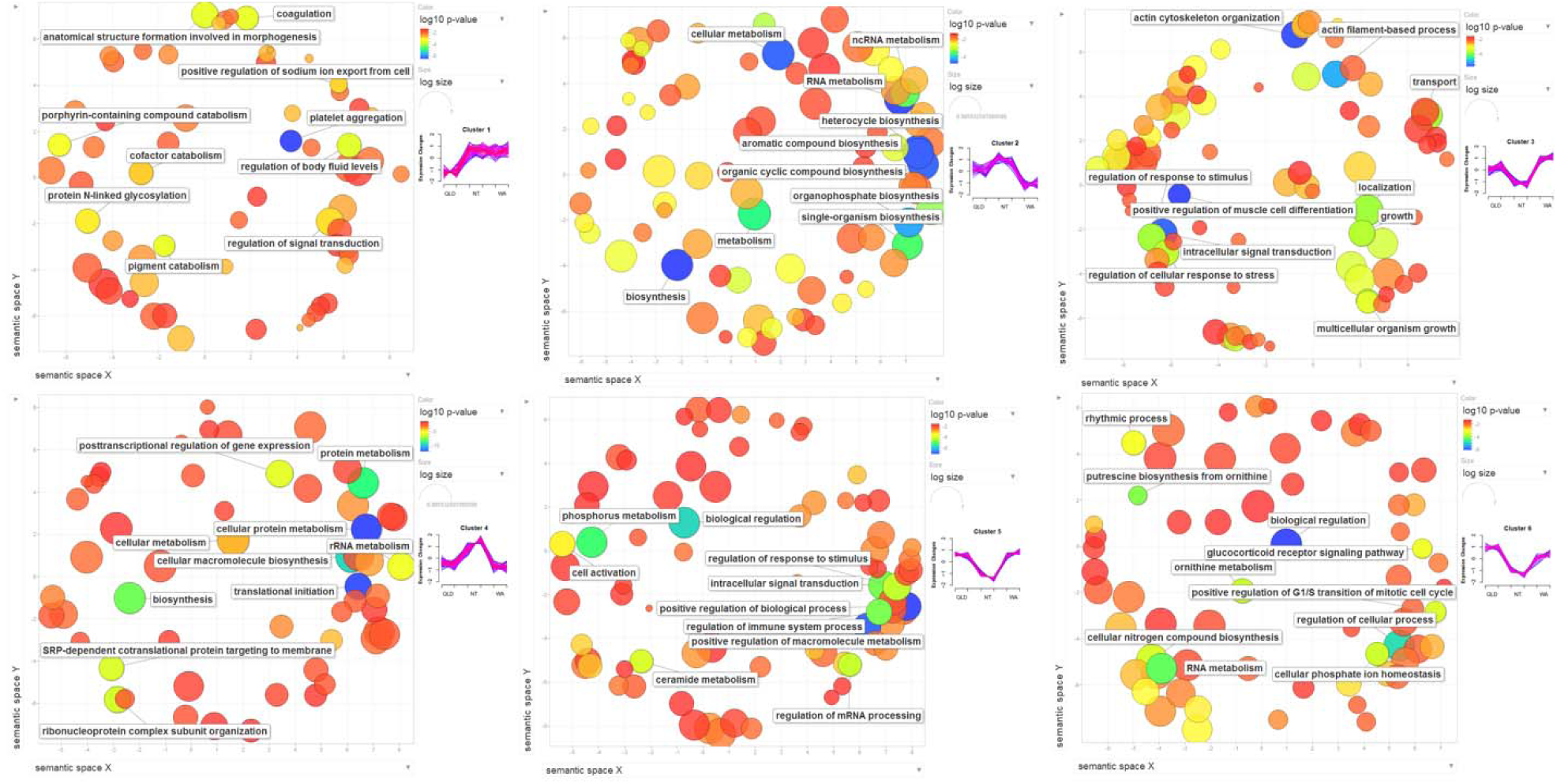
REVIGO plots displaying gene ontology (GO) terms (concepts/classes used to characterize gene function) depicting biological processes associated with transcripts following six major expression patterns in cane toads (*Rhinella marina*) across their Australian range (Figure 1). RNA-Seq data from spleens was used to identify differentially expressed transcripts between invasion phases, then soft clustering was performed to visualize the expression patterns that these transcripts follow (Figure 2). Circles represent GO terms; those with the highest statistical significance are labelled. Circle size relates to breadth of GO terms. Colors show log_10_ p-values.

### Immune genes

The most common immune functions that we found among our differentially expressed transcripts were activation (list of transcripts and their expression patterns/clusters in Table 2a) and suppression (Table 2b) of inflammatory pathways, and cytotoxicity (Table 2c). Immune genes with other roles were too few to allow clear inferences to be drawn. Most pro-inflammatory transcripts were down-regulated in toads from intermediate areas relative to toads from the range core and invasion front. This trend is reflected in the spatial expression data; in the fifth cluster (low expression in intermediate areas, high expression on either end of the range), approximately one quarter of transcripts were identified as relating to immune function (the highest proportion of any cluster), and this is the only cluster in which a GO term directly related to immunity was among the most significant (Figure 3, Table 1). Furthermore, additional immune transcripts were seen in the third and sixth clusters (which also exhibit lowest expression in toads from intermediate areas). Conversely, in the fourth cluster (high expression in intermediate areas, low expression on either end of the range), no immune transcripts were identified and all the most significant GO terms are related to translation (Figure 3, Table 1). Nonetheless, a few pro- and anti-inflammatory transcripts were up-regulated in intermediate toads (Table 2).

### Climate-influenced gene expression

Our LFMM revealed eleven transcripts with expression levels associated with maximum temperature during the hottest month, rainfall during the driest quarter, or both (list of transcripts in Appendix II). Three of these transcripts followed the expression pattern of the first cluster (low expression at the range core, high expression throughout the rest of the range): two are involved in cell adhesion and platelet activity, and function of the third is unknown. Two other transcripts were also down-regulated at the core (but not a member of the first cluster); these are involved in transcription regulation and metabolism. Conversely, two transcripts involved in inflammation activation were up-regulated at the core. Another transcript, involved in cell cycle regulation, was up-regulated in intermediate areas. The three remaining transcripts were not differentially expressed: the first is involved in cell signaling in response to damage, the second is involved in blood circulation and response to nitric oxide (NO), and the third is unknown.

### Coordination in gene expression

We tested whether some transcripts were co-associated across invasion phases by examining the expected value of ρ metric (Quinn et al., 2017a), but only found large groups of co-associated transcripts involved in fundamental cellular processes such as translation (Figure S3; list of proportional transcripts in Appendix III).

### Isolation by distance

Our Mantel test revealed a significant relationship between geographic distance and gene expression distance (*p* = 0.005, R^2^ = 0.02; Figure 4).

**Figure 4.**
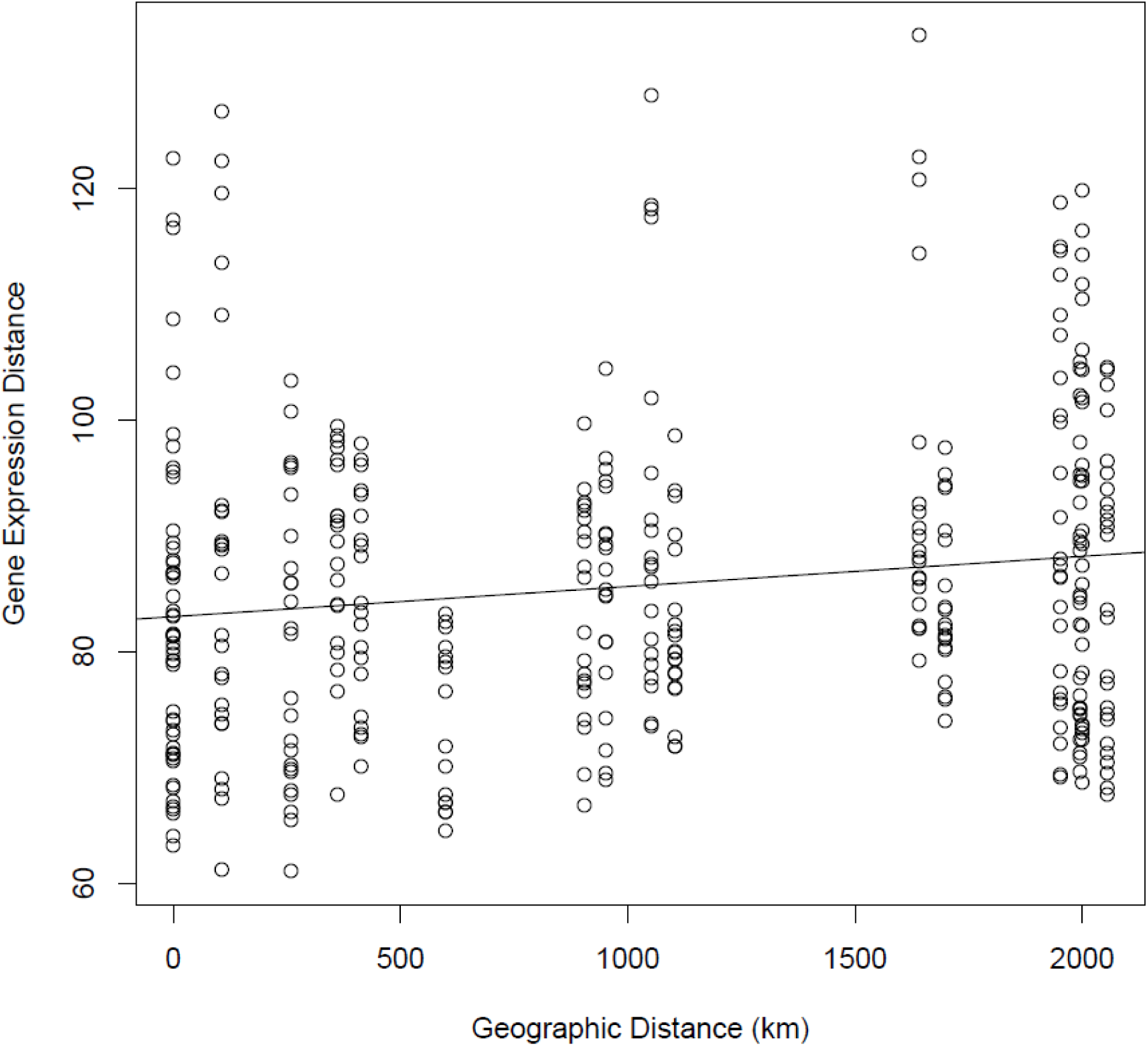
Correlation between geographic distance and gene expression distance of invasive cane toad (*Rhinella marina*) populations across their Australian range (Figure 1). Euclidean distances in geographic location and gene expression between populations were calculated using the dist function in R. A mantel test (performed with the ade4 package) confirmed that these were significantly correlated (*p* = 0.005).

## Discussion

The conceptual scheme of Lee and Klasing (2004), based on the ERH, predicts that the expression of genes encoding mediators of costly immune responses (e.g. inflammation) would follow a density-driven curvilinear pattern, with highest expression in toads from intermediate areas (where density is presumably highest, facilitating parasite transmission). However, most immune transcripts (involved pro- or anti-inflammatory signaling) in our study followed the opposite pattern: curvilinear, but with lowest expression in toads from intermediate areas.

Although not all immune transcripts followed this pattern, our data largely do not support the enemy release hypothesis. It must be noted that our inferences are based on gene annotations sourced from many different taxa. These genes may not function the same way in cane toads as they do in the organisms from which they were described. Furthermore, although we only sampled toads that appeared to be healthy, the infection status of each toad was not manipulated nor controlled for. Thus, spatial heterogeneity in pathogen pressure or environmental conditions may play a role in gene expression; however, parasite prevalence is generally higher in intermediate-age populations than in younger and older populations (Freeland et al., 1986), inconsistent with the general pattern for down-regulation of immune transcripts in toads from intermediate-age populations. Our results may be driven by other environmental variables, or may have a genetic basis through polymorphism in promoter regions, which are not sequenced when using RNA-Seq (Wang et al., 2009).

Our findings are consistent with previous phenotypic data on toads: spleen sizes and fat body masses also follow a curvilinear pattern in which they are lowest in toads from intermediate parts of the range (Brown et al., 2015b). If toads indeed follow a ‘traveling wave’ density pattern, then the increased densities that occur several years-post colonization would not just heighten pathogen-mediated selection, but also intraspecific competition; reduced access to food resources may thus constrain the toad’s ability to increase investment in immunity (Brown et al., 2015b). This may explain smaller spleen sizes and lower expression of transcripts involved in inflammatory immune responses in toads from intermediate areas. Availability of energetic resources was not considered in the predictions of Lee & Klasing (2004), and although individuals at an invasion front may have a reduced benefit from energetic investment into immunity, they may invest anyway due to reduced intraspecific competition for food. Thus, the strength of immune functioning in invasion front individuals may depend on factors other than enemy release, such as prey abundance and interspecific competition.

Enemy release is not the only potential selective force acting on cane toads as they disperse across Australia; we also searched for evidence of adaptation to aridity. The range core is cooler and receives more rainfall than do intermediate areas and the invasion front. The expression pattern depicted in our first cluster (low expression in the core, high throughout the rest of the range) matches the trend across the Australian range in temperature, and is opposite to the trend in rainfall. Population genetic structure in toads appears to be driven by these environmental variables (Selechnik et al., 2019). Three of the eleven transcripts with expression levels associated with temperature or rainfall belonged to this first cluster; the rest were not members of any of the six clusters. Two out of these three are involved in blood clotting, as are most of the transcripts in the first cluster. Proportionality analysis revealed two small groups of clotting-related transcripts with coordinated expression, suggesting coordinated function. Blood clotting is affected by hydration levels (El-Sabban et al., 1996), and excessive blood clotting can impair health (PubmedHealth, 2014). Changes to the rate of blood clotting may reflect hydric-related adaptations of intermediate and frontal toads living in drier conditions. Furthermore, four candidate SNP loci (with outlier F_ST_ values and associations with rainfall, indicating they may be under selection) have previously been found in a gene involved in blood clotting (Selechnik et al., 2019). However, if lowering blood clotting rates is indeed an adaptation to aridity in toads from intermediate areas and the front, then one might expect that transcripts promoting blood clotting would be down-regulated in these individuals (yet in our study, they are up-regulated). This may be because toads were collected from the wild and the environment was not controlled for. A common-garden experiment with toads from across the range may reveal whether modification of blooding clotting rates is an evolved adaptation to aridity or simply a physiological reaction to different environments. Nonetheless, the differences in expression of genes underlying these traits supports the hypothesis that cane toads are adapting to their abiotic environment.

Our results suggest that other evolutionary forces are at work as well. The curvilinearity in expression of many of our differentially expressed genes resembles that of other phenotypic traits affected by spatial sorting, such as leg length (Hudson et al., 2016). Physical activity can have transgenerational effects on gene expression (Barres & Zierath, 2016; Murashov et al., 2016), so dispersal may directly affect expression patterns. Additionally, the significant result of our Mantel test is likely due to either genetic drift or a balance between geographically varying selection and gene flow (Endler, 1977). Although this result may be influenced by environmental gradients, it is unlikely that selection driven by environmental factors would act on a genome-wide level (all expressed transcripts were used in our calculations for gene expression distance). This result suggests that genetic drift may be responsible for isolation by distance in Australian cane toads. Because spatial sorting and genetic drift drive non-adaptive variation, their effects may obscure adaptive variation, particularly when they act on the same traits as selection (i.e. physical activity is also linked to expression of inflammatory genes (Baynard et al., 2012; Gjevestad et al., 2015)).

## Conclusions

The expression patterns that we observed in pro- and anti-inflammatory transcripts generally do not support the predictions of Lee & Klasing (2004) based on the ERH. Notably, our study suggests that the predicted consequences of enemy release are difficult to study in the wild because host-parasite dynamics and their impacts are affected by many factors. Increasing host densities bolsters both parasite transmission and host intraspecific competition, but these two factors exert opposite effects on host immune function: while high rates of parasite transmission favour powerful immune responses, high levels of intraspecific competition limit the energetic resources available to fuel these immune responses. Common-garden studies may clarify this situation. Although natural selection causes adaptive change in immune function and other traits, this co-occurs with non-adaptive variation driven by forces such as spatial sorting and genetic drift, particularly in invasive species. These complicated dynamics may explain why support for the predictions of the ERH has been so mixed over the past two decades. Nonetheless, methods such as RNA-Seq remain a powerful tool for uncovering the diverse and sometimes opposing forces that underpin rapid evolution in invasive species.

## Acknowledgements

This work was supported by the Australian Research Council (FL120100074, DE150101393) and the Equity Trustees Charitable Foundation (Holsworth Wildlife Research Endowment). We thank BriAnne Addison and Thom Quinn for their input during discussions about our analyses. We thank John Endler and Lynn B. Martin for their useful comments that improved our manuscript. We thank Cam Hudson, Serena Lam, and Chris Jolly for their assistance with sample collection.

## Data Accessibility

The dataset supporting the conclusions of this article is available in the NCBI short read archive (SRA) under the BioProject Accession PRJNA395127 in FASTQ format (https://www.ncbi.nlm.nih.gov/bioproject/PRJNA395127/).

## Authors’ contributions

Dan Selechnik, Mark Richardson, Lee Ann Rollins, Greg Brown, and Richard Shine designed the experiment. Dan Selechnik, Mark Richardson, and Lee Ann Rollins performed data collection. Dan Selechnik, Mark Richardson, and Lee Ann Rollins performed data analysis. Dan Selechnik wrote the manuscript. Mark Richardson, Lee Ann Rollins, Greg Brown, and Richard Shine revised the manuscript.

