## Supplementary material for "Immune and environment-driven gene expression during invasion: An eco-immunological application of RNA-Seq": Table 1

Table 1. Most common functions of transcripts in each of the six major expression patterns in cane toads (*Rhinella marina*) across their Australian range (Figure 1). RNA-Seq data from spleens was used to identify differentially expressed transcripts between invasion phases, then soft clustering was performed to visualize the expression patterns that these transcripts follow (Figure 2).

| Cluster | Number of transcripts | Most common biological function | Most significantly enriched GO term(s) | Less common biological functions | |
| --- | --- | --- | --- | --- | --- |
| 1 | 71 | blood coagulation/circulation (24 transcripts) | platelet aggregation | signal transduction, immune function, transcription regulation, viral processes | |
| 2 | 49 | none | cellular metabolism,  biosynthesis | metabolism, biosynthesis,  cell cycle regulation, protein ubiquitination,  translation initiation, transcription regulation | |
| 3 | 66 | signal transduction  (24 transcripts) | intracellular signal transduction, actin cytoskeleton organization | protein transport, immune function, smooth muscle contraction,  angiogenesis, cell cycle regulation | |
| 4 | 75 | translation initiation  (28 transcripts) | translation initiation |  | metabolism, transcription regulation,  protein ubiquitination, cell cycle regulation |
| 5 | 49 | immune function  (12 transcripts) | regulation of biological process |  | transcription regulation,  signal transduction, cell cycle regulation,  metabolism |
| 6 | 30 | none | biological regulation |  | transcription regulation,  signal transduction, cell cycle regulation, immune function |
