## Supplementary material for "Immune and environment-driven gene expression during invasion: An eco-immunological application of RNA-Seq": Table 2

Table 2. Genes involved in immune function that are differentially expressed across the range of Australian cane toads. Spleens were collected from toads in the range core (QLD: Gordonvale and Daintree, N=5 each), intermediate areas (NT: Cape Crawford and Timber Creek, N=4 each) and invasion front (WA: Caroline Pool and Durack River, N=5 each). Soft clustering was performed to visualize differential expression patterns between different phases of the invasion (Figure 2).

|  | ***(a) Inflammation*** |  |
| --- | --- | --- |
| Gene | Protein | Expression Pattern |
| *MAP3K2* | Mitogen-activated protein kinase kinase kinase 2 | NT down; Cluster 5 |
| *pik3r5* | Phosphoinositide 3-kinase regulatory subunit 5 | NT down; Cluster 5 |
| *CSF2RB* | Cytokine receptor common subunit beta | NT down; Cluster 5 |
| *srf* | Serum response factor | NT down; Cluster 5 |
| *PTK2B* | Protein-tyrosine kinase 2-beta | NT down; Cluster 5 |
| *PYCARD* (isoform 1) | Apoptosis-associated speck-like protein containing a CARD (ASC) | NT down; Cluster 5 |
| *PYCARD* (isoform 2) | Apoptosis-associated speck-like protein containing a CARD (ASC) | NT down; Cluster 6 |
| *Nlrp1b* | NACHT; LRR and PYD domains-containing protein 1b allele 3 (NLRP1b) | NT down; Cluster 6 |
| *ANKRD17* | Ankyrin repeat domain-containing protein 17 | NT down; Cluster 6 |
| *mapk8* | Mitogen-activated protein kinase 8 | NT down; Cluster 6 |
| *SPAG9* | C-Jun-amino-terminal kinase-interacting protein 4 | NT down; Cluster 3 |
| *Tab1* | TGF-beta-activated kinase 1 and MAP3K7-binding protein 1 | NT down; Cluster 3 |
| *CAMK2G* | Calcium/calmodulin-dependent protein kinase type II subunit gamma | NT down; Cluster 3 |
| *TNFAIP2* | Tumor necrosis factor alpha-induced protein 2 | WA up; NT down; Cluster 3 |
| *PYCARD* (isoform 3) | Apoptosis-associated speck-like protein containing a CARD (ASC) | NT down |
| *P2RX7* | P2X purinoceptor 7 (P2P7) | NT down |
| *NFATC2* | Nuclear factor of activated T-cells; cytoplasmic 2 | NT down |
| *Rps6ka3* | Ribosomal protein S6 kinase alpha-3 | NT down |
| *PIK3CB* | Phosphatidylinositol 4;5-bisphosphate 3-kinase catalytic subunit beta isoform | NT down |
| *NOS2* | Nitric oxide synthase; inducible | NT down |
| *MAP4K5* | Mitogen-activated protein kinase kinase kinase kinase 5 | NT down |
| *HIPK1* | Homeodomain-interacting protein kinase 1 (HIP1) | NT down |
| *STAT1* | Signal transducer and activator of transcription 1 (STAT1) | NT down |
| *prkcb* | Protein kinase C beta type | NT down |
| *RIPK3* | Receptor-interacting serine/threonine-protein kinase 3 (RIPK3) | NT down |
| *Pak2* | Serine/threonine-protein kinase PAK 2 | NT down |
| *TRPC4AP* | Short transient receptor potential channel 4-associated protein | NT down |
| *Nsmaf* | Protein FAN | NT down |
| *PLEKHG5* | Pleckstrin homology domain-containing family G member 5 | NT down |
| *IL7R* | Interleukin-7 receptor subunit alpha | NT down |
| *Tnfrsf21* | Tumor necrosis factor receptor superfamily member 21 | NT down |
| *Rxra* | Retinoic acid receptor RXR-alpha | NT down |
| *TRAF2* | TNF receptor-associated factor 2 | NT down |
| *IKBKE* | Inhibitor of nuclear factor kappa-B kinase subunit epsilon | NT down |
| *Map3k14* | Mitogen-activated protein kinase kinase kinase 14 | NT down |
| *MTOR* | Serine/threonine-protein kinase mTOR | NT down |
| *gtpbp1* | GTP-binding protein 1 | NT down |
| *ADAMTS1* | A disintegrin and metalloproteinase with thrombospondin motifs 1 | NT down |
| *Erc1* | ELKS/Rab6-interacting/CAST family member 1 | NT down |
| *ARHGEF17* | Rho guanine nucleotide exchange factor 17 | NT down; WA up |
| *Dele* | Death ligand signal enhancer | WA up; Cluster 3 |
| *Cd84* | SLAM family member 5 | WA up |
| *F2RL2* | Proteinase-activated receptor 3 | WA up |
| *TNFRSF19* | Tumor necrosis factor receptor superfamily member 19 | WA up |
| *IL4R* | Interleukin-4 receptor subunit alpha | WA up |
| *mapk1* | Mitogen-activated protein kinase 1 | WA up |
| *Irgc* (isoform 1) | Interferon-inducible GTPase 5 | QLD up |
| *Irgc* (isoform 2) | Interferon-inducible GTPase 5 | QLD up |
| *Irgc* (isoform 3) | Interferon-inducible GTPase 5 | QLD up |
| *Irgc* (isoform 4) | Interferon-inducible GTPase 5 | QLD up |
| *Irgc* (isoform 5) | Interferon-inducible GTPase 5 | QLD up |
| *mul1a* | Mitochondrial ubiquitin ligase activator of nfkb 1-A | QLD down; Cluster 1 |
| *CSF2RA* | Granulocyte-macrophage colony-stimulating factor receptor subunit alpha | QLD down; Cluster 1 |
| *TRIM25* | E3 ubiquitin/ISG15 ligase TRIM25 | QLD down |
| *PYCARD* (isoform 4) | Apoptosis-associated speck-like protein containing a CARD (ASC) | NT up |
| *ecsit* | Evolutionarily conserved signaling intermediate in Toll pathway; mitochondrial | NT up |
| *NKAP* | NF-kappa-B-activating protein | NT up |
|  | ***(b) Anti-Inflammation*** |  |
| Gene | Protein | Expression Pattern |
| *Tank* | TRAF family member-associated NF-kappa-B activator | NT down; Cluster 5 |
| *ERBIN* | Erbin | NT down; Cluster 5 |
| *Itch* | E3 ubiquitin-protein ligase Itchy | NT down; Cluster 5 |
| *Sbno1* | Protein strawberry notch homolog 1 | NT down; Cluster 5 |
| *Smad6* | Mothers against decapentaplegic homolog 6 | NT down; Cluster 3 |
| *inpp5d* | Phosphatidylinositol 3,4,5-trisphosphate 5-phosphatase 1 | NT down |
| *SBNO2* | Protein strawberry notch homolog 2 | NT down |
| *SYNCRIP* | Heterogeneous nuclear ribonucleoprotein Q | NT down |
| *ATF3* | Cyclic AMP-dependent transcription factor ATF-3 | NT down |
| *Dusp4* | Dual specificity protein phosphatase 4 | NT down |
| *Rps6ka4* | Ribosomal protein S6 kinase alpha-4 | NT down |
| *AHR* | Aryl hydrocarbon receptor | NT down |
| *PTPRE* | Receptor-type tyrosine-protein phosphatase epsilon | NT down |
| *HAX1* | HCLS1-associated protein X-1 | NT up |
| *ppp4c* | Serine/threonine-protein phosphatase 4 catalytic subunit | NT up |
| *Nlrc3* (isoform 1) | Protein NLRC3 | NT up |
| *Nlrc3* (isoform 2) | Protein NLRC3 | NT up |
| *ANXA1* | Annexin A1 | NT up |
| *ciapin1* | Anamorsin | NT up |
| *impdh2* | Inosine-5'-monophosphate dehydrogenase 2 | NT up |
| *DHCR24* | Delta(24)-sterol reductase | NT up |
| *CD200R1B* | Cell surface glycoprotein CD200 receptor 1-B | QLD down; Cluster 1 |
| *TNFRSF6B* | Tumor necrosis factor receptor superfamily member 6B | QLD down |
|  | ***(c) Cytotoxicity*** |  |
| Gene | Protein | Expression Pattern |
| *STXBP2* | Syntaxin-binding protein 2 | NT down; Cluster 5 |
| *NCR3LG1* (isoform 1) | Natural cytotoxicity triggering receptor 3 ligand 1 | NT down; Cluster 5 |
| *NCR3LG1* (isoform 2) | Natural cytotoxicity triggering receptor 3 ligand 1 | NT down |
| *NCR3LG1* (isoform 3) | Natural cytotoxicity triggering receptor 3 ligand 1 | NT down |
| *NCR3LG1* (isoform 4) | Natural cytotoxicity triggering receptor 3 ligand 1 | NT down |
| *Slamf7* | SLAM family member 7 | NT down; WA up |
