## Supplementary material for "Immune and environment-driven gene expression during invasion: An eco-immunological application of RNA-Seq": Table S1

**Table of Contents:**

| **Supp. Methods** | Page 1 |
| --- | --- |
| **Supp. Results & Discussion** | Page 2 |
| **Supp. Tables & Figures** | Page 3 |
| **References** | Page 7 |

**Supplemental Methods**

*Available annotation information*

In the reference transcriptome (Richardson et al., 2018) that we used for performing alignments, approximately 30% of all transcripts are functionally characterized, and approximately 4% of all transcripts are characterized in *Xenopus* (the phylogenetically closest model taxa to *Rhinella marina*). In our list of differentially expressed genes, approximately 85% of all transcripts are functionally characterized, and approximately 17% of all transcripts are characterized in *Xenopus*.

*Coordination in gene expression*

Correlation in expression patterns of different transcripts is of interest because it may suggest that the expression of one transcript is coordinated with the expression of another, which may extend to functional relations between the two; however, correlation is not a statistically valid measure for compositional data such as ours (Quinn et al., 2017a). Proportionality (ρ) is an alternative measure of coordination that is statistically valid for compositional data (Quinn et al., 2017a); it is a modification of the variance of the log ratios (VLR) that sets a scale on which transcript expression can be compared to look for coordination (Quinn et al., 2017a). Similarly to the correlation coefficient *r*, ρ exists on a scale from -1 to 1; in this way, they are analogous (Quinn et al., 2017a). To see if there was coordination among different transcripts in expression, we tested our transcripts for proportionality using propr v2.1.9 (Quinn et al., 2017a). We used the iqlr-transformed counts produced by ALDEx2 as expression values. Proportionality was calculated as:

$\rho\left( A_{i},A_{j} \right)=1- \frac{var(A_{i}-A_{j})}{var\left( A_{i} \right)+var(A_{j})}$	(1)

In this equation (Erb & Notredame, 2016; Quinn et al., 2017a), A_i_ and A_j_ are the log-ratio transformed vectors of two different transcripts (Quinn et al., 2017a). Pairs of transcripts with ρ > 0.90 or ρ < -0.90 were classified as proportional; we chose the ρ cutoff at 0.90 based on our sample size (28) and number of transcripts (17597), as suggested by Quinn *et al* (2017). We chose to investigate all groups of proportional transcripts that contained at least one differentially expressed transcript, as well as any group containing more than 50 transcripts (N > 50).

*Isolation by distance*

To examine the effects of geographic distance on genetic distance, we performed a Mantel test using ade4 v1.7-5 (Thioulouse & Dray, 2007). If gene expression differences are primarily driven by changes in allele proportions due to drift or selection driven by environmental gradients co-occurring with the expanding range, then we might expect a significant linear relationship between geographic distance and genetic distance. However, if gene expression differences are primarily driven by heterogeneous environments at sampling sites, then we would expect a non-significant relationship between geographic distance and genetic distance. We used the dist function in R (Team, 2016) to calculate the Euclidean distances in geographic space between all samples using the coordinates of their collection sites. We then used the dist function to calculate the Euclidean distances in gene expression using the iqlr-transformed counts of all transcripts, thus generating a genetic distance matrix. Genetic distance was calculated as:

$d=\sqrt{\sum_{1}^{N} {(x_{i}-x_{j})}^{2}}$	(3)

In this equation (Team, 2016), *x_i_* is the log-transformed count value of transcript *x* in sample *i*, and *x_j_* is the log-transformed count value of the same transcript in sample *j*. The expression difference in each of N transcripts is calculated, and then the square root of the sum of squares of all expression differences is computed.

**Supplemental Results & Discussion**

*Coordination in gene expression*

We identified many groups of proportionally expressed (co-associated) transcripts; however, few of these groups contained transcripts that were also differentially expressed. We focused only on those that were (list of proportional transcripts in Appendix III). The largest group of proportionally expressed transcripts (N=124; Figure S3A), some of which were part of the fourth cluster (high expression in intermediate areas, equally low expression at the ends of the range) was involved in translation, with functions such as ribosome binding, initiation, elongation, termination, and fidelity. We also identified two small groups of proportionally expressed transcripts mostly involved in platelet activation and adhesion, most of which were part of the first cluster (low expression at the core, equally high expression throughout the rest of the range).

Although we found few immune transcripts up-regulated in toads from intermediate areas (and none in our fourth cluster), this cluster does consist of many transcripts with similar functions: approximately half of the transcripts are involved in translation initiation, and proportionality analysis revealed that many of them are coordinated. Because translation serves a wide variety of roles, it is surprising to see a large group of transcripts encoding ribosome components and translation initiation factors up-regulated in intermediate toads. One explanation for this may be that transcripts up-regulated by intermediate toads are, on average, shorter than transcripts up-regulated by toads from the other invasion phases (2555 bases in core toads, 1012 bases in intermediate toads, 1824 bases in frontal toads). Shorter transcripts yield higher ribosome density, and the rate of translation initiation may be higher when transcripts are shorter (Ingolia et al., 2009). Ribosomes are likely able to begin a new round of translation more frequently if they are translating shorter transcripts. Intermediate toads up-regulate the shortest transcripts of any phase, and down-regulate the longest transcripts; there is no clear adaptive explanation for this trend.

**Supplemental Tables & Figures**

Table S1. ERCC spike-in mixes added to RNA from spleens collected from cane toads (*Rhinella marina*) across the Australian range (main text, Figure 1), RNA integrity numbers (RINs), sexes, accession numbers, RNA concentrations, numbers of reads, and numbers of transcripts retained after quality control filtering (out of 18945 transcripts total). RNA-Seq data from spleens was used to quantify differential expression (DE) analysis between localities (states). ERCC spike-ins are used to assess technical performance of sequencing. RINs indicate the quality of the extracted RNA.

| Spleen ID | Location | Spike-In | RIN | Sex | SRA Accession | RNA conc (ng/μL) | Reads | Transcripts retained after QC |
| --- | --- | --- | --- | --- | --- | --- | --- | --- |
| S1 | Gordonvale, QLD | 1 | 8.6 | F | SRX3030411 | 84.01 | 23784416 | 18810 |
| S2 | Gordonvale, QLD | 1 | 8.7 | F | SRX3030410 | 44.78 | 24958948 | 18839 |
| S3 | Gordonvale, QLD | 2 | 7.4 | F | SRX3030409 | 71.90 | 24238536 | 18786 |
| S4 | Gordonvale, QLD | 2 | 7.1 | F | SRX3030408 | 65.34 | 25172372 | 18780 |
| S5 | Gordonvale, QLD | 1 | 7.4 | F | SRX3030397 | 41.98 | 23863836 | 18809 |
| S6 | Daintree, QLD | 2 | 7.4 | F | SRX3030404 | 80.98 | 25504510 | 18825 |
| S7 | Daintree, QLD | 1 | 9.5 | F | SRX3030403 | 56.05 | 23452460 | 18679 |
| S8 | Daintree, QLD | 1 | 7.3 | F | SRX3030405 | 40.72 | 24403770 | 18781 |
| S9 | Daintree, QLD | 1 | 9.4 | F | SRX3030406 | 71.26 | 21885152 | 18811 |
| S10 | Daintree, QLD | 2 | 8.9 | F | SRX3030401 | 56.55 | 22693350 | 18805 |
| S11 | Halls Creek, WA | 2 | 8.9 | F | SRX3030396 | 70.10 | 22491142 | 18851 |
| S12 | Halls Creek, WA | 2 | 9.2 | F | SRX3030395 | 67.09 | 24040758 | 18779 |
| S13 | Halls Creek, WA | 2 | 7.0 | F | SRX3030394 | 65.81 | 23746642 | 18835 |
| S14 | Halls Creek, WA | 1 | 9.2 | F | SRX3030393 | 77.76 | 24426716 | 18852 |
| S15 | Halls Creek, WA | 1 | 9.7 | F | SRX3030392 | 78.91 | 23278544 | 18709 |
| S16 | Durack River, WA | 2 | 8.6 | F | SRX3030385 | 172.58 | 24374724 | 18803 |
| S17 | Durack River, WA | 1 | 8.5 | F | SRX3030389 | 99.38 | 23344088 | 18828 |
| S18 | Durack River, WA | 2 | 8.8 | F | SRX3030407 | 41.42 | 24654968 | 18740 |
| S19 | Durack River, WA | 1 | 9.0 | F | SRX3030384 | 77.47 | 23608774 | 18840 |
| S20 | Durack River, WA | 2 | 8.5 | F | SRX3030402 | 61.83 | 25004504 | 18834 |
| S21 | Cape Crawford, NT | 1 | 8.2 | F | SRX3030387 | 103.51 | 25116754 | 18835 |
| S22 | Cape Crawford, NT | 2 | 8.1 | F | SRX3030388 | 34.76 | 19923840 | 18761 |
| S23 | Cape Crawford, NT | 1 | 9.6 | F | SRX3030390 | 92.68 | 26615752 | 18791 |
| S24 | Cape Crawford, NT | 2 | 9.3 | F | SRX3030391 | 83.97 | 27652796 | 18773 |
| S25 | Timber Creek, NT | 2 | 8.7 | F | SRX3030386 | 43.59 | 19004402 | 18758 |
| S26 | Timber Creek, NT | 2 | 9.2 | F | SRX3030400 | 102.68 | 27999430 | 18820 |
| S27 | Timber Creek, NT | 1 | 9.0 | F | SRX3030399 | 73.58 | 29437400 | 19794 |
| S28 | Timber Creek, NT | 1 | 7.1 | F | SRX3030398 | 69.74 | 23455440 | 18771 |

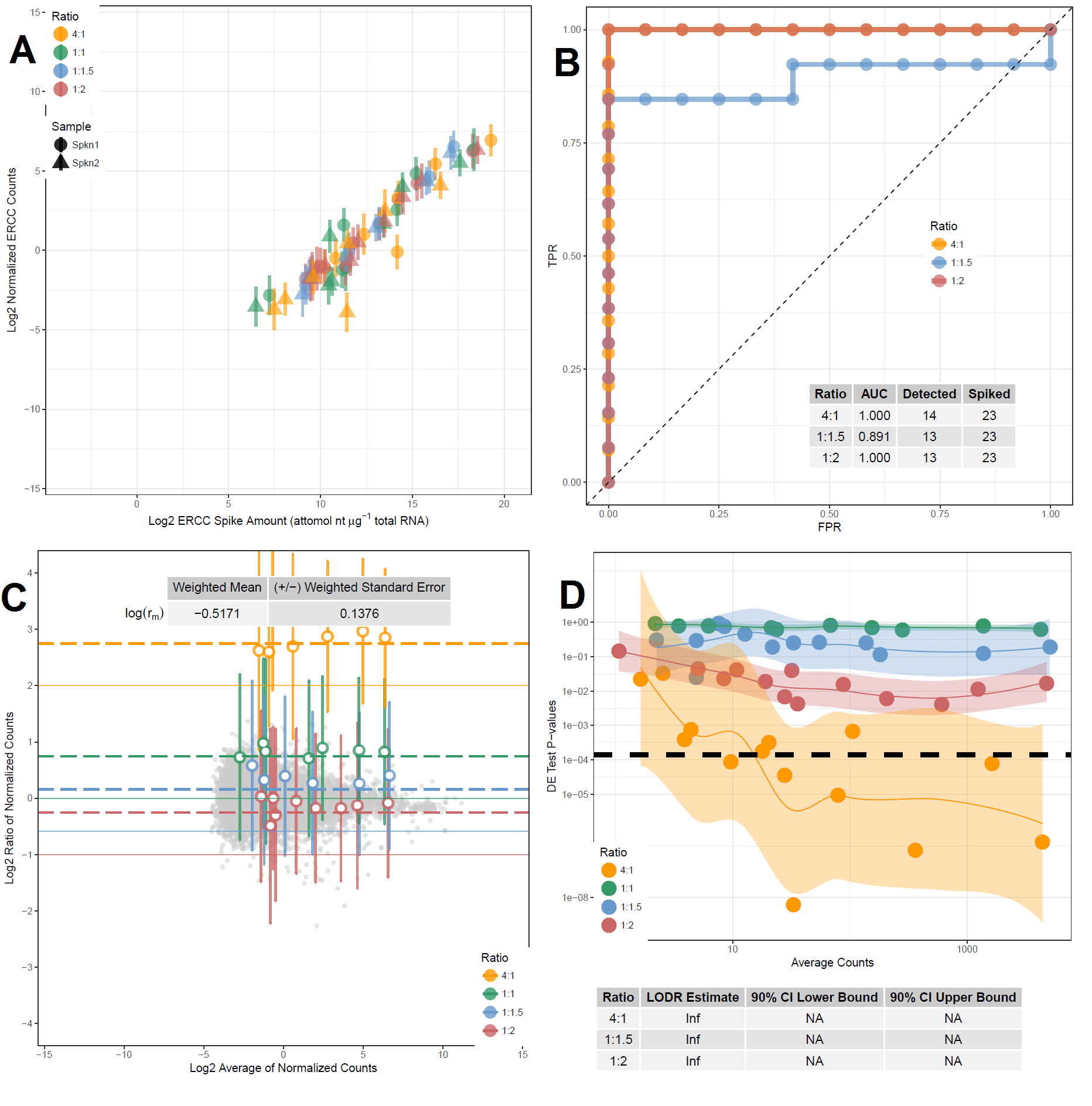

Figure S1. ERCC technical and diagnostic plots produced by erccdashboard. (A) Signal-abundance plot. Colors represent ERCC sequence sets that are present in abundances of different ratios between mix 1 and mix 2, shape represents the sample type, and error bars represent standard deviation. (B) ROC curves and AUC statistics for each set of true-positive ERCC controls. “Detected” represents the number of controls used in our experiment, and “spiked” represents the total that were present in the ERCC control mixture. (C) MA plot of ERCC ratio measurement variability and bias. Colors represent ERCC sequence sets that are present in abundances of different ratios between mix 1 and mix 2, and error bars represent standard deviation. Grey points represent endogenous transcript ratios. Solid colored lines show the nominal ERCC ratios for each ERCC sequence set, and dashed lines (r_m_) represent the corrected ratios for each ERCC subset based on the estimate of mRNA fraction differences between samples. (D) Lower limit of differential expression detection estimates (LODR) plot. The black dashed line represents the threshold p-value for the specified FDR (0.05). Colors represent ERCC sequence sets that are present in abundances of different ratios between mix 1 and mix 2. Colored lines are the LODR estimates, indicating if the p-value of each ratio is ever low enough to cross the threshold p-value and be called DE. LODR results and 90% confidence intervals are provided in the table below the plot.

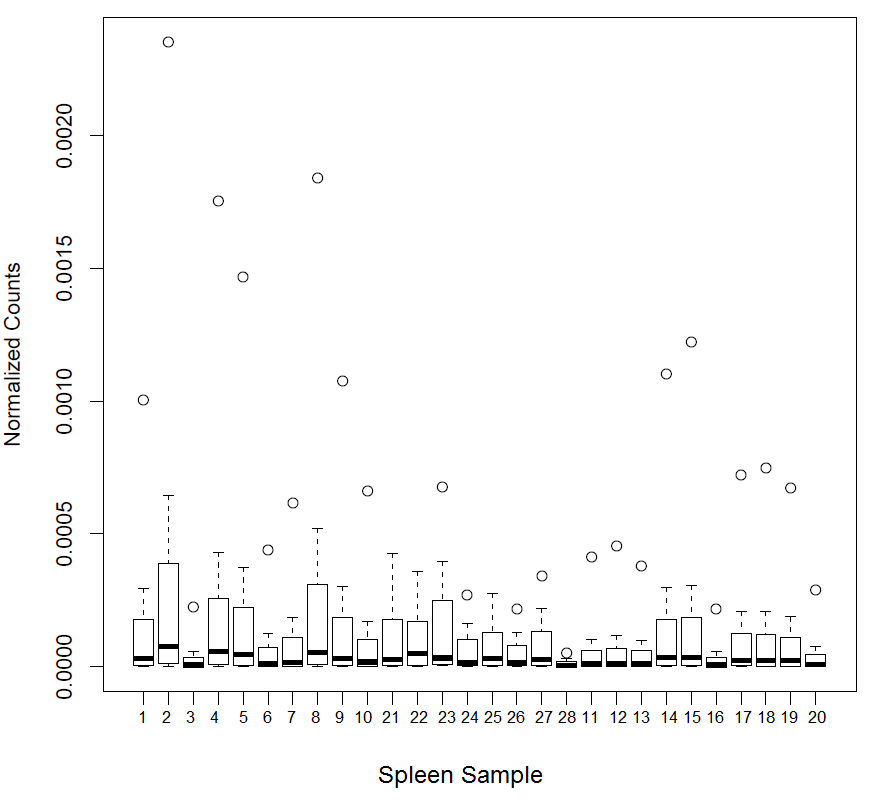

Figure S2. Normalized counts of seven invariant ERCC sequences (mix 1 versus mix 2 fold-change 1:1) across all samples. ERCC spike-in mixes were added to RNA from spleens collected from cane toads (*Rhinella marina*) across their Australian range (main text, Figure 1). Counts of ERCC sequences and endogenous transcripts were generated, then filtered to remove zeroes. Only seven invariant ERCC sequences were retained; count values of these seven ERCC sequences in each sample were divided by the library size of the sample to generate normalized counts.

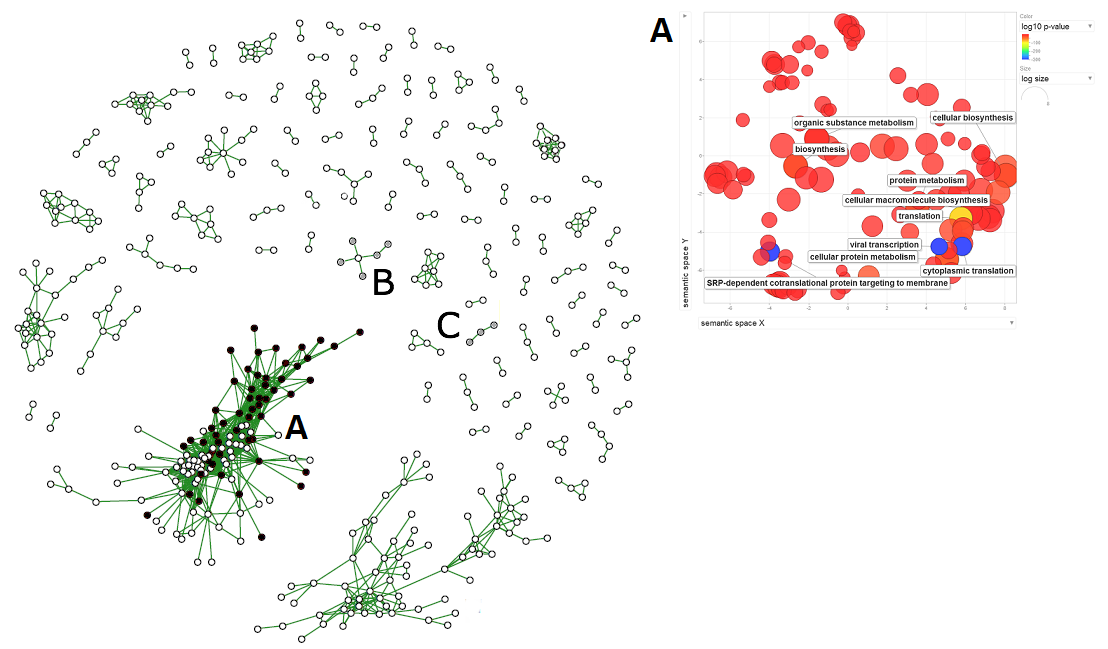

Figure S3. Groups of proportionally expressed transcripts in spleen tissue from the invasive Australian cane toad (*Rhinella marina*). Nodes represent transcripts, and lines indicate proportionality between them. Black fill denotes up-regulation in toads from intermediate areas, gray fill denotes down-regulation in toads from the core, and no fill denotes no differential expression. All transcripts were positively proportional. The propr package in R was used to identify transcripts with expression patterns that were proportional to those of other transcripts from RNA-Seq data obtained from spleens sampled across the Australian range (Figure 1). A REVIGO plot of the largest group of proportionally expressed transcripts is included to show their most common functions.
