## Appendix I for "Immune and environment-driven gene expression during invasion: An eco-immunological application of RNA-Seq"

**Supplemental Information for:**

**How important is enemy release in explaining immune gene expression during invasion? An eco-immunological application of RNA-Seq**

Selechnik D, Richardson MF, Shine R, Brown GP, and Rollins LA

**Table of Contents:**

| **Appendix I (all DE genes)** | Page 2 |
| --- | --- |
| **Core (QLD)** | Page 2 |
| **Core (QLD) Down** | Page 3 |
| **Intermediate (NT) Up** | Page 7 |
| **Intermediate (NT) Down** | Page 21 |
| **Front (WA) Up** | Page 39 |
| **Front (WA) Down** | Page 41 |
| **Cluster 1** | Page 44 |
| **Cluster 2** | Page 47 |
| **Cluster 3** | Page 49 |
| **Cluster 4** | Page 51 |
| **Cluster 5** | Page 54 |
| **Cluster 6** | Page 56 |
| **Appendix III (proportional genes)** | Page 58 |

**Appendix I**

Differentially expressed transcripts in cane toads (*Rhinella marina*) across the Australian range (main text, Figure 1). RNA-Seq data from spleens was used to quantify differential expression (DE) analysis between phases of the invasion.

Table AI1. Core (QLD) up-regulation

| Gene | Protein | effect | *p* |
| --- | --- | --- | --- |
| *Irgc* | Interferon-inducible GTPase 5 | 2.03 | 0.00 |
| *Irgc* | Interferon-inducible GTPase 5 | 1.59 | 0.00 |
| *QRICH1* | Glutamine-rich protein 1 | 1.57 | 0.00 |
| *AIM1L* | Absent in melanoma 1-like protein | 1.35 | 0.01 |
| *stmn1-a* | Stathmin-1-A | 1.34 | 0.01 |
| *.* | . | 1.34 | 0.01 |
| *.* | . | 1.30 | 0.02 |
| *COL4A1* | Collagen alpha-1(IV) chain | 1.24 | 0.02 |
| *MGEA5* | Protein O-GlcNAcase | 1.23 | 0.02 |
| *GRK6* | G protein-coupled receptor kinase 6 | 1.17 | 0.03 |
| *Irgc* | Interferon-inducible GTPase 5 | 1.16 | 0.02 |
| *ezh2* | Histone-lysine N-methyltransferase EZH2 | 1.16 | 0.02 |
| *Irgc* | Interferon-inducible GTPase 5 | 1.15 | 0.02 |
| *AZIN1* | Antizyme inhibitor 1 | 1.14 | 0.03 |
| *.* | . | 1.12 | 0.03 |
| *RBM25* | RNA-binding protein 25 | 1.11 | 0.03 |
| *PTPRJ* | Receptor-type tyrosine-protein phosphatase eta | 1.09 | 0.02 |
| *PRPF4B* | Serine/threonine-protein kinase PRP4 homolog | 1.08 | 0.02 |
| *Xpo1* | Exportin-1 | 1.08 | 0.03 |
| *chmp4b* | Charged multivesicular body protein 4b | 1.08 | 0.02 |
| *DNAH11* | Dynein heavy chain 11, axonemal | 1.08 | 0.04 |
| *GPCPD1* | Glycerophosphocholine phosphodiesterase GPCPD1 | 1.08 | 0.04 |
| *Nav3* | Neuron navigator 3 | 1.06 | 0.02 |
| *ZNF341* | Zinc finger protein 341 | 1.04 | 0.03 |
| *camsap1* | Calmodulin-regulated spectrin-associated protein 1 | 1.04 | 0.04 |
| *Irgc* | Interferon-inducible GTPase 5 | 1.04 | 0.04 |
| *.* | Oocyte zinc finger protein XlCOF22 | 1.03 | 0.05 |
| *PLEKHA7* | Pleckstrin homology domain-containing family A member 7 | 1.02 | 0.04 |
| *CCDC150* | Coiled-coil domain-containing protein 150 | 1.02 | 0.05 |
| *NOP56* | Nucleolar protein 56 | 1.00 | 0.02 |
| *WDR33* | pre-mRNA 3' end processing protein WDR33 | 1.00 | 0.03 |
| *UPF1* | Regulator of nonsense transcripts 1 | 0.99 | 0.04 |
| *CK095* | Uncharacterized protein C11orf95 | 0.99 | 0.04 |
| *BTAF1* | TATA-binding protein-associated factor 172 | 0.99 | 0.05 |
| *.* | . | 0.98 | 0.05 |
| *NOP58* | Nucleolar protein 58 | 0.96 | 0.05 |
| *impdh1a* | Inosine-5'-monophosphate dehydrogenase 1a | 0.96 | 0.03 |
| *DYR1A* | Dual specificity tyrosine-phosphorylation-regulated kinase 1A | 0.95 | 0.04 |
| *MOV10* | Putative helicase MOV-10 | 0.87 | 0.05 |

Table AI2. Core (QLD) down-regulation

| Gene | Protein | effect | *p* |
| --- | --- | --- | --- |
| *.* | LINE-1 reverse transcriptase homolog | -1.88 | 0.00 |
| *.* | . | -1.72 | 0.00 |
| *CLUL1* | Clusterin-like protein 1 | -1.64 | 0.00 |
| *.* | . | -1.60 | 0.00 |
| *ERVW-1* | Syncytin-1 | -1.60 | 0.00 |
| *CREG1* | Protein CREG1 | -1.56 | 0.00 |
| *ENDOD1* | Endonuclease domain-containing 1 protein | -1.55 | 0.00 |
| *CTSK* | Cathepsin K | -1.43 | 0.01 |
| *Hmox2* | Heme oxygenase 2 | -1.42 | 0.01 |
| *.* | . | -1.39 | 0.00 |
| *PXN1* | Jeltraxin | -1.38 | 0.01 |
| *.* | **.** | -1.38 | 0.01 |
| *prr5* | Proline-rich protein 5 | -1.36 | 0.01 |
| *CFH* | Complement factor H | -1.33 | 0.02 |
| *Pafah2* | Platelet-activating factor acetylhydrolase 2, cytoplasmic | -1.32 | 0.01 |
| *.* | . | -1.31 | 0.02 |
| *NMT2* | Phosphomethylethanolamine N-methyltransferase | -1.31 | 0.02 |
| *FCGR2* | Low affinity immunoglobulin gamma Fc region receptor II | -1.30 | 0.00 |
| *Tub* | Tubby protein | -1.28 | 0.01 |
| *itln1* | Intelectin-1 | -1.27 | 0.01 |
| *CD200R1B* | Cell surface glycoprotein CD200 receptor 1-B | -1.27 | 0.00 |
| *REEP5* | Receptor expression-enhancing protein 5 | -1.26 | 0.01 |
| *PXN1* | Jeltraxin | -1.26 | 0.01 |
| *pol* | Pol polyprotein | -1.26 | 0.02 |
| *mul1a* | Mitochondrial ubiquitin ligase activator of nfkb 1-A | -1.25 | 0.02 |
| *CYP3A29* | Cytochrome P450 3A29 | -1.25 | 0.01 |
| *ST3GAL6* | Type 2 lactosamine alpha-2,3-sialyltransferase | -1.25 | 0.01 |
| *pol* | Pol polyprotein | -1.25 | 0.02 |
| *MPL* | Thrombopoietin receptor | -1.25 | 0.02 |
| *Gp1bb* | Platelet glycoprotein Ib beta chain | -1.23 | 0.02 |
| *.* | . | -1.21 | 0.02 |
| *U88* | Uncharacterized protein U88 | -1.20 | 0.03 |
| *Arsa* | Arylsulfatase A | -1.18 | 0.01 |
| *St3gal5* | Lactosylceramide alpha-2,3-sialyltransferase | -1.18 | 0.04 |
| *PXDC1* | PX domain-containing protein 1 | -1.17 | 0.03 |
| *ERVPABLB-1* | Endogenous retrovirus group PABLB member 1 Env polyprotein | -1.16 | 0.03 |
| *LORF2* | LINE-1 retrotransposable element ORF2 protein | -1.15 | 0.01 |
| *.* | . | -1.15 | 0.02 |
| *.* | . | -1.15 | 0.01 |
| *F10* | Coagulation factor X | -1.14 | 0.02 |
| *CYR61* | Protein CYR61 | -1.14 | 0.02 |
| *TRIM25* | E3 ubiquitin/ISG15 ligase TRIM25 | -1.14 | 0.04 |
| *MED12L* | Mediator of RNA polymerase II transcription subunit 12-like protein | -1.13 | 0.03 |
| *Pinlyp* | phospholipase A2 inhibitor and Ly6/PLAUR domain-containing protein | -1.12 | 0.03 |
| *pssA* | CDP-diacylglycerol--serine O-phosphatidyltransferase | -1.11 | 0.04 |
| *ARHGEF19* | Rho guanine nucleotide exchange factor 19 | -1.11 | 0.03 |
| *TESPA1* | Protein TESPA1 | -1.10 | 0.02 |
| *Gp9* | Platelet glycoprotein IX | -1.09 | 0.03 |
| *ITGA2B* | Integrin alpha-IIb | -1.09 | 0.03 |
| *Dhrs9* | Dehydrogenase/reductase SDR family member 9 | -1.09 | 0.03 |
| *.* | . | -1.08 | 0.04 |
| *PTPRM* | Receptor-type tyrosine-protein phosphatase mu | -1.08 | 0.02 |
| *.* | . | -1.08 | 0.04 |
| *.* | **.** | -1.08 | 0.05 |
| *.* | . | -1.07 | 0.03 |
| *fn1* | Fibronectin | -1.07 | 0.02 |
| *Ecm1* | Extracellular matrix protein 1 | -1.07 | 0.03 |
| *CCDC126* | Coiled-coil domain-containing protein 126 | -1.06 | 0.04 |
| *Igk-V19-17* | Ig kappa chain V19-17 | -1.06 | 0.03 |
| *ZNF84* | Zinc finger protein 84 | -1.06 | 0.02 |
| *VOPP1* | Vesicular, overexpressed in cancer, prosurvival protein 1 | -1.05 | 0.05 |
| *Mtcl1* | Microtubule cross-linking factor 1 | -1.05 | 0.04 |
| *c1galt1* | Glycoprotein-N-acetylgalactosamine 3-beta-galactosyltransferase 1 | -1.04 | 0.02 |
| *Rtn4rl1* | Reticulon-4 receptor-like 1 | -1.04 | 0.03 |
| *Dcst1* | DC-STAMP domain-containing protein 1 | -1.03 | 0.04 |
| *liph-a* | Lipase member H-A | -1.03 | 0.03 |
| *.* | . | -1.03 | 0.04 |
| *ME1* | NADP-dependent malic enzyme | -1.03 | 0.04 |
| *GPR21* | Probable G-protein coupled receptor 21 | -1.02 | 0.03 |
| *FNDC7* | Fibronectin type III domain-containing protein 7 | -1.00 | 0.04 |
| *IGKV6-21* | Immunoglobulin kappa variable 6-21 | -1.00 | 0.04 |
| *CSF2RA* | Granulocyte-macrophage colony-stimulating factor receptor subunit alpha | -1.00 | 0.04 |
| *IIGP5* | Interferon-inducible GTPase 5 | -1.00 | 0.02 |
| *FCN1* | Ficolin-1 | -1.00 | 0.04 |
| *P2RY12* | P2Y purinoceptor 12 | -0.99 | 0.04 |
| *Nptn* | Neuroplastin | -0.98 | 0.04 |
| *.* | **.** | -0.97 | 0.03 |
| *ARHGEF37* | Rho guanine nucleotide exchange factor 37 | -0.97 | 0.04 |
| *.* | . | -0.97 | 0.04 |
| *Esm1* | Endothelial cell-specific molecule 1 | -0.96 | 0.03 |
| *Gp5* | Platelet glycoprotein V | -0.95 | 0.04 |
| *CYP8B1* | 5-beta-cholestane-3-alpha,7-alpha-diol 12-alpha-hydroxylase | -0.95 | 0.04 |
| *lingo1* | Leucine-rich repeat and immunoglobulin-like domain-containing nogo receptor-interacting protein 1 | -0.95 | 0.04 |
| *PXN1* | Jeltraxin | -0.93 | 0.04 |
| *.* | . | -0.91 | 0.04 |
| *NUCB2* | Nucleobindin-2 | -0.90 | 0.02 |
| *NXPE3* | NXPE family member 3 | -0.90 | 0.03 |
| *.* | **.** | -0.87 | 0.04 |
| *.* | . | -0.81 | 0.05 |
| *HLA-DRB1* | HLA class II histocompatibility antigen, DRB1-4 beta chain | -0.80 | 0.04 |
| *.* | **.** | -0.74 | 0.04 |

Table AI3. Intermediate (NT) up-regulation

| Gene | Protein | effect | *p* |
| --- | --- | --- | --- |
| *prkra-a* | Interferon-inducible double-stranded RNA-dependent protein kinase activator A homolog A | 2.10 | 0.00 |
| *ISG20L2* | Interferon-stimulated 20 kDa exonuclease-like 2 | 2.03 | 0.00 |
| *.* | . | 1.96 | 0.00 |
| *.* | . | 1.94 | 0.00 |
| *DNAJA2* | DnaJ homolog subfamily A member 2 | 1.87 | 0.00 |
| *DLST* | Dihydrolipoyllysine-residue succinyltransferase component of 2-oxoglutarate dehydrogenase complex, mitochondrial | 1.87 | 0.00 |
| *.* | . | 1.83 | 0.00 |
| *.* | . | 1.81 | 0.00 |
| *gsn* | Gelsolin | 1.80 | 0.00 |
| *.* | . | 1.73 | 0.00 |
| *Serbp1* | Plasminogen activator inhibitor 1 RNA-binding protein | 1.73 | 0.00 |
| *rps8* | 40S ribosomal protein S8 | 1.72 | 0.00 |
| *Fip1l1* | Pre-mRNA 3'-end-processing factor FIP1 | 1.71 | 0.00 |
| *Smarce1* | SWI/SNF-related matrix-associated actin-dependent regulator of chromatin subfamily E member 1 | 1.70 | 0.00 |
| *TPT1* | Translationally-controlled tumor protein homolog | 1.67 | 0.00 |
| *tram1l1* | . | 1.65 | 0.00 |
| *rps2* | 40S ribosomal protein S2 | 1.64 | 0.00 |
| *tmem251* | Transmembrane protein 251 | 1.64 | 0.00 |
| *Pnrc1* | Proline-rich nuclear receptor coactivator 1 | 1.64 | 0.00 |
| *MRPL35* | 39S ribosomal protein L35, mitochondrial | 1.64 | 0.00 |
| *KCTD2* | BTB/POZ domain-containing protein KCTD2 | 1.63 | 0.00 |
| *.* | . | 1.63 | 0.01 |
| *.* | . | 1.55 | 0.01 |
| *GLTSCR2* | Glioma tumor suppressor candidate region gene 2 protein | 1.54 | 0.00 |
| *B4galt3* | Beta-1,4-galactosyltransferase 3 | 1.54 | 0.00 |
| *TMED10* | Transmembrane emp24 domain-containing protein 10 | 1.53 | 0.00 |
| *ppp4c* | Serine/threonine-protein phosphatase 4 catalytic subunit | 1.53 | 0.00 |
| *.* | . | 1.51 | 0.00 |
| *PGM1* | Phosphoglucomutase-1 | 1.51 | 0.00 |
| *Csnk2a2* | Casein kinase II subunit alpha' | 1.51 | 0.00 |
| *EEF1A* | Elongation factor 1-alpha 1 | 1.50 | 0.00 |
| *.* | . | 1.50 | 0.00 |
| *PSMC3* | 26S protease regulatory subunit 6A | 1.49 | 0.00 |
| *Pafah2* | Platelet-activating factor acetylhydrolase 2, cytoplasmic | 1.49 | 0.01 |
| *.* | . | 1.48 | 0.00 |
| *ITM2A* | Integral membrane protein 2A | 1.48 | 0.00 |
| *rpl7a* | 60S ribosomal protein L7a | 1.47 | 0.00 |
| *EEF2* | Elongation factor 2 | 1.47 | 0.00 |
| *METTL9* | Methyltransferase-like protein 9 | 1.47 | 0.01 |
| *.* | Heterogeneous nuclear ribonucleoprotein A3 homolog 1 | 1.46 | 0.00 |
| *MCUB* | Calcium uniporter regulatory subunit MCUb, mitochondrial | 1.46 | 0.01 |
| *PSMB5* | Proteasome subunit beta type-5 | 1.45 | 0.01 |
| *pkmyt1* | Membrane-associated tyrosine- and threonine-specific cdc2-inhibitory kinase | 1.45 | 0.00 |
| *smyd5* | SET and MYND domain-containing protein 5 | 1.44 | 0.00 |
| *.* | . | 1.42 | 0.01 |
| *FBXL3* | F-box/LRR-repeat protein 3 | 1.42 | 0.01 |
| *.* | . | 1.42 | 0.01 |
| *eif3g-b* | Eukaryotic translation initiation factor 3 subunit G-B | 1.42 | 0.00 |
| *.* | . | 1.42 | 0.01 |
| *eef1g-a* | Elongation factor 1-gamma-A | 1.41 | 0.00 |
| *.* | Oocyte zinc finger protein XlCOF7.1 | 1.39 | 0.01 |
| *.* | . | 1.38 | 0.01 |
| *eif3h* | Eukaryotic translation initiation factor 3 subunit H | 1.37 | 0.00 |
| *med8-b* | Mediator of RNA polymerase II transcription subunit 8-B | 1.36 | 0.00 |
| *ZNF567* | Zinc finger protein 567 | 1.36 | 0.00 |
| *.* | . | 1.35 | 0.01 |
| *Rack1* | Receptor of activated protein C kinase 1 | 1.35 | 0.00 |
| *ispd* | Isoprenoid synthase domain-containing protein | 1.35 | 0.01 |
| *.* | . | 1.34 | 0.00 |
| *NDUFV2* | NADH dehydrogenase [ubiquinone] flavoprotein 2, mitochondrial | 1.34 | 0.01 |
| *.* | . | 1.34 | 0.00 |
| *NKAP* | NF-kappa-B-activating protein | 1.34 | 0.00 |
| *.* | . | 1.34 | 0.01 |
| *.* | Gastrula zinc finger protein XlCGF17.1 | 1.33 | 0.01 |
| *.* | . | 1.33 | 0.00 |
| *PMVK* | Phosphomevalonate kinase | 1.33 | 0.01 |
| *RABEP2* | Rab GTPase-binding effector protein 2 | 1.33 | 0.02 |
| *Gpbp1* | Vasculin | 1.33 | 0.01 |
| *Xrcc6* | X-ray repair cross-complementing protein 6 | 1.33 | 0.01 |
| *Atp5o* | ATP synthase subunit O, mitochondrial | 1.33 | 0.01 |
| *.* | . | 1.31 | 0.01 |
| *Rps27* | 40S ribosomal protein S27 | 1.31 | 0.00 |
| *Rps3* | 40S ribosomal protein S3 | 1.30 | 0.00 |
| *Mrps10* | 28S ribosomal protein S10, mitochondrial | 1.30 | 0.01 |
| *Eif2b4* | Translation initiation factor eIF-2B subunit delta | 1.30 | 0.02 |
| *ART2* | Putative uncharacterized protein ART2 | 1.29 | 0.00 |
| *rps3a* | 40S ribosomal protein S3a | 1.29 | 0.01 |
| *.* | Tubulin alpha chain | 1.29 | 0.01 |
| *Gabarap* | Gamma-aminobutyric acid receptor-associated protein | 1.28 | 0.01 |
| *get4* | Golgi to ER traffic protein 4 homolog | 1.28 | 0.01 |
| *.* | . | 1.28 | 0.01 |
| *prr5* | Proline-rich protein 5 | 1.28 | 0.01 |
| *rsph9* | Radial spoke head protein 9 homolog | 1.28 | 0.01 |
| *Rps19* | 40S ribosomal protein S19 | 1.27 | 0.00 |
| *eif3a* | Eukaryotic translation initiation factor 3 subunit A | 1.27 | 0.01 |
| *.* | . | 1.27 | 0.01 |
| *.* | . | 1.27 | 0.00 |
| *Eef1d* | Elongation factor 1-delta | 1.27 | 0.01 |
| *rps23* | 40S ribosomal protein S23 | 1.27 | 0.01 |
| *nae1* | NEDD8-activating enzyme E1 regulatory subunit | 1.27 | 0.01 |
| *IFITM10* | Interferon-induced transmembrane protein 10 | 1.27 | 0.01 |
| *.* | . | 1.26 | 0.01 |
| *ART2* | Putative uncharacterized protein ART2 | 1.26 | 0.00 |
| *PIGF* | Phosphatidylinositol-glycan biosynthesis class F protein | 1.26 | 0.01 |
| *PDE12* | 2',5'-phosphodiesterase 12 | 1.26 | 0.03 |
| *.* | . | 1.25 | 0.02 |
| *.* | . | 1.25 | 0.02 |
| *GORASP2* | Golgi reassembly-stacking protein 2 | 1.25 | 0.01 |
| *.* | . | 1.25 | 0.02 |
| *Cox11* | Cytochrome c oxidase assembly protein COX11, mitochondrial | 1.25 | 0.01 |
| *.* | . | 1.24 | 0.02 |
| *TCP1* | T-complex protein 1 subunit alpha | 1.24 | 0.01 |
| *ART2* | Putative uncharacterized protein ART2 | 1.24 | 0.00 |
| *.* | . | 1.23 | 0.01 |
| *FAM47C* | Putative protein FAM47C | 1.23 | 0.00 |
| *Pol* | LINE-1 retrotransposable element ORF2 protein | 1.23 | 0.00 |
| *S100A11* | Protein S100-A11 | 1.22 | 0.02 |
| *rpl4-b* | 60S ribosomal protein L4-B | 1.22 | 0.01 |
| *ZNF300* | Zinc finger protein 300 | 1.22 | 0.02 |
| *pssA* | CDP-diacylglycerol--serine O-phosphatidyltransferase | 1.21 | 0.01 |
| *.* | . | 1.21 | 0.01 |
| *ZNF300* | Zinc finger protein 300 | 1.21 | 0.01 |
| *Glo1* | Lactoylglutathione lyase | 1.21 | 0.01 |
| *.* | . | 1.21 | 0.02 |
| *.* | . | 1.20 | 0.02 |
| *SLC24A2* | Sodium/potassium/calcium exchanger 2 | 1.20 | 0.01 |
| *yipf3* | Protein YIPF3 | 1.20 | 0.01 |
| *.* | . | 1.19 | 0.02 |
| *Nlrc3* | Protein NLRC3 | 1.18 | 0.00 |
| *MTMR9* | Myotubularin-related protein 9 | 1.18 | 0.01 |
| *Eloc* | Elongin-C | 1.18 | 0.03 |
| *ndor1* | NADPH-dependent diflavin oxidoreductase 1 | 1.18 | 0.01 |
| *.* | . | 1.18 | 0.02 |
| *ERI3* | ERI1 exoribonuclease 3 | 1.18 | 0.02 |
| *pno1* | RNA-binding protein PNO1 | 1.17 | 0.01 |
| *TXNDC5* | Thioredoxin domain-containing protein 5 | 1.17 | 0.02 |
| *RPL3* | 60S ribosomal protein L3 | 1.17 | 0.01 |
| *.* | . | 1.17 | 0.03 |
| *Agr3* | Anterior gradient protein 3 | 1.17 | 0.03 |
| *Borcs6* | BLOC-1-related complex subunit 6 | 1.17 | 0.01 |
| *NDUFS8* | NADH dehydrogenase [ubiquinone] iron-sulfur protein 8, mitochondrial | 1.17 | 0.02 |
| *.* | . | 1.17 | 0.00 |
| *NAXD* | ATP-dependent (S)-NAD(P)H-hydrate dehydratase | 1.16 | 0.01 |
| *rnf2-a* | E3 ubiquitin-protein ligase RING2-A | 1.16 | 0.01 |
| *med9* | Mediator of RNA polymerase II transcription subunit 9 | 1.16 | 0.02 |
| *.* | . | 1.16 | 0.02 |
| *ELMO2* | Engulfment and cell motility protein 2 | 1.16 | 0.02 |
| *.* | Nuclear factor 7, ovary | 1.16 | 0.02 |
| *.* | . | 1.16 | 0.02 |
| *ATF4* | Cyclic AMP-dependent transcription factor ATF-4 | 1.16 | 0.03 |
| *Vamp2* | Vesicle-associated membrane protein 2 | 1.15 | 0.02 |
| *HSP90AB1* | Heat shock cognate protein HSP 90-beta | 1.15 | 0.01 |
| *CCT6* | T-complex protein 1 subunit zeta | 1.15 | 0.01 |
| *eif3d* | Eukaryotic translation initiation factor 3 subunit D | 1.15 | 0.01 |
| *.* | . | 1.15 | 0.02 |
| *.* | . | 1.14 | 0.02 |
| *.* | . | 1.14 | 0.03 |
| *LCMT1* | Leucine carboxyl methyltransferase 1 | 1.14 | 0.01 |
| *.* | . | 1.14 | 0.02 |
| *.* | . | 1.14 | 0.03 |
| *ybx1* | Nuclease-sensitive element-binding protein 1 | 1.14 | 0.01 |
| *Gtpbp4* | Nucleolar GTP-binding protein 1 | 1.14 | 0.01 |
| *.* | . | 1.14 | 0.02 |
| *ciapin1* | Anamorsin | 1.14 | 0.02 |
| *.* | . | 1.14 | 0.02 |
| *.* | . | 1.13 | 0.02 |
| *pi4kb* | Phosphatidylinositol 4-kinase beta | 1.13 | 0.02 |
| *.* | . | 1.12 | 0.03 |
| *ZNF268* | Zinc finger protein 268 | 1.12 | 0.03 |
| *Pkd1l3* | Polycystic kidney disease protein 1-like 3 | 1.12 | 0.02 |
| *ORF V* | Enzymatic polyprotein | 1.12 | 0.02 |
| *Rpl17* | 60S ribosomal protein L17 | 1.12 | 0.01 |
| *fbxl5* | F-box/LRR-repeat protein 5 | 1.12 | 0.01 |
| *rps15* | 40S ribosomal protein S15 | 1.12 | 0.02 |
| *.* | . | 1.11 | 0.01 |
| *RPL10* | 60S ribosomal protein L10 | 1.11 | 0.01 |
| *Tmed8* | Protein TMED8 | 1.11 | 0.03 |
| *eif3b* | Eukaryotic translation initiation factor 3 subunit B | 1.11 | 0.01 |
| *ACADVL* | Very long-chain specific acyl-CoA dehydrogenase, mitochondrial | 1.11 | 0.02 |
| *.* | . | 1.11 | 0.02 |
| *.* | . | 1.11 | 0.02 |
| *sde2* | Protein SDE2 homolog | 1.10 | 0.02 |
| *LRRC41* | Leucine-rich repeat-containing protein 41 | 1.10 | 0.03 |
| *QPCTL* | Glutaminyl-peptide cyclotransferase-like protein | 1.10 | 0.02 |
| *GARS* | Glycine--tRNA ligase | 1.10 | 0.05 |
| *Rpl37* | 60S ribosomal protein L37 | 1.10 | 0.02 |
| *aad-a* | Alpha-aspartyl dipeptidase | 1.10 | 0.02 |
| *Rpl11* | 60S ribosomal protein L11 | 1.09 | 0.02 |
| *RWDD1* | RWD domain-containing protein 1 | 1.09 | 0.01 |
| *eif3l* | Eukaryotic translation initiation factor 3 subunit L | 1.09 | 0.01 |
| *.* | . | 1.09 | 0.03 |
| *rpl18-b* | 60S ribosomal protein L18-B | 1.09 | 0.02 |
| *Psmc5* | 26S protease regulatory subunit 8 | 1.09 | 0.04 |
| *rpl18a* | 60S ribosomal protein L18a | 1.09 | 0.02 |
| *.* | . | 1.09 | 0.03 |
| *.* | . | 1.09 | 0.04 |
| *CLPP* | ATP-dependent Clp protease proteolytic subunit, mitochondrial | 1.09 | 0.03 |
| *Rpl23* | 60S ribosomal protein L23 | 1.08 | 0.02 |
| *Tuba1b* | Tubulin alpha-1B chain | 1.08 | 0.03 |
| *ccdc93* | Coiled-coil domain-containing protein 93 | 1.08 | 0.02 |
| *.* | . | 1.08 | 0.03 |
| *ZNF41* | Zinc finger protein 41 | 1.08 | 0.04 |
| *PSMC4* | 26S protease regulatory subunit 6B | 1.08 | 0.01 |
| *RSPO3* | R-spondin-3 | 1.08 | 0.02 |
| *STARD6* | StAR-related lipid transfer protein 6 | 1.08 | 0.02 |
| *.* | . | 1.08 | 0.04 |
| *.* | . | 1.08 | 0.03 |
| *TOMM40L* | Mitochondrial import receptor subunit TOM40B | 1.07 | 0.03 |
| *.* | . | 1.07 | 0.03 |
| *PCNP* | PEST proteolytic signal-containing nuclear protein | 1.07 | 0.03 |
| *RPL13A* | 60S ribosomal protein L13a | 1.07 | 0.02 |
| *MRPL2* | 39S ribosomal protein L2, mitochondrial | 1.07 | 0.03 |
| *PFDN6* | Prefoldin subunit 6 | 1.07 | 0.02 |
| *.* | . | 1.07 | 0.03 |
| *Dph5* | Diphthine methyl ester synthase | 1.07 | 0.02 |
| *UBQLN4* | Ubiquilin-4 | 1.07 | 0.02 |
| *.* | Nucleoside diphosphate kinase A1 | 1.07 | 0.01 |
| *ARPC1A* | Actin-related protein 2/3 complex subunit 1A | 1.07 | 0.04 |
| *CTDSP2* | Carboxy-terminal domain RNA polymerase II polypeptide A small phosphatase 2 | 1.06 | 0.03 |
| *infB* | Translation initiation factor IF-2 | 1.06 | 0.03 |
| *RpL13* | 60S ribosomal protein L13 | 1.06 | 0.02 |
| *Abra* | Actin-binding Rho-activating protein | 1.06 | 0.02 |
| *.* | . | 1.06 | 0.03 |
| *ddost* | Dolichyl-diphosphooligosaccharide--protein glycosyltransferase 48 kDa subunit | 1.06 | 0.01 |
| *.* | . | 1.06 | 0.04 |
| *CD200R1B* | Cell surface glycoprotein CD200 receptor 1-B | 1.06 | 0.04 |
| *EIF2B3* | Translation initiation factor eIF-2B subunit gamma | 1.05 | 0.02 |
| *tmem147* | Transmembrane protein 147 | 1.05 | 0.03 |
| *svbp* | Small vasohibin-binding protein | 1.05 | 0.03 |
| *.* | . | 1.05 | 0.02 |
| *PLEK* | Pleckstrin | 1.05 | 0.04 |
| *.* | . | 1.05 | 0.03 |
| *.* | . | 1.05 | 0.04 |
| *aida-b* | Axin interactor, dorsalization-associated protein B | 1.05 | 0.05 |
| *IMMT* | MICOS complex subunit MIC60 | 1.05 | 0.04 |
| *TBXAS1* | Thromboxane-A synthase | 1.05 | 0.03 |
| *.* | Gastrula zinc finger protein XlCGF57.1 | 1.04 | 0.02 |
| *EIF3F* | Eukaryotic translation initiation factor 3 subunit F | 1.04 | 0.04 |
| *.* | . | 1.04 | 0.04 |
| *alkbh5* | RNA demethylase ALKBH5 | 1.04 | 0.01 |
| *NDUFB9* | NADH dehydrogenase [ubiquinone] 1 beta subcomplex subunit 9 | 1.04 | 0.02 |
| *PYCARD* | Apoptosis-associated speck-like protein containing a CARD | 1.04 | 0.04 |
| *Ccdc58* | Coiled-coil domain-containing protein 58 | 1.04 | 0.04 |
| *Rpl12* | 60S ribosomal protein L12 | 1.04 | 0.02 |
| *Rpl17* | 60S ribosomal protein L17 | 1.03 | 0.03 |
| *TAR1* | Protein TAR1 | 1.03 | 0.05 |
| *.* | . | 1.03 | 0.03 |
| *ak2* | Adenylate kinase 2, mitochondrial | 1.03 | 0.04 |
| *.* | Oocyte zinc finger protein XlCOF7.1 | 1.03 | 0.03 |
| *SLC35A1* | CMP-sialic acid transporter | 1.03 | 0.02 |
| *.* | Gastrula zinc finger protein XlCGF8.2DB | 1.03 | 0.04 |
| *PSMD4* | 26S proteasome non-ATPase regulatory subunit 4 | 1.03 | 0.04 |
| *coil* | Coilin | 1.03 | 0.04 |
| *ZNF84* | Zinc finger protein 84 | 1.03 | 0.04 |
| *.* | Gastrula zinc finger protein XlCGF67.1 | 1.03 | 0.03 |
| *.* | . | 1.02 | 0.04 |
| *.* | . | 1.02 | 0.03 |
| *Stt3a* | Dolichyl-diphosphooligosaccharide--protein glycosyltransferase subunit STT3A | 1.02 | 0.02 |
| *Tmem115* | Transmembrane protein 115 | 1.02 | 0.04 |
| *MOSPD1* | Motile sperm domain-containing protein 1 | 1.02 | 0.03 |
| *pgap2* | Post-GPI attachment to proteins factor 2 | 1.02 | 0.03 |
| *BEND4* | BEN domain-containing protein 4 | 1.02 | 0.04 |
| *hexim* | Protein HEXIM | 1.02 | 0.03 |
| *impdh2* | Inosine-5'-monophosphate dehydrogenase 2 | 1.02 | 0.03 |
| *HAX1* | HCLS1-associated protein X-1 | 1.02 | 0.02 |
| *.* | . | 1.02 | 0.02 |
| *NGB* | Neuroglobin | 1.02 | 0.03 |
| *.* | . | 1.02 | 0.05 |
| *ddx21-b* | Nucleolar RNA helicase 2-B | 1.02 | 0.02 |
| *Paics* | Multifunctional protein ADE2 | 1.02 | 0.05 |
| *exosc6* | Exosome complex component MTR3 | 1.02 | 0.03 |
| *GPR21* | Probable G-protein coupled receptor 21 | 1.01 | 0.03 |
| *.* | . | 1.01 | 0.04 |
| *ecsit* | Evolutionarily conserved signaling intermediate in Toll pathway, mitochondrial | 1.01 | 0.04 |
| *.* | . | 1.01 | 0.03 |
| *.* | . | 1.01 | 0.04 |
| *Psmg3* | Proteasome assembly chaperone 3 | 1.00 | 0.03 |
| *.* | . | 1.00 | 0.05 |
| *ZNF8* | Zinc finger protein 8 | 1.00 | 0.03 |
| *rps4* | 40S ribosomal protein S4 | 1.00 | 0.03 |
| *Terb2* | Telomere repeats-binding bouquet formation protein 2 | 1.00 | 0.04 |
| *rpl7a* | 60S ribosomal protein L7a | 1.00 | 0.02 |
| *TXNL1* | Thioredoxin-like protein 1 | 1.00 | 0.03 |
| *.* | . | 1.00 | 0.05 |
| *CD2* | T-cell surface antigen CD2 | 1.00 | 0.04 |
| *sh3bp5l* | SH3 domain-binding protein 5-like | 1.00 | 0.03 |
| *.* | . | 1.00 | 0.03 |
| *EIF2S3* | Eukaryotic translation initiation factor 2 subunit 3 | 0.99 | 0.02 |
| *DHCR24* | Delta(24)-sterol reductase | 0.99 | 0.03 |
| *PLCL1* | Inactive phospholipase C-like protein 1 | 0.99 | 0.05 |
| *LRSAM1* | E3 ubiquitin-protein ligase LRSAM1 | 0.99 | 0.02 |
| *.* | . | 0.99 | 0.04 |
| *.* | . | 0.99 | 0.03 |
| *CIR1* | Corepressor interacting with RBPJ 1 | 0.99 | 0.03 |
| *Acads* | Short-chain specific acyl-CoA dehydrogenase, mitochondrial | 0.99 | 0.04 |
| *ZNF484* | Zinc finger protein 484 | 0.99 | 0.04 |
| *.* | . | 0.98 | 0.03 |
| *RPL35* | 60S ribosomal protein L35 | 0.98 | 0.05 |
| *.* | . | 0.98 | 0.02 |
| *TTLL9* | Probable tubulin polyglutamylase TTLL9 | 0.98 | 0.05 |
| *ASB6* | Ankyrin repeat and SOCS box protein 6 | 0.98 | 0.04 |
| *SLC27A3* | Long-chain fatty acid transport protein 3 | 0.98 | 0.02 |
| *glmp-a* | Glycosylated lysosomal membrane protein A | 0.98 | 0.04 |
| *.* | . | 0.98 | 0.03 |
| *naxe* | NAD(P)H-hydrate epimerase | 0.98 | 0.05 |
| *MIEN1* | Migration and invasion enhancer 1 | 0.98 | 0.04 |
| *EIF5A2* | Eukaryotic translation initiation factor 5A-2 | 0.97 | 0.04 |
| *RAB24* | Ras-related protein Rab-24 | 0.97 | 0.03 |
| *eif2a* | Eukaryotic translation initiation factor 2A | 0.97 | 0.03 |
| *.* | . | 0.97 | 0.04 |
| *nudt19* | Nucleoside diphosphate-linked moiety X motif 19 | 0.97 | 0.04 |
| *SLC5A3* | Sodium/myo-inositol cotransporter | 0.97 | 0.03 |
| *PHF1* | PHD finger protein 1 | 0.97 | 0.04 |
| *CCT4* | T-complex protein 1 subunit delta | 0.97 | 0.04 |
| *nmd3* | 60S ribosomal export protein NMD3 | 0.97 | 0.04 |
| *ZNF586* | Zinc finger protein 586 | 0.97 | 0.04 |
| *eif3k* | Eukaryotic translation initiation factor 3 subunit K | 0.97 | 0.04 |
| *Tbl1x* | F-box-like/WD repeat-containing protein TBL1X | 0.97 | 0.04 |
| *PNPO* | Pyridoxine-5'-phosphate oxidase | 0.97 | 0.04 |
| *Atp5i* | ATP synthase subunit e, mitochondrial | 0.97 | 0.03 |
| *.* | . | 0.96 | 0.03 |
| *ostc* | Oligosaccharyltransferase complex subunit ostc | 0.96 | 0.03 |
| *.* | . | 0.96 | 0.05 |
| *Ier5* | Immediate early response gene 5 protein | 0.96 | 0.04 |
| *POLM* | DNA-directed DNA/RNA polymerase mu | 0.96 | 0.03 |
| *.* | . | 0.96 | 0.04 |
| *MOCS2* | Molybdopterin synthase catalytic subunit | 0.96 | 0.04 |
| *.* | . | 0.96 | 0.03 |
| *.* | . | 0.96 | 0.05 |
| *.* | . | 0.96 | 0.03 |
| *fam32a* | Protein FAM32A | 0.96 | 0.04 |
| *CCDC13* | Coiled-coil domain-containing protein 13 | 0.96 | 0.03 |
| *ORC3* | Origin recognition complex subunit 3 | 0.96 | 0.03 |
| *MVK* | Mevalonate kinase | 0.96 | 0.04 |
| *.* | . | 0.95 | 0.04 |
| *Fxyd1* | Phospholemman | 0.95 | 0.04 |
| *nubp1-A* | Cytosolic Fe-S cluster assembly factor nubp1-A | 0.95 | 0.02 |
| *PRKACA* | cAMP-dependent protein kinase catalytic subunit alpha | 0.95 | 0.03 |
| *Eef1a2* | Elongation factor 1-alpha 2 | 0.95 | 0.02 |
| *.* | . | 0.95 | 0.03 |
| *hacd4* | Very-long-chain (3R)-3-hydroxyacyl-CoA dehydratase | 0.95 | 0.04 |
| *.* | . | 0.95 | 0.04 |
| *pxn1* | Pentraxin fusion protein | 0.95 | 0.05 |
| *Kars* | Lysine--tRNA ligase | 0.95 | 0.04 |
| *Polr2c* | DNA-directed RNA polymerase II subunit RPB3 | 0.94 | 0.04 |
| *Was* | Wiskott-Aldrich syndrome protein homolog | 0.94 | 0.02 |
| *Dnajc4* | DnaJ homolog subfamily C member 4 | 0.94 | 0.04 |
| *.* | . | 0.94 | 0.03 |
| *CRLS1* | Cardiolipin synthase (CMP-forming) | 0.94 | 0.04 |
| *GFM2* | Ribosome-releasing factor 2, mitochondrial | 0.94 | 0.05 |
| *.* | . | 0.94 | 0.04 |
| *RNF40* | E3 ubiquitin-protein ligase BRE1B | 0.94 | 0.04 |
| *ZNF300* | Zinc finger protein 300 | 0.94 | 0.04 |
| *RPLP0* | 60S acidic ribosomal protein P0 | 0.94 | 0.05 |
| *ZCHC3* | Zinc finger CCHC domain-containing protein 3 | 0.94 | 0.05 |
| *.* | . | 0.94 | 0.04 |
| *Ssr4* | Translocon-associated protein subunit delta | 0.94 | 0.04 |
| *.* | Cytochrome c oxidase subunit 4 isoform 2, mitochondrial | 0.94 | 0.04 |
| *Slc35b2* | Adenosine 3'-phospho 5'-phosphosulfate transporter 1 | 0.93 | 0.04 |
| *RPS14* | 40S ribosomal protein S14 | 0.93 | 0.03 |
| *THAP5* | THAP domain-containing protein 5 | 0.93 | 0.04 |
| *MPL* | Thrombopoietin receptor | 0.93 | 0.04 |
| *ELOVL1* | Elongation of very long chain fatty acids protein 1 | 0.93 | 0.04 |
| *Nlrc3* | Protein NLRC3 | 0.93 | 0.01 |
| *.* | . | 0.93 | 0.03 |
| *ZNF300* | Zinc finger protein 300 | 0.92 | 0.04 |
| *rpl8* | 60S ribosomal protein L8 | 0.92 | 0.03 |
| *Fhl1* | Four and a half LIM domains protein 1 | 0.92 | 0.03 |
| *VIPAS39* | Spermatogenesis-defective protein 39 homolog | 0.92 | 0.04 |
| *rps16* | 40S ribosomal protein S16 | 0.92 | 0.04 |
| *eif3c* | Eukaryotic translation initiation factor 3 subunit C | 0.92 | 0.02 |
| *Rpl32* | 60S ribosomal protein L32 | 0.92 | 0.03 |
| *ANXA1* | Annexin A1 | 0.92 | 0.03 |
| *.* | . | 0.92 | 0.05 |
| *.* | . | 0.91 | 0.04 |
| *Rps9* | 40S ribosomal protein S9 | 0.91 | 0.05 |
| *NSD3* | Histone-lysine N-methyltransferase NSD3 | 0.91 | 0.04 |
| *mvp* | Major vault protein | 0.91 | 0.02 |
| *SUGP1* | SURP and G-patch domain-containing protein 1 | 0.91 | 0.04 |
| *.* | . | 0.91 | 0.05 |
| *RAB10* | Ras-related protein Rab-10 | 0.91 | 0.03 |
| *.* | Gastrula zinc finger protein XlCGF57.1 | 0.90 | 0.04 |
| *AL3B1* | Aldehyde dehydrogenase family 3 member B1 | 0.90 | 0.05 |
| *RPS5* | 40S ribosomal protein S5 | 0.90 | 0.03 |
| *.* | . | 0.90 | 0.04 |
| *.* | . | 0.90 | 0.05 |
| *UBXN2A* | UBX domain-containing protein 2A | 0.90 | 0.04 |
| *cct3* | T-complex protein 1 subunit gamma | 0.90 | 0.04 |
| *Y325* | Uncharacterized oxidoreductase TM_0325 | 0.89 | 0.05 |
| *pld3* | Phospholipase D3 | 0.89 | 0.05 |
| *.* | . | 0.89 | 0.05 |
| *p33monox* | Putative monooxygenase p33MONOX | 0.89 | 0.04 |
| *Rpl27* | 60S ribosomal protein L27 | 0.89 | 0.04 |
| *SEC22A* | Vesicle-trafficking protein SEC22a | 0.89 | 0.04 |
| *TESK2* | Dual specificity testis-specific protein kinase 2 | 0.89 | 0.04 |
| *CPA3* | Mast cell carboxypeptidase A | 0.87 | 0.05 |
| *gkap1* | G kinase-anchoring protein 1 | 0.87 | 0.04 |
| *.* | . | 0.86 | 0.04 |
| *PDCL* | Phosducin-like protein | 0.86 | 0.05 |
| *SLC25A3* | Phosphate carrier protein, mitochondrial | 0.86 | 0.04 |
| *Imp3* | U3 small nucleolar ribonucleoprotein protein IMP3 | 0.85 | 0.04 |
| *spice1* | Spindle and centriole-associated protein 1 | 0.85 | 0.04 |
| *ABHD16A* | Protein ABHD16A | 0.85 | 0.03 |
| *.* | . | 0.85 | 0.05 |
| *emc3* | ER membrane protein complex subunit 3 | 0.85 | 0.03 |
| *DCTN3* | Dynactin subunit 3 | 0.85 | 0.05 |
| *.* | . | 0.85 | 0.04 |
| *RNF112* | RING finger protein 112 | 0.85 | 0.04 |
| *.* | . | 0.83 | 0.04 |

Table AI4. Intermediate (NT) down-regulation

| Gene | Protein | effect | *p* |
| --- | --- | --- | --- |
| *.* | . | -7.56 | 0.00 |
| *.* | . | -5.67 | 0.00 |
| *YCX91* | Uncharacterized protein ORF91 | -4.74 | 0.00 |
| *.* | . | -4.39 | 0.00 |
| *YCX91* | Uncharacterized protein ORF91 | -4.16 | 0.00 |
| *.* | . | -3.99 | 0.00 |
| *.* | . | -3.67 | 0.00 |
| *YCX91* | Uncharacterized protein ORF91 | -3.38 | 0.00 |
| *.* | . | -3.16 | 0.00 |
| *.* | . | -3.11 | 0.00 |
| *POL* | Gag-Pol polyprotein | -3.06 | 0.00 |
| *.* | . | -2.99 | 0.00 |
| *.* | . | -2.87 | 0.00 |
| *.* | . | -2.55 | 0.00 |
| *TRMD* | tRNA (guanine-N(1)-)-methyltransferase | -2.53 | 0.00 |
| *RPOA* | DNA-directed RNA polymerase subunit alpha | -2.52 | 0.00 |
| *.* | . | -2.36 | 0.00 |
| *RL2* | 50S ribosomal protein L2 | -2.03 | 0.00 |
| *YCX91* | Uncharacterized protein ORF91 | -1.91 | 0.00 |
| *MAP3K2* | Mitogen-activated protein kinase kinase kinase 2 | -1.88 | 0.00 |
| *Tbc1d9b* | TBC1 domain family member 9B | -1.88 | 0.00 |
| *SECY* | Protein translocase subunit SecY | -1.87 | 0.00 |
| *SEPT8* | Septin-8 | -1.84 | 0.00 |
| *Sbno1* | Protein strawberry notch homolog 1 | -1.81 | 0.00 |
| *PARVA* | Alpha-parvin | -1.67 | 0.01 |
| *MTHFD1L* | Monofunctional C1-tetrahydrofolate synthase, mitochondrial | -1.66 | 0.00 |
| *Larp1* | La-related protein 1 | -1.64 | 0.00 |
| *ZBTB10* | Zinc finger and BTB domain-containing protein 10 | -1.63 | 0.00 |
| *SPTBN1* | Spectrin beta chain, non-erythrocytic 1 | -1.63 | 0.00 |
| *Myo1c* | Unconventional myosin-Ic | -1.58 | 0.00 |
| *CHD6* | Chromodomain-helicase-DNA-binding protein 6 | -1.58 | 0.00 |
| *CSF2RB* | Cytokine receptor common subunit beta | -1.51 | 0.00 |
| *HTT* | Huntingtin | -1.50 | 0.00 |
| *Ppp6c* | Serine/threonine-protein phosphatase 6 catalytic subunit | -1.50 | 0.00 |
| *B4GALT1* | Beta-1,4-galactosyltransferase 1 | -1.50 | 0.00 |
| *VPS13A* | Vacuolar protein sorting-associated protein 13A | -1.48 | 0.00 |
| *SRCAP* | Helicase SRCAP | -1.48 | 0.00 |
| *SBNO2* | Protein strawberry notch homolog 2 | -1.47 | 0.00 |
| *Itch* | E3 ubiquitin-protein ligase Itchy | -1.46 | 0.00 |
| *jmjd6-b* | Bifunctional arginine demethylase and lysyl-hydroxylase JMJD6-B | -1.46 | 0.02 |
| *nt5c2* | Cytosolic purine 5'-nucleotidase | -1.45 | 0.00 |
| *INPP5K* | Inositol polyphosphate 5-phosphatase K | -1.44 | 0.00 |
| *DOCK1* | Dedicator of cytokinesis protein 1 | -1.43 | 0.00 |
| *MYO18A* | Unconventional myosin-XVIIIa | -1.43 | 0.00 |
| *PEX14* | Peroxisomal membrane protein PEX14 | -1.42 | 0.01 |
| *.* | . | -1.41 | 0.01 |
| *KRAS* | GTPase KRas | -1.39 | 0.01 |
| *ANO5* | Anoctamin-5 | -1.38 | 0.00 |
| *Pi4ka* | Phosphatidylinositol 4-kinase alpha | -1.38 | 0.01 |
| *HECTD4* | Probable E3 ubiquitin-protein ligase HECTD4 | -1.38 | 0.00 |
| *SPAG9* | C-Jun-amino-terminal kinase-interacting protein 4 | -1.38 | 0.01 |
| *PBXIP1* | Pre-B-cell leukemia transcription factor-interacting protein 1 | -1.37 | 0.00 |
| *TNRC6C* | Trinucleotide repeat-containing gene 6C protein | -1.37 | 0.00 |
| *ywhag-a* | 14-3-3 protein gamma-A | -1.37 | 0.00 |
| *Rps6ka3* | Ribosomal protein S6 kinase alpha-3 | -1.37 | 0.00 |
| *TNFAIP2* | Tumor necrosis factor alpha-induced protein 2 | -1.36 | 0.01 |
| *Meioc* | Meiosis-specific coiled-coil domain-containing protein MEIOC | -1.36 | 0.01 |
| *paxip1* | PAX-interacting protein 1 | -1.36 | 0.01 |
| *PCNX1* | Pecanex-like protein 1 | -1.36 | 0.01 |
| *Xpo1* | Exportin-1 | -1.36 | 0.01 |
| *PPP4R1* | Serine/threonine-protein phosphatase 4 regulatory subunit 1 | -1.36 | 0.01 |
| *Pak2* | Serine/threonine-protein kinase PAK 2 | -1.35 | 0.00 |
| *HNRNPLL* | Heterogeneous nuclear ribonucleoprotein L-like | -1.35 | 0.01 |
| *tmem168* | Transmembrane protein 168 | -1.34 | 0.01 |
| *EP300* | Histone acetyltransferase p300 | -1.34 | 0.00 |
| *ASXL2* | Putative Polycomb group protein ASXL2 | -1.34 | 0.01 |
| *KIF21A* | Kinesin-like protein KIF21A | -1.34 | 0.01 |
| *KANK1* | KN motif and ankyrin repeat domain-containing protein 1 | -1.34 | 0.01 |
| *srf* | Serum response factor | -1.34 | 0.01 |
| *FERMT2* | Fermitin family homolog 2 | -1.33 | 0.00 |
| *pum1* | Pumilio homolog 1 | -1.33 | 0.00 |
| *Prrc2c* | Protein PRRC2C | -1.32 | 0.00 |
| *TRIM39* | E3 ubiquitin-protein ligase TRIM39 | -1.32 | 0.01 |
| *slc25a37* | Mitoferrin-1 | -1.32 | 0.01 |
| *PDS5A* | Sister chromatid cohesion protein PDS5 homolog A | -1.32 | 0.01 |
| *ZNF516* | Zinc finger protein 516 | -1.32 | 0.01 |
| *PIP4K2B* | Phosphatidylinositol 5-phosphate 4-kinase type-2 beta | -1.31 | 0.01 |
| *.* | . | -1.31 | 0.00 |
| *SDCBP* | Syntenin-1 | -1.31 | 0.01 |
| *DPP8* | Dipeptidyl peptidase 8 | -1.31 | 0.00 |
| *MPRIP* | Myosin phosphatase Rho-interacting protein | -1.30 | 0.01 |
| *DMD* | Dystrophin | -1.30 | 0.01 |
| *PPP1R12A* | Protein phosphatase 1 regulatory subunit 12A | -1.30 | 0.01 |
| *SETD5* | SET domain-containing protein 5 | -1.30 | 0.01 |
| *qki-b* | Protein quaking-B | -1.29 | 0.01 |
| *BICD2* | Protein bicaudal D homolog 2 | -1.29 | 0.01 |
| *timp3* | Metalloproteinase inhibitor 3 | -1.29 | 0.00 |
| *Foxp1* | Forkhead box protein P1 | -1.29 | 0.01 |
| *SLC7A3* | Cationic amino acid transporter 3 | -1.29 | 0.01 |
| *ZNF148* | Zinc finger protein 148 | -1.28 | 0.01 |
| *ADAMTS1* | A disintegrin and metalloproteinase with thrombospondin motifs 1 | -1.28 | 0.01 |
| *qsox2* | Sulfhydryl oxidase 2 | -1.28 | 0.01 |
| *NAA25* | N-alpha-acetyltransferase 25, NatB auxiliary subunit | -1.28 | 0.01 |
| *Nckipsd* | NCK-interacting protein with SH3 domain | -1.28 | 0.01 |
| *FLNB* | Filamin-B | -1.27 | 0.01 |
| *Smad6* | Mothers against decapentaplegic homolog 6 | -1.27 | 0.01 |
| *PPIP5K2* | Inositol hexakisphosphate and diphosphoinositol-pentakisphosphate kinase 2 | -1.27 | 0.01 |
| *SMARCD2* | SWI/SNF-related matrix-associated actin-dependent regulator of chromatin subfamily D member 2 | -1.27 | 0.01 |
| *Tcf12* | Transcription factor 12 | -1.26 | 0.01 |
| *COL4A1* | Collagen alpha-1(IV) chain | -1.26 | 0.01 |
| *MAP7* | Ensconsin | -1.26 | 0.01 |
| *Erc1* | ELKS/Rab6-interacting/CAST family member 1 | -1.26 | 0.01 |
| *ZFHX2* | Zinc finger homeobox protein 2 | -1.25 | 0.01 |
| *pnrc2-a* | Proline-rich nuclear receptor coactivator 2 A | -1.25 | 0.01 |
| *Ash2l* | Set1/Ash2 histone methyltransferase complex subunit ASH2 | -1.25 | 0.02 |
| *AFAP1* | Actin filament-associated protein 1 | -1.25 | 0.01 |
| *ZFHX3* | Zinc finger homeobox protein 3 | -1.25 | 0.01 |
| *MAST4* | Microtubule-associated serine/threonine-protein kinase 4 | -1.25 | 0.01 |
| *RAB5B* | Ras-related protein Rab-5B | -1.25 | 0.01 |
| *MICU2* | Calcium uptake protein 2, mitochondrial | -1.25 | 0.01 |
| *Kat6a* | Histone acetyltransferase KAT6A | -1.25 | 0.01 |
| *DAGLB* | Sn1-specific diacylglycerol lipase beta | -1.24 | 0.01 |
| *mrtfb* | Myocardin-related transcription factor B | -1.24 | 0.01 |
| *FAM46A* | Protein FAM46A | -1.24 | 0.02 |
| *slc43a2* | Large neutral amino acids transporter small subunit 4 | -1.24 | 0.01 |
| *TAF2* | Transcription initiation factor TFIID subunit 2 | -1.24 | 0.01 |
| *ATP11C* | Phospholipid-transporting ATPase IG | -1.23 | 0.01 |
| *.* | . | -1.23 | 0.01 |
| *HSPA4* | Heat shock 70 kDa protein 4 | -1.23 | 0.01 |
| *HERC2* | E3 ubiquitin-protein ligase HERC2 | -1.23 | 0.01 |
| *Slamf7* | SLAM family member 7 | -1.23 | 0.01 |
| *ATF3* | Cyclic AMP-dependent transcription factor ATF-3 | -1.22 | 0.01 |
| *Lbr* | Lamin-B receptor | -1.22 | 0.01 |
| *NCR3LG1* | Natural cytotoxicity triggering receptor 3 ligand 1 | -1.22 | 0.01 |
| *TNRC18* | Trinucleotide repeat-containing gene 18 protein | -1.22 | 0.02 |
| *cmip* | C-Maf-inducing protein | -1.22 | 0.02 |
| *NFAT5* | Nuclear factor of activated T-cells 5 | -1.22 | 0.01 |
| *pafah1b1* | Lissencephaly-1 homolog | -1.21 | 0.01 |
| *Rxra* | Retinoic acid receptor RXR-alpha | -1.21 | 0.01 |
| *RUBCN* | Run domain Beclin-1-interacting and cysteine-rich domain-containing protein | -1.21 | 0.01 |
| *ABCC1* | Multidrug resistance-associated protein 1 | -1.21 | 0.02 |
| *IKBKE* | Inhibitor of nuclear factor kappa-B kinase subunit epsilon | -1.21 | 0.00 |
| *ANKRD17* | Ankyrin repeat domain-containing protein 17 | -1.21 | 0.01 |
| *Pcnt* | Pericentrin | -1.21 | 0.01 |
| *WWTR1* | WW domain-containing transcription regulator protein 1 | -1.21 | 0.02 |
| *PLCG1* | 1-phosphatidylinositol 4,5-bisphosphate phosphodiesterase gamma-1 | -1.20 | 0.01 |
| *topbp1-A* | DNA topoisomerase 2-binding protein 1-A | -1.20 | 0.02 |
| *EMSY* | BRCA2-interacting transcriptional repressor EMSY | -1.19 | 0.01 |
| *KIF13B* | Kinesin-like protein KIF13B | -1.18 | 0.01 |
| *RIPK3* | Receptor-interacting serine/threonine-protein kinase 3 | -1.18 | 0.01 |
| *NFATC2* | Nuclear factor of activated T-cells, cytoplasmic 2 | -1.18 | 0.01 |
| *WDR11* | WD repeat-containing protein 11 | -1.18 | 0.01 |
| *MARK2* | Serine/threonine-protein kinase MARK2 | -1.18 | 0.02 |
| *HERC1* | Probable E3 ubiquitin-protein ligase HERC1 | -1.18 | 0.01 |
| *ZBTB40* | Zinc finger and BTB domain-containing protein 40 | -1.17 | 0.02 |
| *BLVRA* | Biliverdin reductase A | -1.17 | 0.03 |
| *mical3a* | Protein-methionine sulfoxide oxidase mical3a | -1.17 | 0.01 |
| *NPR3* | Atrial natriuretic peptide receptor 3 | -1.17 | 0.02 |
| *dopey2* | Protein dopey-2 | -1.17 | 0.01 |
| *pum2* | Pumilio homolog 2 | -1.16 | 0.01 |
| *RALGAPA1* | Ral GTPase-activating protein subunit alpha-1 | -1.16 | 0.02 |
| *BNIP2* | BCL2/adenovirus E1B 19 kDa protein-interacting protein 2 | -1.16 | 0.01 |
| *SPEN* | Msx2-interacting protein | -1.16 | 0.01 |
| *prkcb* | Protein kinase C beta type | -1.16 | 0.01 |
| *S1pr1* | Sphingosine 1-phosphate receptor 1 | -1.16 | 0.01 |
| *CAMK2G* | Calcium/calmodulin-dependent protein kinase type II subunit gamma | -1.16 | 0.01 |
| *Tnfrsf21* | Tumor necrosis factor receptor superfamily member 21 | -1.16 | 0.02 |
| *FV3-043R* | Uncharacterized protein 043R | -1.16 | 0.01 |
| *GRAMD1A* | GRAM domain-containing protein 1A | -1.16 | 0.02 |
| *ABI1* | Abl interactor 1 | -1.16 | 0.01 |
| *RREB1* | Ras-responsive element-binding protein 1 | -1.15 | 0.02 |
| *KIAA0232* | Uncharacterized protein KIAA0232 | -1.15 | 0.01 |
| *Thbd* | Thrombomodulin | -1.15 | 0.01 |
| *CROT* | Peroxisomal carnitine O-octanoyltransferase | -1.15 | 0.02 |
| *BCAT1* | Branched-chain-amino-acid aminotransferase, cytosolic | -1.15 | 0.01 |
| *coq10b-a* | Coenzyme Q-binding protein COQ10 homolog A, mitochondrial | -1.15 | 0.03 |
| *AKNA* | AT-hook-containing transcription factor | -1.15 | 0.02 |
| *DGKZ* | Diacylglycerol kinase zeta | -1.14 | 0.02 |
| *.* | . | -1.14 | 0.01 |
| *USP34* | Ubiquitin carboxyl-terminal hydrolase 34 | -1.14 | 0.03 |
| *TRPC4AP* | Short transient receptor potential channel 4-associated protein | -1.14 | 0.02 |
| *rnf44* | RING finger protein 44 | -1.13 | 0.03 |
| *PYCARD* | Apoptosis-associated speck-like protein containing a CARD | -1.13 | 0.03 |
| *IDE* | Insulin-degrading enzyme | -1.13 | 0.02 |
| *Pigq* | Phosphatidylinositol N-acetylglucosaminyltransferase subunit Q | -1.13 | 0.03 |
| *EEF2K* | Eukaryotic elongation factor 2 kinase | -1.13 | 0.01 |
| *UBR1* | E3 ubiquitin-protein ligase UBR1 | -1.13 | 0.02 |
| *mlec-b* | Malectin-B | -1.13 | 0.01 |
| *tshz1-b* | Teashirt homolog 1-B | -1.13 | 0.02 |
| *RIC1* | RAB6A-GEF complex partner protein 1 | -1.12 | 0.02 |
| *Higd1c* | HIG1 domain family member 1C | -1.12 | 0.01 |
| *ZEB1* | Zinc finger E-box-binding homeobox 1 | -1.12 | 0.01 |
| *NCR3LG1* | Natural cytotoxicity triggering receptor 3 ligand 1 | -1.12 | 0.02 |
| *ccdc88c* | Daple-like protein | -1.11 | 0.01 |
| *ACACB* | Acetyl-CoA carboxylase 2 | -1.11 | 0.02 |
| *Stard13* | StAR-related lipid transfer protein 13 | -1.11 | 0.02 |
| *itga5* | Integrin alpha-5 | -1.11 | 0.02 |
| *TRIM24* | Transcription intermediary factor 1-alpha | -1.11 | 0.01 |
| *VPS13D* | Vacuolar protein sorting-associated protein 13D | -1.11 | 0.02 |
| *Ahcyl1* | S-adenosylhomocysteine hydrolase-like protein 1 | -1.11 | 0.02 |
| *cwf19l1* | CWF19-like protein 1 | -1.11 | 0.01 |
| *NR2C2* | Nuclear receptor subfamily 2 group C member 2 | -1.11 | 0.03 |
| *gtpbp1* | GTP-binding protein 1 | -1.11 | 0.02 |
| *tubg1* | Tubulin gamma-1 chain | -1.11 | 0.02 |
| *Znf367* | Zinc finger protein 367 | -1.11 | 0.02 |
| *Map3k14* | Mitogen-activated protein kinase kinase kinase 14 | -1.11 | 0.02 |
| *EVI5L* | EVI5-like protein | -1.11 | 0.01 |
| *TIMP2* | Metalloproteinase inhibitor 2 | -1.11 | 0.01 |
| *Tra2b* | Transformer-2 protein homolog beta | -1.10 | 0.01 |
| *RIC1* | Receptor-type tyrosine-protein phosphatase epsilon | -1.10 | 0.02 |
| *ST3GAL1* | CMP-N-acetylneuraminate-beta-galactosamide-alpha-2,3-sialyltransferase 1 | -1.10 | 0.02 |
| *Sik3* | Serine/threonine-protein kinase SIK3 | -1.10 | 0.03 |
| *UPF1* | Regulator of nonsense transcripts 1 | -1.10 | 0.01 |
| *SLK* | STE20-like serine/threonine-protein kinase | -1.10 | 0.02 |
| *.* | . | -1.10 | 0.02 |
| *NYAP1* | Neuronal tyrosine-phosphorylated phosphoinositide-3-kinase adapter 1 | -1.10 | 0.01 |
| *TFEB* | Transcription factor EB | -1.10 | 0.02 |
| *MARE2* | Microtubule-associated protein RP/EB family member 2 | -1.10 | 0.05 |
| *top1* | DNA topoisomerase 1 | -1.10 | 0.01 |
| *ITPKB* | Inositol-trisphosphate 3-kinase B | -1.10 | 0.02 |
| *STK17A* | Serine/threonine-protein kinase 17A | -1.09 | 0.02 |
| *HUWE1* | E3 ubiquitin-protein ligase HUWE1 | -1.09 | 0.01 |
| *Papss1* | Bifunctional 3'-phosphoadenosine 5'-phosphosulfate synthase 1 | -1.09 | 0.03 |
| *Eml3* | Echinoderm microtubule-associated protein-like 3 | -1.09 | 0.02 |
| *.* | . | -1.09 | 0.01 |
| *DYSF* | Dysferlin | -1.09 | 0.02 |
| *USP9X* | Probable ubiquitin carboxyl-terminal hydrolase FAF-X | -1.08 | 0.01 |
| *top1* | DNA topoisomerase 1 | -1.08 | 0.01 |
| *PARP10* | Poly [ADP-ribose] polymerase 10 | -1.08 | 0.01 |
| *NCOR2* | Nuclear receptor corepressor 2 | -1.08 | 0.02 |
| *ccdc88c* | Daple-like protein | -1.08 | 0.03 |
| *CPNE1* | Copine-1 | -1.08 | 0.02 |
| *Ube2j1* | Ubiquitin-conjugating enzyme E2 J1 | -1.08 | 0.03 |
| *Tmco3* | Transmembrane and coiled-coil domain-containing protein 3 | -1.08 | 0.03 |
| *.* | . | -1.08 | 0.03 |
| *CD40* | Tumor necrosis factor receptor superfamily member 5 | -1.08 | 0.03 |
| *Zmiz1* | Zinc finger MIZ domain-containing protein 1 | -1.08 | 0.01 |
| *NBEAL1* | Neurobeachin-like protein 1 | -1.08 | 0.02 |
| *METTL22* | Methyltransferase-like protein 22 | -1.07 | 0.03 |
| *TRAF2* | TNF receptor-associated factor 2 | -1.07 | 0.02 |
| *golim4* | Golgi integral membrane protein 4 | -1.07 | 0.02 |
| *mybbp1a* | Myb-binding protein 1A-like protein | -1.07 | 0.02 |
| *ralbp1-a* | RalA-binding protein 1-A | -1.07 | 0.01 |
| *DPEP2* | Dipeptidase 2 | -1.07 | 0.03 |
| *HSPG2* | Basement membrane-specific heparan sulfate proteoglycan core protein | -1.07 | 0.02 |
| *ATP11B* | Probable phospholipid-transporting ATPase IF | -1.07 | 0.02 |
| *PIAS4* | E3 SUMO-protein ligase PIAS4 | -1.07 | 0.01 |
| *TRIM39* | E3 ubiquitin-protein ligase TRIM39 | -1.06 | 0.03 |
| *dym* | Dymeclin | -1.06 | 0.03 |
| *KMT2D* | Histone-lysine N-methyltransferase 2D | -1.06 | 0.04 |
| *Mycbp2* | E3 ubiquitin-protein ligase MYCBP2 | -1.06 | 0.04 |
| *SEMA6D* | Semaphorin-6D | -1.06 | 0.03 |
| *pacsin2* | Protein kinase C and casein kinase substrate in neurons protein 2 | -1.06 | 0.02 |
| *Cox10* | Protoheme IX farnesyltransferase, mitochondrial | -1.06 | 0.03 |
| *NCR3LG1* | Natural cytotoxicity triggering receptor 3 ligand 1 | -1.06 | 0.03 |
| *TCF4* | Transcription factor 4 | -1.06 | 0.02 |
| *TNS1* | Tensin-1 | -1.06 | 0.03 |
| *RSBN1* | Round spermatid basic protein 1 | -1.05 | 0.02 |
| *TANC1* | Protein TANC1 | -1.05 | 0.02 |
| *MLXIP* | MLX-interacting protein | -1.05 | 0.05 |
| *Tank* | TRAF family member-associated NF-kappa-B activator | -1.05 | 0.02 |
| *Psd4* | PH and SEC7 domain-containing protein 4 | -1.05 | 0.03 |
| *GNE* | Bifunctional UDP-N-acetylglucosamine 2-epimerase/N-acetylmannosamine kinase | -1.05 | 0.01 |
| *EFNB2* | Ephrin-B2 | -1.05 | 0.02 |
| *cfap36* | Cilia- and flagella-associated protein 36 | -1.05 | 0.03 |
| *RBMS1* | RNA-binding motif, single-stranded-interacting protein 1 | -1.05 | 0.02 |
| *LRP6* | Low-density lipoprotein receptor-related protein 6 | -1.05 | 0.02 |
| *FOSL2* | Fos-related antigen 2 | -1.04 | 0.02 |
| *BRAP* | BRCA1-associated protein | -1.04 | 0.04 |
| *ccnd2* | G1/S-specific cyclin-D2 | -1.04 | 0.02 |
| *MDN1* | Midasin | -1.04 | 0.02 |
| *Col4a2* | Collagen alpha-2(IV) chain | -1.04 | 0.01 |
| *Upp1* | Uridine phosphorylase 1 | -1.04 | 0.02 |
| *G6PD* | Glucose-6-phosphate 1-dehydrogenase | -1.04 | 0.03 |
| *tcf3* | Transcription factor E2-alpha | -1.04 | 0.01 |
| *SYNCRIP* | Heterogeneous nuclear ribonucleoprotein Q | -1.04 | 0.01 |
| *RIN2* | Ras and Rab interactor 2 | -1.04 | 0.04 |
| *SP1* | Transcription factor Sp1 | -1.04 | 0.02 |
| *IFRD1* | Interferon-related developmental regulator 1 | -1.03 | 0.02 |
| *ascc3* | Activating signal cointegrator 1 complex subunit 3 | -1.03 | 0.02 |
| *Arid1a* | AT-rich interactive domain-containing protein 1A | -1.03 | 0.04 |
| *PTK2B* | Protein-tyrosine kinase 2-beta | -1.03 | 0.04 |
| *HDAC7* | Histone deacetylase 7 | -1.03 | 0.02 |
| *UBN2* | Ubinuclein-2 | -1.03 | 0.04 |
| *Dusp4* | Dual specificity protein phosphatase 4 | -1.03 | 0.02 |
| *inf2* | Inverted formin-2 | -1.03 | 0.03 |
| *CDCA7* | Cell division cycle-associated protein 7 | -1.03 | 0.02 |
| *GPCPD1* | Glycerophosphocholine phosphodiesterase GPCPD1 | -1.03 | 0.02 |
| *ldlr-b* | Low-density lipoprotein receptor 2 | -1.02 | 0.02 |
| *HEATR3* | HEAT repeat-containing protein 3 | -1.02 | 0.03 |
| *MESDC2* | LDLR chaperone MESD | -1.02 | 0.02 |
| *slc23a2* | Solute carrier family 23 member 2 | -1.02 | 0.04 |
| *odc1-a* | Ornithine decarboxylase 1 | -1.02 | 0.02 |
| *Ulk2* | Serine/threonine-protein kinase ULK2 | -1.02 | 0.03 |
| *PHF8* | Histone lysine demethylase PHF8 | -1.02 | 0.03 |
| *STAR7* | StAR-related lipid transfer protein 7, mitochondrial | -1.02 | 0.04 |
| *SP4* | Transcription factor Sp4 | -1.02 | 0.03 |
| *KIDINS220* | Kinase D-interacting substrate of 220 kDa | -1.02 | 0.02 |
| *ZMYM2* | Zinc finger MYM-type protein 2 | -1.02 | 0.04 |
| *RHOBTB2* | Rho-related BTB domain-containing protein 2 | -1.02 | 0.02 |
| *mboat7* | Lysophospholipid acyltransferase 7 | -1.01 | 0.03 |
| *Ep400* | E1A-binding protein p400 | -1.01 | 0.02 |
| *.* | . | -1.01 | 0.02 |
| *TOB2* | Protein Tob2 | -1.01 | 0.04 |
| *FAM160A2* | FTS and Hook-interacting protein | -1.01 | 0.01 |
| *NCR3LG1* | Natural cytotoxicity triggering receptor 3 ligand 1 | -1.01 | 0.01 |
| *Atp2b1* | Plasma membrane calcium-transporting ATPase 1 | -1.00 | 0.04 |
| *.* | . | -1.00 | 0.04 |
| *FNDC3A* | Fibronectin type-III domain-containing protein 3a | -1.00 | 0.02 |
| *Tab1* | TGF-beta-activated kinase 1 and MAP3K7-binding protein 1 | -1.00 | 0.03 |
| *HIC1* | Hypermethylated in cancer 1 protein | -1.00 | 0.03 |
| *EPB41L2* | Band 4.1-like protein 2 | -1.00 | 0.04 |
| *degs1* | Sphingolipid delta(4)-desaturase DES1 | -1.00 | 0.04 |
| *pik3r5* | Phosphoinositide 3-kinase regulatory subunit 5 | -1.00 | 0.03 |
| *KIF16B* | Kinesin-like protein KIF16B | -1.00 | 0.03 |
| *CHSY1* | Chondroitin sulfate synthase 1 | -1.00 | 0.03 |
| *ddb2* | DNA damage-binding protein 2 | -1.00 | 0.03 |
| *Smg6* | Telomerase-binding protein EST1A | -1.00 | 0.02 |
| *SLC46A3* | Solute carrier family 46 member 3 | -1.00 | 0.04 |
| *PIK3CB* | Phosphatidylinositol 4,5-bisphosphate 3-kinase catalytic subunit beta isoform | -1.00 | 0.03 |
| *STAT1* | Signal transducer and activator of transcription 1 | -0.99 | 0.02 |
| *Capn15* | Calpain-15 | -0.99 | 0.04 |
| *TFB2M* | Dimethyladenosine transferase 2, mitochondrial | -0.99 | 0.03 |
| *ZDHHC3* | Palmitoyltransferase ZDHHC3 | -0.99 | 0.03 |
| *Emilin1* | EMILIN-1 | -0.99 | 0.03 |
| *FCHSD2* | F-BAR and double SH3 domains protein 2 | -0.99 | 0.02 |
| *ZBTB24* | Zinc finger and BTB domain-containing protein 24 | -0.99 | 0.03 |
| *KCNQ4* | Potassium voltage-gated channel subfamily KQT member 4 | -0.99 | 0.03 |
| *loxl2* | Lysyl oxidase homolog 2 | -0.99 | 0.03 |
| *.* | . | -0.99 | 0.04 |
| *Pycard* | Apoptosis-associated speck-like protein containing a CARD | -0.99 | 0.03 |
| *ADRB4C* | Beta-4C adrenergic receptor | -0.99 | 0.03 |
| *SPRYD3* | SPRY domain-containing protein 3 | -0.98 | 0.05 |
| *LRCH3* | Leucine-rich repeat and calponin homology domain-containing protein 3 | -0.98 | 0.03 |
| *LTBP4* | Latent-transforming growth factor beta-binding protein 4 | -0.98 | 0.04 |
| *xpr1* | Xenotropic and polytropic retrovirus receptor 1 homolog | -0.98 | 0.03 |
| *MVB12B* | Multivesicular body subunit 12B | -0.98 | 0.04 |
| *pgap1* | GPI inositol-deacylase | -0.98 | 0.04 |
| *igf2-b* | Insulin-like growth factor II-B | -0.98 | 0.03 |
| *ZFYVE26* | Zinc finger FYVE domain-containing protein 26 | -0.98 | 0.03 |
| *PDZD8* | PDZ domain-containing protein 8 | -0.98 | 0.02 |
| *.* | . | -0.98 | 0.04 |
| *ctdspl2* | CTD small phosphatase-like protein 2 | -0.98 | 0.04 |
| *kcp* | Kielin/chordin-like protein | -0.98 | 0.04 |
| *STK17A* | Serine/threonine-protein kinase 17A | -0.97 | 0.03 |
| *oxr1* | Oxidation resistance protein 1 | -0.97 | 0.04 |
| *ARHGEF17* | Rho guanine nucleotide exchange factor 17 | -0.97 | 0.04 |
| *TRI93* | E3 ubiquitin-protein ligase TRIM39 | -0.97 | 0.03 |
| *TJP1* | Tight junction protein ZO-1 | -0.97 | 0.03 |
| *CNN2* | Calponin-2 | -0.97 | 0.03 |
| *TFE3* | Transcription factor E3 | -0.97 | 0.05 |
| *MAPK6* | Mitogen-activated protein kinase 6 | -0.97 | 0.03 |
| *ARHGAP4* | Rho GTPase-activating protein 4 | -0.97 | 0.03 |
| *elp2* | Elongator complex protein 2 | -0.96 | 0.05 |
| *Pycard* | Apoptosis-associated speck-like protein containing a CARD | -0.96 | 0.03 |
| *DIP2B* | Disco-interacting protein 2 homolog B | -0.96 | 0.03 |
| *MED12* | Mediator of RNA polymerase II transcription subunit 12 | -0.96 | 0.04 |
| *DAAM2* | Disheveled-associated activator of morphogenesis 2 | -0.96 | 0.03 |
| *SMCHD1* | Structural maintenance of chromosomes flexible hinge domain-containing protein 1 | -0.96 | 0.04 |
| *RIOX2* | Ribosomal oxygenase 2 | -0.96 | 0.04 |
| *C2CD2* | C2 domain-containing protein 2 | -0.96 | 0.01 |
| *pik3r1-a* | Phosphatidylinositol 3-kinase regulatory subunit alpha | -0.96 | 0.04 |
| *inpp5d* | Phosphatidylinositol 3,4,5-trisphosphate 5-phosphatase 1 | -0.96 | 0.04 |
| *Ppp2r5e* | Serine/threonine-protein phosphatase 2A 56 kDa regulatory subunit epsilon isoform | -0.96 | 0.03 |
| *MTMR14* | Myotubularin-related protein 14 | -0.96 | 0.03 |
| *celf1* | CUGBP Elav-like family member 1 | -0.96 | 0.03 |
| *SHOC2* | Leucine-rich repeat protein SHOC-2 | -0.96 | 0.03 |
| *.* | . | -0.96 | 0.02 |
| *THEMIS* | Protein THEMIS | -0.96 | 0.04 |
| *GPR174* | Probable G-protein coupled receptor 174 | -0.96 | 0.03 |
| *si:dkey-18l1.1* | von Willebrand factor A domain-containing protein 8 | -0.96 | 0.04 |
| *NIN* | Ninein | -0.96 | 0.02 |
| *SLC6A15* | Sodium-dependent neutral amino acid transporter B(0)AT2 | -0.96 | 0.04 |
| *.* | Oocyte zinc finger protein XlCOF22 | -0.96 | 0.02 |
| *Sec23b* | HEAT repeat-containing protein 5A | -0.96 | 0.04 |
| *RAPGEF2* | Rap guanine nucleotide exchange factor 2 | -0.96 | 0.03 |
| *ICAM4* | Intercellular adhesion molecule 4 | -0.95 | 0.03 |
| *RASA2* | Ras GTPase-activating protein 2 | -0.95 | 0.05 |
| *MTMR3* | Myotubularin-related protein 3 | -0.95 | 0.05 |
| *NOS2* | Nitric oxide synthase, inducible | -0.95 | 0.02 |
| *SLAIN2* | SLAIN motif-containing protein 2 | -0.95 | 0.04 |
| *IBP1* | Insulin-like growth factor-binding protein 1 | -0.95 | 0.02 |
| *IQEC1* | IQ motif and SEC7 domain-containing protein 1 | -0.95 | 0.05 |
| *ATRX* | Transcriptional regulator ATRX | -0.94 | 0.03 |
| *cdc42se1-a* | CDC42 small effector protein 1-A | -0.94 | 0.04 |
| *POLE* | DNA polymerase epsilon catalytic subunit A | -0.94 | 0.04 |
| *twf1* | Twinfilin-1 | -0.94 | 0.04 |
| *med23* | Mediator of RNA polymerase II transcription subunit 23 | -0.94 | 0.03 |
| *DGKI* | Diacylglycerol kinase iota | -0.94 | 0.04 |
| *RRN3* | RNA polymerase I-specific transcription initiation factor RRN3 | -0.94 | 0.05 |
| *FACR1* | Fatty acyl-CoA reductase 1 | -0.94 | 0.05 |
| *MPZL1* | Myelin protein zero-like protein 1 | -0.94 | 0.03 |
| *Nsmaf* | Protein FAN | -0.94 | 0.05 |
| *CD37* | Leukocyte antigen CD37 | -0.93 | 0.04 |
| *HLA-DRB1* | HLA class II histocompatibility antigen, DRB1-4 beta chain | -0.93 | 0.02 |
| *P2RX7* | P2X purinoceptor 7 | -0.93 | 0.05 |
| *LPIN2* | Phosphatidate phosphatase LPIN2 | -0.93 | 0.03 |
| *R3hdm2* | R3H domain-containing protein 2 | -0.93 | 0.03 |
| *.* | . | -0.93 | 0.03 |
| *kpnb1* | Importin subunit beta | -0.93 | 0.05 |
| *PLEKHG5* | Pleckstrin homology domain-containing family G member 5 | -0.93 | 0.04 |
| *Limk2* | LIM domain kinase 2 | -0.93 | 0.03 |
| *Trim25* | E3 ubiquitin/ISG15 ligase TRIM25 | -0.93 | 0.04 |
| *.* | . | -0.93 | 0.02 |
| *RARA* | Retinoic acid receptor alpha | -0.93 | 0.04 |
| *ARHGAP31* | Rho GTPase-activating protein 31 | -0.93 | 0.04 |
| *SIK2* | Serine/threonine-protein kinase SIK2 | -0.93 | 0.01 |
| *yrbE* | Uncharacterized oxidoreductase YrbE | -0.93 | 0.04 |
| *med24* | Mediator of RNA polymerase II transcription subunit 24 | -0.93 | 0.04 |
| *PLEKHO2* | Pleckstrin homology domain-containing family O member 2 | -0.92 | 0.04 |
| *PQLC2* | Lysosomal amino acid transporter 1 homolog | -0.92 | 0.05 |
| *HIPK1* | Homeodomain-interacting protein kinase 1 | -0.92 | 0.04 |
| *USP32* | Ubiquitin carboxyl-terminal hydrolase 32 | -0.92 | 0.04 |
| *IQGAP1* | Ras GTPase-activating-like protein IQGAP1 | -0.92 | 0.04 |
| *MID1* | E3 ubiquitin-protein ligase Midline-1 | -0.92 | 0.03 |
| *ZNF268* | Zinc finger protein 268 | -0.92 | 0.05 |
| *ERBIN* | Erbin | -0.91 | 0.03 |
| *agap1* | Arf-GAP with GTPase, ANK repeat and PH domain-containing protein 1 | -0.91 | 0.04 |
| *ARHGAP12* | Rho GTPase-activating protein 12 | -0.91 | 0.03 |
| *.* | . | -0.91 | 0.04 |
| *L3MBTL3* | Lethal(3)malignant brain tumor-like protein 3 | -0.91 | 0.04 |
| *mdk-a* | Midkine-A | -0.91 | 0.04 |
| *DNMBP* | Dynamin-binding protein | -0.91 | 0.04 |
| *SPTLC1* | Serine palmitoyltransferase 1 | -0.91 | 0.04 |
| *CTSK* | Cathepsin K | -0.91 | 0.02 |
| *heatr6* | HEAT repeat-containing protein 6 | -0.91 | 0.03 |
| *IL7R* | Interleukin-7 receptor subunit alpha | -0.91 | 0.04 |
| *MAP4K5* | Mitogen-activated protein kinase kinase kinase kinase 5 | -0.91 | 0.04 |
| *Nlrp1b* | NACHT, LRR and PYD domains-containing protein 1b allele 3 | -0.91 | 0.04 |
| *MED13L* | Mediator of RNA polymerase II transcription subunit 13-like | -0.91 | 0.03 |
| *.* | . | -0.90 | 0.04 |
| *PPFIA3* | Liprin-alpha-3 | -0.90 | 0.02 |
| *.* | . | -0.90 | 0.05 |
| *PDE7A* | High affinity cAMP-specific 3',5'-cyclic phosphodiesterase 7A | -0.90 | 0.04 |
| *myadm* | Myeloid-associated differentiation marker homolog | -0.90 | 0.04 |
| *ZBED4* | Zinc finger BED domain-containing protein 4 | -0.90 | 0.04 |
| *Rps6ka4* | Ribosomal protein S6 kinase alpha-4 | -0.90 | 0.04 |
| *Rras2* | Ras-related protein R-Ras2 | -0.90 | 0.04 |
| *stk10* | Serine/threonine-protein kinase 10 | -0.90 | 0.04 |
| *cluh* | Clustered mitochondria protein homolog | -0.89 | 0.04 |
| *FILIP1L* | Filamin A-interacting protein 1-like | -0.89 | 0.03 |
| *.* | . | -0.89 | 0.02 |
| *Sec23b* | Protein transport protein Sec23B | -0.89 | 0.04 |
| *ICAM5* | Intercellular adhesion molecule 5 | -0.89 | 0.04 |
| *APMAP* | Adipocyte plasma membrane-associated protein | -0.89 | 0.04 |
| *ST2B1* | Sulfotransferase family cytosolic 2B member 1 | -0.89 | 0.05 |
| *AZIN1* | Antizyme inhibitor 1 | -0.89 | 0.05 |
| *TNS4* | Tensin-4 | -0.89 | 0.04 |
| *ACIN1* | Apoptotic chromatin condensation inducer in the nucleus | -0.89 | 0.05 |
| *CRAT* | Carnitine O-acetyltransferase | -0.89 | 0.02 |
| *CA8* | Carbonic anhydrase-related protein | -0.88 | 0.03 |
| *VPS35* | Vacuolar protein sorting-associated protein 35 | -0.88 | 0.05 |
| *col4a3bp* | Collagen type IV alpha-3-binding protein | -0.88 | 0.04 |
| *snrpc* | U1 small nuclear ribonucleoprotein C | -0.88 | 0.04 |
| *Cables2* | CDK5 and ABL1 enzyme substrate 2 | -0.88 | 0.03 |
| *RPRD2* | Regulation of nuclear pre-mRNA domain-containing protein 2 | -0.88 | 0.04 |
| *UBR4* | E3 ubiquitin-protein ligase UBR4 | -0.87 | 0.02 |
| *PELI1* | E3 ubiquitin-protein ligase pellino homolog 1 | -0.87 | 0.05 |
| *ME2* | NAD-dependent malic enzyme, mitochondrial | -0.87 | 0.05 |
| *.* | Tanabin | -0.87 | 0.03 |
| *Sept7* | Septin-7 | -0.87 | 0.03 |
| *Chd2* | Chromodomain-helicase-DNA-binding protein 2 | -0.87 | 0.04 |
| *SLC5A6* | Sodium-dependent multivitamin transporter | -0.87 | 0.05 |
| *FLO11* | Flocculation protein FLO11 | -0.87 | 0.05 |
| *NIPA1* | Magnesium transporter NIPA1 | -0.87 | 0.04 |
| *GCP3* | Gamma-tubulin complex component 3 homolog | -0.87 | 0.05 |
| *DNMT3A* | DNA (cytosine-5)-methyltransferase 3A | -0.87 | 0.05 |
| *pol* | Retrovirus-related Pol polyprotein from transposon 17.6 | -0.86 | 0.04 |
| *ATF7IP* | Activating transcription factor 7-interacting protein 1 | -0.86 | 0.04 |
| *MYH10* | Myosin-10 | -0.86 | 0.03 |
| *NCOA6* | Nuclear receptor coactivator 6 | -0.86 | 0.04 |
| *GPDM* | Glycerol-3-phosphate dehydrogenase, mitochondrial | -0.86 | 0.05 |
| *ITGAV* | Integrin alpha-V | -0.86 | 0.05 |
| *.* | . | -0.86 | 0.04 |
| *GLYM* | Serine hydroxymethyltransferase, mitochondrial | -0.86 | 0.05 |
| *EURL* | Protein EURL homolog | -0.86 | 0.05 |
| *PFKFB3* | 6-phosphofructo-2-kinase/fructose-2,6-bisphosphatase 3 | -0.85 | 0.04 |
| *Nt5c1a* | Cytosolic 5'-nucleotidase 1A | -0.85 | 0.05 |
| *AHR* | Aryl hydrocarbon receptor | -0.85 | 0.04 |
| *ANR11* | Ankyrin repeat domain-containing protein 11 | -0.85 | 0.05 |
| *FAAH1* | Fatty-acid amide hydrolase 1 | -0.85 | 0.05 |
| *Golm1* | Golgi membrane protein 1 | -0.85 | 0.04 |
| *Arfip2* | Arfaptin-2 | -0.85 | 0.04 |
| *apc* | Adenomatous polyposis coli homolog | -0.85 | 0.03 |
| *Arl4a* | ADP-ribosylation factor-like protein 4A | -0.84 | 0.03 |
| *ARHGAP15* | Rho GTPase-activating protein 15 | -0.84 | 0.04 |
| *PTPN9* | Tyrosine-protein phosphatase non-receptor type 9 | -0.83 | 0.05 |
| *KANSL1* | KAT8 regulatory NSL complex subunit 1 | -0.83 | 0.03 |
| *FLT1* | Vascular endothelial growth factor receptor 1 | -0.83 | 0.04 |
| *CLINT1* | Clathrin interactor 1 | -0.83 | 0.04 |
| *CAN5* | Calpain-5 | -0.82 | 0.05 |
| *CCDC50* | Coiled-coil domain-containing protein 50 | -0.82 | 0.04 |
| *USP24* | Ubiquitin carboxyl-terminal hydrolase 24 | -0.81 | 0.05 |
| *SMPD1* | Sphingomyelin phosphodiesterase | -0.81 | 0.04 |
| *PCGF5* | Polycomb group RING finger protein 5 | -0.81 | 0.05 |
| *col6a1* | Collagen alpha-1(VI) chain | -0.79 | 0.05 |
| *B3GT1* | Beta-1,3-galactosyltransferase 1 | -0.79 | 0.05 |
| *Rbm12b1* | RNA-binding protein 12B-A | -0.79 | 0.04 |
| *PARP11* | Poly [ADP-ribose] polymerase 11 | -0.79 | 0.05 |
| *stag2* | Cohesin subunit SA-2 | -0.78 | 0.05 |
| *ARFGEF2* | Brefeldin A-inhibited guanine nucleotide-exchange protein 2 | -0.77 | 0.04 |
| *HYAL1* | Hyaluronidase-1 | -0.77 | 0.04 |
| *.* | . | -0.77 | 0.04 |
| *EPG5* | Ectopic P granules protein 5 homolog | -0.76 | 0.05 |
| *.* | . | -0.75 | 0.02 |
| *Fbxo6* | F-box only protein 6 | -0.74 | 0.02 |
| *Cbfb* | Core-binding factor subunit beta | -0.73 | 0.04 |
| *USF3* | Basic helix-loop-helix domain-containing protein USF3 | -0.73 | 0.03 |
| *CCDC162P* | Coiled-coil domain-containing protein 162 | -0.73 | 0.04 |
| *HTR5B* | HEAT repeat-containing protein 5B | -0.72 | 0.05 |
| *BFAR* | Bifunctional apoptosis regulator | -0.55 | 0.04 |

Table AI5. Front (WA) up-regulation

| Gene | Protein | effect | *p* |
| --- | --- | --- | --- |
| *RALGAPA1* | Ral GTPase-activating protein subunit alpha-1 | 1.61 | 0.01 |
| *ICAM4* | Intercellular adhesion molecule 4 | 1.51 | 0.02 |
| *ADAMTSL4* | ADAMTS-like protein 4 | 1.47 | 0.01 |
| *pol* | Pol polyprotein | 1.38 | 0.01 |
| *csnk1d* | Casein kinase I isoform delta | 1.37 | 0.02 |
| *BCAT1* | Branched-chain-amino-acid aminotransferase, cytosolic | 1.35 | 0.02 |
| *mapk1* | Mitogen-activated protein kinase 1 | 1.35 | 0.02 |
| *mafb* | Transcription factor MafB | 1.33 | 0.02 |
| *Cdc42ep2* | Cdc42 effector protein 2 | 1.31 | 0.01 |
| *qki-b* | Protein quaking-B | 1.28 | 0.02 |
| *FLNB* | Filamin-B | 1.25 | 0.02 |
| *HLA-DRB1* | HLA class II histocompatibility antigen, DRB1-13 beta chain | 1.25 | 0.02 |
| *ATP13A2* | Cation-transporting ATPase 13A2 | 1.23 | 0.02 |
| *timp3* | Metalloproteinase inhibitor 3 | 1.21 | 0.02 |
| *PRSS23* | Serine protease 23 | 1.20 | 0.02 |
| *.* | . | 1.19 | 0.03 |
| *IDE* | Insulin-degrading enzyme | 1.19 | 0.03 |
| *Tmed2* | Transmembrane emp24 domain-containing protein 2 | 1.18 | 0.03 |
| *Dele* | Death ligand signal enhancer | 1.18 | 0.03 |
| *tppp3* | Tubulin polymerization-promoting protein family member 3 | 1.18 | 0.03 |
| *ITSN1* | Intersectin-1 | 1.17 | 0.03 |
| *ARHGEF17* | Rho guanine nucleotide exchange factor 17 | 1.15 | 0.03 |
| *RBMS1* | RNA-binding motif, single-stranded-interacting protein 1 | 1.15 | 0.03 |
| *Cpt1a* | Carnitine O-palmitoyltransferase 1, liver isoform | 1.15 | 0.03 |
| *gnb1* | Guanine nucleotide-binding protein G(I)/G(S)/G(T) subunit beta-1 | 1.14 | 0.03 |
| *MMRN2* | Multimerin-2 | 1.13 | 0.02 |
| *IL4R* | Interleukin-4 receptor subunit alpha | 1.13 | 0.02 |
| *.* | Class I histocompatibility antigen, F10 alpha chain | 1.11 | 0.03 |
| *JAM2* | Junctional adhesion molecule B | 1.10 | 0.03 |
| *pgap3* | Post-GPI attachment to proteins factor 3 | 1.10 | 0.04 |
| *RAB2A* | Ras-related protein Rab-2A | 1.09 | 0.03 |
| *SDPR* | Serum deprivation-response protein | 1.09 | 0.04 |
| *STK17A* | Serine/threonine-protein kinase 17A | 1.08 | 0.02 |
| *ccnd1* | G1/S-specific cyclin-D1 | 1.06 | 0.04 |
| *Msantd2* | Myb/SANT-like DNA-binding domain-containing protein 2 | 1.06 | 0.03 |
| *NTSR1* | Neurotensin receptor type 1 | 1.06 | 0.04 |
| *BICD2* | Protein bicaudal D homolog 2 | 1.06 | 0.04 |
| *OSBPL2* | Oxysterol-binding protein-related protein 2 | 1.05 | 0.04 |
| *ADA10* | Disintegrin and metalloproteinase domain-containing protein 10 | 1.05 | 0.03 |
| *magt1* | Magnesium transporter protein 1 | 1.05 | 0.04 |
| *Pigq* | Phosphatidylinositol N-acetylglucosaminyltransferase subunit Q | 1.05 | 0.05 |
| *Nagpa* | N-acetylglucosamine-1-phosphodiester alpha-N-acetylglucosaminidase | 1.05 | 0.04 |
| *SPAG9* | C-Jun-amino-terminal kinase-interacting protein 4 | 1.04 | 0.04 |
| *GAA* | Lysosomal alpha-glucosidase | 1.04 | 0.03 |
| *Arsa* | Arylsulfatase A | 1.04 | 0.04 |
| *FKBP8* | Peptidyl-prolyl cis-trans isomerase FKBP8 | 1.03 | 0.04 |
| *GDI1* | Rab GDP dissociation inhibitor alpha | 1.03 | 0.04 |
| *HLA-DRB1* | HLA class II histocompatibility antigen, DRB1-13 beta chain | 1.02 | 0.04 |
| *spns2* | Protein spinster homolog 2 | 1.01 | 0.04 |
| *dbnl-a* | Drebrin-like protein A | 1.01 | 0.04 |
| *Cd84* | SLAM family member 5 | 1.01 | 0.03 |
| *Slamf7* | SLAM family member 7 | 1.00 | 0.04 |
| *PRELID3B* | PRELI domain containing protein 3B | 1.00 | 0.05 |
| *pik3r1-a* | Phosphatidylinositol 3-kinase regulatory subunit alpha | 0.99 | 0.04 |
| *EVI5L* | EVI5-like protein | 0.97 | 0.04 |
| *HLA-DRA* | HLA class II histocompatibility antigen, DR alpha chain | 0.94 | 0.05 |
| *ATL3* | Atlastin-3 | 0.87 | 0.05 |
| *TNFRSF19* | Tumor necrosis factor receptor superfamily member 19 | 0.84 | 0.04 |

Table AI6. Front (WA) down-regulation

| Gene | Protein | effect | p |
| --- | --- | --- | --- |
| *S100A11* | Protein S100-A11 | -1.57 | 0.01 |
| *PLCL1* | Inactive phospholipase C-like protein 1 | -1.34 | 0.02 |
| *.* | . | -1.33 | 0.02 |
| *Kars* | Lysine--tRNA ligase | -1.30 | 0.03 |
| *GPI* | Glucose-6-phosphate isomerase | -1.28 | 0.02 |
| *NOP56* | Nucleolar protein 56 | -1.27 | 0.03 |
| *Paics* | Multifunctional protein ADE2 | -1.27 | 0.02 |
| *.* | . | -1.20 | 0.04 |
| *rbm8a* | RNA-binding protein 8A | -1.20 | 0.03 |
| *ZNF300* | Zinc finger protein 300 | -1.18 | 0.02 |
| *eif3a* | Eukaryotic translation initiation factor 3 subunit A | -1.17 | 0.03 |
| *PPP6R1* | Serine/threonine-protein phosphatase 6 regulatory subunit 1 | -1.16 | 0.04 |
| *SLC24A2* | Sodium/potassium/calcium exchanger 2 | -1.15 | 0.02 |
| *UBP5* | Ubiquitin carboxyl-terminal hydrolase 5 | -1.14 | 0.03 |
| *Myg1* | UPF0160 protein MYG1, mitochondrial | -1.14 | 0.03 |
| *.* | . | -1.14 | 0.04 |
| *ZNF84* | Zinc finger protein 84 | -1.14 | 0.03 |
|  | . | -1.12 | 0.03 |
| *ZNF546* | Zinc finger protein 546 | -1.11 | 0.03 |
| *NDUFB8* | NADH dehydrogenase [ubiquinone] 1 beta subcomplex subunit 8, mitochondrial | -1.10 | 0.03 |
| *Ppp1ca* | Serine/threonine-protein phosphatase PP1-alpha catalytic subunit | -1.10 | 0.04 |
| *.* | . | -1.08 | 0.03 |
| *.* | . | -1.05 | 0.02 |
| *.* | . | -1.05 | 0.05 |
| *ZBTB49* | Zinc finger and BTB domain-containing protein 49 | -1.04 | 0.04 |
| *.* | . | -1.04 | 0.04 |
| *SNRPA1* | U2 small nuclear ribonucleoprotein A' | -1.04 | 0.05 |
| *.* | . | -1.03 | 0.03 |
| *.* | . | -1.03 | 0.04 |
| *.* | . | -1.02 | 0.04 |
| *NANS* | Sialic acid synthase | -1.02 | 0.04 |
| *Dapk2* | Death-associated protein kinase 2 | -1.01 | 0.04 |
| *COX11* | Cytochrome c oxidase assembly protein COX11, mitochondrial | -1.01 | 0.05 |
| *NUP62* | Nuclear pore glycoprotein p62 | -1.00 | 0.04 |
| *.* | . | -1.00 | 0.04 |
| *.* | . | -0.99 | 0.04 |
| *Eif2b4* | Translation initiation factor eIF-2B subunit delta | -0.99 | 0.04 |
| *Cd2bp2* | CD2 antigen cytoplasmic tail-binding protein 2 | -0.99 | 0.03 |
| *.* | . | -0.98 | 0.05 |
| *COG8* | Conserved oligomeric Golgi complex subunit 8 | -0.98 | 0.05 |
| *.* | . | -0.98 | 0.05 |
| *Omg* | Oligodendrocyte-myelin glycoprotein | -0.98 | 0.04 |
| *.* | . | -0.97 | 0.04 |
| *.* | . | -0.97 | 0.05 |
| *dcaf13* | DDB1- and CUL4-associated factor 13 | -0.96 | 0.05 |
| *CC174* | Coiled-coil domain-containing protein 174 | -0.96 | 0.04 |
| *KCTD2* | BTB/POZ domain-containing protein KCTD2 | -0.96 | 0.05 |
| *rbm42* | RNA-binding protein 42 | -0.94 | 0.04 |
| *RPP38* | Ribonuclease P protein subunit p38 | -0.93 | 0.05 |
| *.* | . | -0.92 | 0.04 |
| *GNL3L* | Guanine nucleotide-binding protein-like 3-like protein | -0.92 | 0.04 |
| *ZG57* | Gastrula zinc finger protein XlCGF57.1 | -0.92 | 0.05 |
| *.* | . | -0.91 | 0.05 |
| *MRPS2* | 28S ribosomal protein S2, mitochondrial | -0.91 | 0.04 |
| *.* | . | -0.91 | 0.05 |
| *PGM1* | Phosphoglucomutase-1 | -0.90 | 0.04 |
| *.* | . | -0.89 | 0.04 |
| *DPP10* | Inactive dipeptidyl peptidase 10 | -0.88 | 0.04 |
| *Gatad2b* | Transcriptional repressor p66-beta | -0.88 | 0.05 |
| *NDUFAB1* | Acyl carrier protein, mitochondrial | -0.87 | 0.05 |
| *.* | . | -0.87 | 0.05 |
| *got2* | Aspartate aminotransferase, mitochondrial | -0.85 | 0.04 |
| *nxnl1* | Nucleoredoxin-like protein 1 | -0.84 | 0.05 |

Differentially expressed transcripts in cane toads (*Rhinella marina*) across the Australian range (main text, Figure 5). Soft clustering of RNA-Seq data from spleens was performed to visualize expression patterns across the range (main text, Figure 3). Membership indicates how closely the expression pattern of each transcript fits to that of the cluster it was grouped into.

Table AI7. Cluster 1

| Gene | Protein | Membership |
| --- | --- | --- |
| *PXN1* | Jeltraxin | 0.97 |
| *--* | -- | 0.96 |
| *REEP5* | Receptor expression-enhancing protein 5 | 0.95 |
| *FCGR2* | Low affinity immunoglobulin gamma Fc region receptor II | 0.95 |
| *ERVPABLB-1* | Endogenous retrovirus group PABLB member 1 Env polyprotein | 0.94 |
| *ARHGEF19* | Rho guanine nucleotide exchange factor 19 | 0.94 |
| *MPL* | Thrombopoietin receptor | 0.93 |
| *PXN1* | Jeltraxin | 0.93 |
| *TESPA1* | Protein TESPA1 | 0.92 |
| *lingo1* | Leucine-rich repeat and immunoglobulin-like domain-containing nogo receptor-interacting protein 1 | 0.92 |
| *P2RY12* | P2Y purinoceptor 12 | 0.91 |
| *PXN1* | Jeltraxin | 0.91 |
| *Gp9* | Platelet glycoprotein IX | 0.91 |
| *Dhrs9* | Dehydrogenase/reductase SDR family member 9 | 0.91 |
| *--* | -- | 0.90 |
| *CLUL1* | Clusterin-like protein 1 | 0.89 |
| *NMT2* | Phosphomethylethanolamine N-methyltransferase | 0.88 |
| *Gp5* | Platelet glycoprotein V | 0.88 |
| *--* | -- | 0.88 |
| *Ecm1* | Extracellular matrix protein 1 | 0.88 |
| *mul1a* | Mitochondrial ubiquitin ligase activator of nfkb 1-A | 0.87 |
| *--* | -- | 0.87 |
| *St3gal5* | Lactosylceramide alpha-2,3-sialyltransferase | 0.87 |
| *ERVW-1* | Syncytin-1 | 0.86 |
| *--* | -- | 0.86 |
| *IGKV6-21* | Immunoglobulin kappa variable 6-21 | 0.86 |
| *--* | LINE-1 reverse transcriptase homolog | 0.86 |
| *--* | -- | 0.85 |
| *CYP8B1* | 5-beta-cholestane-3-alpha,7-alpha-diol 12-alpha-hydroxylase | 0.85 |
| *CD200R1B* | Cell surface glycoprotein CD200 receptor 1-B | 0.85 |
| *--* | -- | 0.85 |
| *itln1* | Intelectin-1 | 0.84 |
| *Igk-V19-17* | Ig kappa chain V19-17 | 0.83 |
| *CYP3A29* | Cytochrome P450 3A29 | 0.83 |
| *Tub* | Tubby protein | 0.82 |
| *--* | -- | 0.82 |
| *ST3GAL6* | Type 2 lactosamine alpha-2,3-sialyltransferase | 0.82 |
| *Pol* | Pol polyprotein | 0.82 |
| *PXDC1* | PX domain-containing protein 1 | 0.82 |
| *ZNF84* | Zinc finger protein 84 | 0.82 |
| *GPR21* | Probable G-protein coupled receptor 21 | 0.81 |
| *HLA-DRB1* | HLA class II histocompatibility antigen, DRB1-4 beta chain | 0.81 |
| *--* | -- | 0.81 |
| *MED12L* | Mediator of RNA polymerase II transcription subunit 12-like protein | 0.80 |
| *FNDC7* | Fibronectin type III domain-containing protein 7 | 0.80 |
| *prr5* | Proline-rich protein 5 | 0.80 |
| *Gp1bb* | Platelet glycoprotein Ib beta chain | 0.79 |
| *--* | -- | 0.79 |
| *hacd4* | Very-long-chain (3R)-3-hydroxyacyl-CoA dehydratase | 0.79 |
| *CFH* | Complement factor H | 0.78 |
| *ENDOD1* | Endonuclease domain-containing 1 protein | 0.78 |
| *--* | -- | 0.78 |
| *Pafah2* | Platelet-activating factor acetylhydrolase 2, cytoplasmic | 0.78 |
| *ARHGEF37* | Rho guanine nucleotide exchange factor 37 | 0.78 |
| *CSF2RA* | Granulocyte-macrophage colony-stimulating factor receptor subunit alpha | 0.77 |
| *Rnf130* | E3 ubiquitin-protein ligase RNF130 | 0.76 |
| *F10* | Coagulation factor X | 0.76 |
| *PLEK* | Pleckstrin | 0.76 |
| *CCDC126* | Coiled-coil domain-containing protein 126 | 0.75 |
| *c1galt1* | Glycoprotein-N-acetylgalactosamine 3-beta-galactosyltransferase 1 | 0.75 |
| *fn1* | Fibronectin | 0.74 |
| *ITGA2B* | Integrin alpha-IIb | 0.74 |
| *--* | -- | 0.73 |
| *STON1* | Stonin-1 | 0.73 |
| *--* | -- | 0.73 |
| *Fxyd1* | Phospholemman | 0.73 |
| *Dcst1* | DC-STAMP domain-containing protein 1 | 0.72 |
| *Hmox2* | Heme oxygenase 2 | 0.71 |
| *--* | -- | 0.71 |
| *rap1b* | Ras-related protein Rap-1b | 0.71 |
| *Pinlyp* | phospholipase A2 inhibitor and Ly6/PLAUR domain-containing protein | 0.71 |
| *--* | -- | 0.70 |
| *TNFRSF6B* | Tumor necrosis factor receptor superfamily member 6B | 0.70 |
| *Esm1* | Endothelial cell-specific molecule 1 | 0.70 |

Table AI8. Cluster 2

| Gene | Protein | Membership |
| --- | --- | --- |
| *Atp5o* | ATP synthase subunit O, mitochondrial | 0.93 |
| *pkmyt1* | Membrane-associated tyrosine- and threonine-specific cdc2-inhibitory kinase | 0.89 |
| *--* | -- | 0.88 |
| *eif3a* | Eukaryotic translation initiation factor 3 subunit A | 0.88 |
| *CIR1* | Corepressor interacting with RBPJ 1 | 0.87 |
| *BEND4* | BEN domain-containing protein 4 | 0.86 |
| *--* | -- | 0.86 |
| *ZNF84* | Zinc finger protein 84 | 0.85 |
| *Kars* | Lysine--tRNA ligase | 0.85 |
| *ccdc93* | Coiled-coil domain-containing protein 93 | 0.83 |
| *--* | Gastrula zinc finger protein XlCGF57--1 | 0.83 |
| *--* | -- | 0.82 |
| *--* | -- | 0.82 |
| *MVK* | Mevalonate kinase | 0.81 |
| *exosc6* | Exosome complex component MTR3 | 0.81 |
| *Pkd1l3* | Polycystic kidney disease protein 1-like 3 | 0.81 |
| *--* | -- | 0.80 |
| *RNF40* | E3 ubiquitin-protein ligase BRE1B | 0.80 |
| *--* | -- | 0.80 |
| *RNF112* | RING finger protein 112 | 0.80 |
| *KCTD2* | BTB/POZ domain-containing protein KCTD2 | 0.79 |
| *Csnk2a2* | Casein kinase II subunit alpha' | 0.79 |
| *pgap2* | Post-GPI attachment to proteins factor 2 | 0.78 |
| *Cox11* | Cytochrome c oxidase assembly protein COX11, mitochondrial | 0.78 |
| *--* | Gastrula zinc finger protein XlCGF67--1 | 0.77 |
| *Eif2b4* | Translation initiation factor eIF-2B subunit delta | 0.76 |
| *--* | -- | 0.76 |
| *PLCL1* | Inactive phospholipase C-like protein 1 | 0.75 |
| *--* | -- | 0.75 |
| *ASB6* | Ankyrin repeat and SOCS box protein 6 | 0.75 |
| *--* | -- | 0.75 |
| *SLC24A2* | Sodium/potassium/calcium exchanger 2 | 0.74 |
| *--* | -- | 0.74 |
| *Gabarap* | Gamma-aminobutyric acid receptor-associated protein | 0.74 |
| *--* | -- | 0.74 |
| *THAP5* | THAP domain-containing protein 5 | 0.74 |
| *--* | -- | 0.74 |
| *--* | -- | 0.73 |
| *--* | -- | 0.73 |
| *LRRC41* | Leucine-rich repeat-containing protein 41 | 0.73 |
| *--* | -- | 0.73 |
| *Dapk2* | Death-associated protein kinase 2 | 0.73 |
| *VIPAS39* | Spermatogenesis-defective protein 39 homolog | 0.72 |
| *--* | -- | 0.72 |
| *UBP5* | Ubiquitin carboxyl-terminal hydrolase 5 | 0.72 |
| *spice1* | Spindle and centriole-associated protein 1 | 0.71 |
| *PSMC4* | 26S protease regulatory subunit 6B | 0.71 |
| *--* | -- | 0.70 |
| *rpl7a* | 60S ribosomal protein L7a | 0.70 |
| *IMMT* | MICOS complex subunit MIC60 | 0.70 |
| *Ier5* | Immediate early response gene 5 protein | 0.70 |
| *ELOVL1* | Elongation of very long chain fatty acids protein 1 | 0.70 |

Table AI9. Cluster 3

| Gene | Protein | Membership |
| --- | --- | --- |
| *BICD2* | Protein bicaudal D homolog 2 | 0.95 |
| *IDE* | Insulin-degrading enzyme | 0.93 |
| *mboat7* | Lysophospholipid acyltransferase 7 | 0.93 |
| *APMAP* | Adipocyte plasma membrane-associated protein | 0.91 |
| *TNS1* | Tensin-1 | 0.90 |
| *SPAG9* | C-Jun-amino-terminal kinase-interacting protein 4 | 0.89 |
| *qki-b* | Protein quaking-B | 0.88 |
| *CROT* | Peroxisomal carnitine O-octanoyltransferase | 0.88 |
| *PARVA* | Alpha-parvin | 0.87 |
| *RALGAPA1* | Ral GTPase-activating protein subunit alpha-1 | 0.87 |
| *Tab1* | TGF-beta-activated kinase 1 and MAP3K7-binding protein 1 | 0.86 |
| *Ube2j1* | Ubiquitin-conjugating enzyme E2 J1 | 0.86 |
| *RBMS1* | RNA-binding motif, single-stranded-interacting protein 1 | 0.85 |
| *Emilin1* | EMILIN-1 | 0.83 |
| *IQSEC1* | IQ motif and SEC7 domain-containing protein 1 | 0.83 |
| *FLNB* | Filamin-B | 0.83 |
| *DPEP2* | Dipeptidase 2 | 0.82 |
| *SPTLC1* | Serine palmitoyltransferase 1 | 0.82 |
| *csnk1d* | Casein kinase I isoform delta | 0.82 |
| *--* | -- | 0.82 |
| *SEPT7* | Septin-7 | 0.82 |
| *EVI5L* | EVI5-like protein | 0.82 |
| *Tmco3* | Transmembrane and coiled-coil domain-containing protein 3 | 0.81 |
| *Larp1* | La-related protein 1 | 0.81 |
| *ARHGEF17* | Rho guanine nucleotide exchange factor 17 | 0.81 |
| *CNN2* | Calponin-2 | 0.80 |
| *ITGAV* | Integrin alpha-V | 0.80 |
| *Dele* | Death ligand signal enhancer | 0.79 |
| *RIN2* | Ras and Rab interactor 2 | 0.79 |
| *Thbd* | Thrombomodulin | 0.78 |
| *ZDHHC3* | Palmitoyltransferase ZDHHC3 | 0.78 |
| *--* | -- | 0.77 |
| *WWTR1* | WW domain-containing transcription regulator protein 1 | 0.77 |
| *SDPR* | Serum deprivation-response protein | 0.76 |
| *IQGAP1* | Ras GTPase-activating-like protein IQGAP1 | 0.76 |
| *RAB2A* | Ras-related protein Rab-2A | 0.76 |
| *FKBP8* | Peptidyl-prolyl cis-trans isomerase FKBP8 | 0.76 |
| *SDCBP* | Syntenin-1 | 0.76 |
| *tppp3* | Tubulin polymerization-promoting protein family member 3 | 0.75 |
| *ccnd1* | G1/S-specific cyclin-D1 | 0.75 |
| *VGLL4* | Transcription cofactor vestigial-like protein 4 | 0.75 |
| *CRAT* | Carnitine O-acetyltransferase | 0.75 |
| *ATL3* | Atlastin-3 | 0.75 |
| *BCAT1* | Branched-chain-amino-acid aminotransferase, cytosolic | 0.75 |
| *kcp* | Kielin/chordin-like protein | 0.75 |
| *ITSN1* | Intersectin-1 | 0.74 |
| *Golm1* | Golgi membrane protein 1 | 0.74 |
| *Rras2* | Ras-related protein R-Ras2 | 0.74 |
| *Smad6* | Mothers against decapentaplegic homolog 6 | 0.74 |
| *ATP13A2* | Cation-transporting ATPase 13A2 | 0.74 |
| *ICAM5* | Intercellular adhesion molecule 5 | 0.74 |
| *Camk1* | Calcium/calmodulin-dependent protein kinase type 1 | 0.74 |
| *adam10* | Disintegrin and metalloproteinase domain-containing protein 10 | 0.73 |
| *Papss1* | Bifunctional 3'-phosphoadenosine 5'-phosphosulfate synthase 1 | 0.73 |
| *Clic4* | Chloride intracellular channel protein 4 | 0.72 |
| *myadm* | Myeloid-associated differentiation marker homolog | 0.72 |
| *C2CD2* | C2 domain-containing protein 2 | 0.72 |
| *gnb1* | Guanine nucleotide-binding protein G(I)/G(S)/G(T) subunit beta-1 | 0.72 |
| *timp3* | Metalloproteinase inhibitor 3 | 0.72 |
| *TNFAIP2* | Tumor necrosis factor alpha-induced protein 2 | 0.72 |
| *STK17A* | Serine/threonine-protein kinase 17A | 0.72 |
| *TNS4* | Tensin-4 | 0.72 |
| *--* | -- | 0.71 |
| *--* | -- | 0.71 |
| *DAAM2* | Disheveled-associated activator of morphogenesis 2 | 0.71 |
| *SPPL2A* | Signal peptide peptidase-like 2A | 0.71 |

Table AI10. Cluster 4

| Gene | Protein | Membership |
| --- | --- | --- |
| *NSD3* | Histone-lysine N-methyltransferase NSD3 | 0.95 |
| *ZNF300* | Zinc finger protein 300 | 0.95 |
| *rps8* | 40S ribosomal protein S8 | 0.95 |
| *eif3d* | Eukaryotic translation initiation factor 3 subunit D | 0.94 |
| *--* | -- | 0.93 |
| *eef1g-a* | Elongation factor 1-gamma-A | 0.92 |
| *CTDSP2* | Carboxy-terminal domain RNA polymerase II polypeptide A small phosphatase 2 | 0.92 |
| *smyd5* | SET and MYND domain-containing protein 5 | 0.89 |
| *MOCS2* | Molybdopterin synthase catalytic subunit | 0.89 |
| *Borcs6* | BLOC-1-related complex subunit 6 | 0.88 |
| *DLST* | Dihydrolipoyllysine-residue succinyltransferase component of 2-oxoglutarate dehydrogenase complex, mitochondrial | 0.88 |
| *eif3l* | Eukaryotic translation initiation factor 3 subunit L | 0.88 |
| *rps2* | 40S ribosomal protein S2 | 0.87 |
| *--* | Gastrula zinc finger protein XlCGF57--1 | 0.87 |
| *pno1* | RNA-binding protein PNO1 | 0.87 |
| *LRSAM1* | E3 ubiquitin-protein ligase LRSAM1 | 0.87 |
| *rps15* | 40S ribosomal protein S15 | 0.86 |
| *GFM2* | Ribosome-releasing factor 2, mitochondrial | 0.86 |
| *ISG20L2* | Interferon-stimulated 20 kDa exonuclease-like 2 | 0.86 |
| *Rps19* | 40S ribosomal protein S19 | 0.85 |
| *eif3h* | Eukaryotic translation initiation factor 3 subunit H | 0.85 |
| *eif3k* | Eukaryotic translation initiation factor 3 subunit K | 0.85 |
| *--* | -- | 0.84 |
| *tmem251* | Transmembrane protein 251 | 0.84 |
| *Rps27* | 40S ribosomal protein S27 | 0.84 |
| *Rpl11* | 60S ribosomal protein L11 | 0.84 |
| *Rps3* | 40S ribosomal protein S3 | 0.84 |
| *RPL35* | 60S ribosomal protein L35 | 0.83 |
| *--* | -- | 0.82 |
| *--* | -- | 0.82 |
| *Rpl12* | 60S ribosomal protein L12 | 0.82 |
| *--* | -- | 0.81 |
| *Rpl23* | 60S ribosomal protein L23 | 0.81 |
| *ndor1* | NADPH-dependent diflavin oxidoreductase 1 | 0.81 |
| *eif3g-b* | Eukaryotic translation initiation factor 3 subunit G-B | 0.81 |
| *Psmg3* | Proteasome assembly chaperone 3 | 0.79 |
| *--* | -- | 0.79 |
| *ZNF300* | Zinc finger protein 300 | 0.79 |
| *eif3b* | Eukaryotic translation initiation factor 3 subunit B | 0.79 |
| *svbp* | Small vasohibin-binding protein | 0.79 |
| *--* | -- | 0.78 |
| *eif3c* | Eukaryotic translation initiation factor 3 subunit C | 0.78 |
| *MTMR9* | Myotubularin-related protein 9 | 0.78 |
| *--* | -- | 0.78 |
| *ZNF300* | Zinc finger protein 300 | 0.77 |
| *--* | -- | 0.77 |
| *rps23* | 40S ribosomal protein S23 | 0.77 |
| *Rpl27* | 60S ribosomal protein L27 | 0.77 |
| *Serbp1* | Plasminogen activator inhibitor 1 RNA-binding protein | 0.77 |
| *RPL3* | 60S ribosomal protein L3 | 0.77 |
| *--* | -- | 0.77 |
| *Eef1d* | Elongation factor 1-delta | 0.76 |
| *--* | -- | 0.76 |
| *--* | -- | 0.76 |
| *--* | Oocyte zinc finger protein XlCOF7--1 | 0.76 |
| *--* | Cytochrome c oxidase subunit 4 isoform 2, mitochondrial | 0.75 |
| *rps3a* | 40S ribosomal protein S3a | 0.75 |
| *--* | -- | 0.75 |
| *MCUB* | Calcium uniporter regulatory subunit MCUb, mitochondrial | 0.74 |
| *EIF5A2* | Eukaryotic translation initiation factor 5A-2 | 0.74 |
| *Tmed8* | Protein TMED8 | 0.73 |
| *--* | -- | 0.73 |
| *rpl18a* | 60S ribosomal protein L18a | 0.73 |
| *Ppp1r35* | Protein phosphatase 1 regulatory subunit 35 | 0.72 |
| *--* | -- | 0.72 |
| *TPT1* | Translationally-controlled tumor protein homolog | 0.72 |
| *--* | -- | 0.72 |
| *med8-b* | Mediator of RNA polymerase II transcription subunit 8-B | 0.72 |
| *get4* | Golgi to ER traffic protein 4 homolog | 0.72 |
| *fam32a* | Protein FAM32A | 0.72 |
| *LCMT1* | Leucine carboxyl methyltransferase 1 | 0.71 |
| *--* | -- | 0.71 |
| *--* | -- | 0.70 |
| *rnf2-a* | E3 ubiquitin-protein ligase RING2-A | 0.70 |
| *NGB* | Neuroglobin | 0.70 |
| *SLC35A1* | CMP-sialic acid transporter | 0.70 |
| *rpl18-b* | 60S ribosomal protein L18-B | 0.70 |
| *Polr2c* | DNA-directed RNA polymerase II subunit RPB3 | 0.70 |

Table AI11. Cluster 5

| Gene | Protein | Membership |
| --- | --- | --- |
| *--* | -- | 0.94 |
| *NCR3LG1* | Natural cytotoxicity triggering receptor 3 ligand 1 | 0.92 |
| *Meioc* | Meiosis-specific coiled-coil domain-containing protein MEIOC | 0.91 |
| *ACACB* | Acetyl-CoA carboxylase 2 | 0.90 |
| *mrtfb* | Myocardin-related transcription factor B | 0.89 |
| *Myo1c* | Unconventional myosin-Ic | 0.88 |
| *ST3GAL1* | CMP-N-acetylneuraminate-beta-galactosamide-alpha-2,3-sialyltransferase 1 | 0.87 |
| *srf* | Serum response factor | 0.87 |
| *--* | -- | 0.86 |
| *DMD* | Dystrophin | 0.86 |
| *tcf3* | Transcription factor E2-alpha | 0.85 |
| *pum2* | Pumilio homolog 2 | 0.85 |
| *TFEB* | Transcription factor EB | 0.85 |
| *FOSL2* | Fos-related antigen 2 | 0.84 |
| *RRN3* | RNA polymerase I-specific transcription initiation factor RRN3 | 0.84 |
| *MAP3K2* | Mitogen-activated protein kinase kinase kinase 2 | 0.83 |
| *ATP11C* | Phospholipid-transporting ATPase IG | 0.82 |
| *DOCK1* | Dedicator of cytokinesis protein 1 | 0.82 |
| *pik3r5* | Phosphoinositide 3-kinase regulatory subunit 5 | 0.81 |
| *GNE* | Bifunctional UDP-N-acetylglucosamine 2-epimerase/N-acetylmannosamine kinase | 0.81 |
| *MICU2* | Calcium uptake protein 2, mitochondrial | 0.81 |
| *stk10* | Serine/threonine-protein kinase 10 | 0.80 |
| *Tank* | TRAF family member-associated NF-kappa-B activator | 0.79 |
| *SRCAP* | Helicase SRCAP | 0.79 |
| *RASA2* | Ras GTPase-activating protein 2 | 0.78 |
| *VPS13A* | Vacuolar protein sorting-associated protein 13A | 0.78 |
| *ATRX* | Transcriptional regulator ATRX | 0.77 |
| *DAGLB* | Sn1-specific diacylglycerol lipase beta | 0.76 |
| *AFAP1* | Actin filament-associated protein 1 | 0.76 |
| *Pycard* | Apoptosis-associated speck-like protein containing a CARD | 0.76 |
| *ZFYVE26* | Zinc finger FYVE domain-containing protein 26 | 0.76 |
| *RREB1* | Ras-responsive element-binding protein 1 | 0.75 |
| *PTK2B* | Protein-tyrosine kinase 2-beta | 0.75 |
| *ABI1* | Abl interactor 1 | 0.75 |
| *Sbno1* | Protein strawberry notch homolog 1 | 0.75 |
| *PPP1R12A* | Protein phosphatase 1 regulatory subunit 12A | 0.75 |
| *HNRNPLL* | Heterogeneous nuclear ribonucleoprotein L-like | 0.74 |
| *NFATC2* | Nuclear factor of activated T-cells, cytoplasmic 2 | 0.74 |
| *NR2C2* | Nuclear receptor subfamily 2 group C member 2 | 0.74 |
| *Ppp6c* | Serine/threonine-protein phosphatase 6 catalytic subunit | 0.74 |
| *Lbr* | Lamin-B receptor | 0.74 |
| *ERBIN* | Erbin | 0.74 |
| *PBXIP1* | Pre-B-cell leukemia transcription factor-interacting protein 1 | 0.73 |
| *STXBP2* | Syntaxin-binding protein 2 | 0.72 |
| *ARHGAP4* | Rho GTPase-activating protein 4 | 0.72 |
| *coq10b-a* | Coenzyme Q-binding protein COQ10 homolog A, mitochondrial | 0.71 |
| *CSF2RB* | Cytokine receptor common subunit beta | 0.71 |
| *Itch* | E3 ubiquitin-protein ligase Itchy | 0.70 |
| *PDS5A* | Sister chromatid cohesion protein PDS5 homolog A | 0.70 |

Table AI12. Cluster 6

| Gene | Protein | Membership |
| --- | --- | --- |
| *ANKRD17* | Ankyrin repeat domain-containing protein 17 | 0.91 |
| *MINA* | Bifunctional lysine-specific demethylase and histidyl-hydroxylase MINA | 0.90 |
| *odc1-a* | Ornithine decarboxylase 1 | 0.90 |
| *TRIM39* | E3 ubiquitin-protein ligase TRIM39 | 0.89 |
| *paxip1* | PAX-interacting protein 1 | 0.88 |
| *Pycard* | Apoptosis-associated speck-like protein containing a CARD | 0.87 |
| *TAF2* | Transcription initiation factor TFIID subunit 2 | 0.87 |
| *HERC2* | E3 ubiquitin-protein ligase HERC2 | 0.87 |
| *RHOBTB2* | Rho-related BTB domain-containing protein 2 | 0.86 |
| *SCAF11* | Protein SCAF11 | 0.85 |
| *UPF1* | Regulator of nonsense transcripts 1 | 0.84 |
| *pol* | Retrovirus-related Pol polyprotein from transposon 17--6 | 0.83 |
| *VPS13D* | Vacuolar protein sorting-associated protein 13D | 0.81 |
| *Arid1a* | AT-rich interactive domain-containing protein 1A | 0.81 |
| *KIF21A* | Kinesin-like protein KIF21A | 0.79 |
| *MAST4* | Microtubule-associated serine/threonine-protein kinase 4 | 0.79 |
| *ZBTB24* | Zinc finger and BTB domain-containing protein 24 | 0.78 |
| *HUWE1* | E3 ubiquitin-protein ligase HUWE1 | 0.78 |
| *COL4A1* | Collagen alpha-1(IV) chain | 0.77 |
| *ZMYM2* | Zinc finger MYM-type protein 2 | 0.77 |
| *--* | -- | 0.75 |
| *--* | -- | 0.74 |
| *GPCPD1* | Glycerophosphocholine phosphodiesterase GPCPD1 | 0.73 |
| *ccnd2* | G1/S-specific cyclin-D2 | 0.73 |
| *DNMT3A* | DNA (cytosine-5)-methyltransferase 3A | 0.73 |
| *Nlrp1b* | NACHT, LRR and PYD domains-containing protein 1b allele 3 | 0.72 |
| *MPZL1* | Myelin protein zero-like protein 1 | 0.72 |
| *mapk8* | Mitogen-activated protein kinase 8 | 0.71 |
| *Nckipsd* | NCK-interacting protein with SH3 domain | 0.71 |
| *HDAC7* | Histone deacetylase 7 | 0.71 |
| *FAM46A* | Protein FAM46A | 0.70 |
| *Xpo1* | Exportin-1 | 0.70 |

**Appendix II**

Transcripts in cane toads (*Rhinella marina*) associated with environmental variables (maximum temperature during the hottest month or rainfall during the driest quarter) across the Australian range (main text, Figure 1). A latent factor mixed model (LFMM) was performed on iqlr-transformed counts produced using RNA-Seq data from spleens in toads across the range (main text, Figure 1).

| Gene | Protein | Associated variable(s) | Temperature p-value | Rain p-value | Cluster |
| --- | --- | --- | --- | --- | --- |
| *--* | -- | temperature & rain | 1.52E-11 | 3.85E-09 | 1 |
| *CLUL1* | Clusterin-like protein 1 | temperature & rain | 1.14E-05 | 2.01E-07 | 1 |
| *PAFA2* | Platelet-activating factor acetylhydrolase 2, cytoplasmic | temperature | 2.60E-05 | none | 1 |
| *CREG1* | Protein CREG1 | temperature | 5.89E-06 | none | none |
| *ARSA* | Arylsulfatase A | temperature | 1.73E-05 | none | none |
| *QRIC1* | Glutamine-rich protein 1 | temperature & rain | 2.21E-08 | 6.42E-09 | none |
| *IIGP5* | Interferon-inducible GTPase 5 | temperature | 9.02E-06 | none | none |
| *SEPT8* | Septin-8 | temperature | 2.60E-06 | none | none |
| *DPYL3* | Dihydropyrimidinase-related protein 3 | temperature | 1.76E-05 | none | none |
| *--* | -- | temperature & rain | 1.47E-05 | 1.10E-05 | none |
| *GCYB1* | Guanylate cyclase soluble subunit beta-1 | rain | none | 1.51E-06 | none |

**Appendix III**

Groups of proportional and differentially proportional transcripts in RNA-Seq data obtained from spleens in cane toads (*Rhinella marina*) sampled across the northern Australian range. The propr package in R was used to identify pairs of transcripts that were coordinated in expression across all sampled states (main text, Figure 1).

Table AIII1. Proportional transcripts (transcripts with coordinated expression across all three states).

| Gene | Protein | Group (Figure S3) | Expression pattern associated with group | Fuzzy cluster associated with group |
| --- | --- | --- | --- | --- |
| *rnaseh2b* | Ribonuclease H2 subunit B | A | NT up | 4 |
| *RPL5* | 60S ribosomal protein L5 | A | NT up | 4 |
| *naca* | Nascent polypeptide-associated complex subunit alpha | A | NT up | 4 |
| *eif3m* | Eukaryotic translation initiation factor 3 subunit M | A | NT up | 4 |
| *--* | -- | A | NT up | 4 |
| *PLGRKT* | Plasminogen receptor (KT) | A | NT up | 4 |
| *POMP* | Proteasome maturation protein | A | NT up | 4 |
| *rps21* | 40S ribosomal protein S21 | A | NT up | 4 |
| *rps15* | 40S ribosomal protein S15 | A | NT up | 4 |
| *SNRPE* | Small nuclear ribonucleoprotein E | A | NT up | 4 |
| *RPS5* | 40S ribosomal protein S5 | A | NT up | 4 |
| *rps17* | 40S ribosomal protein S17 | A | NT up | 4 |
| *--* | -- | A | NT up | 4 |
| *RPS21* | 40S ribosomal protein S21 | A | NT up | 4 |
| *ALG2* | Alpha-1,3/1,6-mannosyltransferase ALG2 | A | NT up | 4 |
| *Atp5g3* | ATP synthase F(0) complex subunit C3, mitochondrial | A | NT up | 4 |
| *rpl22l1* | 60S ribosomal protein L22-like 1 | A | NT up | 4 |
| *ddx21-b* | Nucleolar RNA helicase 2-B | A | NT up | 4 |
| *eef1b* | Elongation factor 1-beta | A | NT up | 4 |
| *bzw1* | Basic leucine zipper and W2 domain-containing protein 1 | A | NT up | 4 |
| *Ndufa1* | NADH dehydrogenase [ubiquinone] 1 alpha subcomplex subunit 1 | A | NT up | 4 |
| *--* | -- | A | NT up | 4 |
| *--* | -- | A | NT up | 4 |
| *rpl18a* | 60S ribosomal protein L18a | A | NT up | 4 |
| *NDUFA8* | NADH dehydrogenase [ubiquinone] 1 alpha subcomplex subunit 8 | A | NT up | 4 |
| *NDUFA7* | NADH dehydrogenase [ubiquinone] 1 alpha subcomplex subunit 7 | A | NT up | 4 |
| *rps4* | 40S ribosomal protein S4 | A | NT up | 4 |
| *RPS25* | 40S ribosomal protein S25 | A | NT up | 4 |
| *Serbp1* | Plasminogen activator inhibitor 1 RNA-binding protein | A | NT up | 4 |
| *Rps15a* | 40S ribosomal protein S15a | A | NT up | 4 |
| *ATP5J* | ATP synthase-coupling factor 6, mitochondrial | A | NT up | 4 |
| *fmc1* | Protein FMC1 homolog | A | NT up | 4 |
| *eif3k* | Eukaryotic translation initiation factor 3 subunit K | A | NT up | 4 |
| *Rpl17* | 60S ribosomal protein L17 | A | NT up | 4 |
| *Rpl37* | 60S ribosomal protein L37 | A | NT up | 4 |
| *rps16* | 40S ribosomal protein S16 | A | NT up | 4 |
| *Eef1d* | Elongation factor 1-delta | A | NT up | 4 |
| *FAU* | Ubiquitin-like protein FUBI | A | NT up | 4 |
| *eif3e-a* | Eukaryotic translation initiation factor 3 subunit E-A | A | NT up | 4 |
| *rpsa* | 40S ribosomal protein SA | A | NT up | 4 |
| *RPS14* | 40S ribosomal protein S14 | A | NT up | 4 |
| *ATP5G2* | ATP synthase F(0) complex subunit C2, mitochondrial | A | NT up | 4 |
| *Rps11* | 40S ribosomal protein S11 | A | NT up | 4 |
| *eif3g-b* | Eukaryotic translation initiation factor 3 subunit G-B | A | NT up | 4 |
| *Tmem258* | Transmembrane protein 258 | A | NT up | 4 |
| *EIF3F* | Eukaryotic translation initiation factor 3 subunit F | A | NT up | 4 |
| *rpl15* | 60S ribosomal protein L15 | A | NT up | 4 |
| *MTMR9* | Myotubularin-related protein 9 | A | NT up | 4 |
| *rps20* | 40S ribosomal protein S20 | A | NT up | 4 |
| *rpl22* | 60S ribosomal protein L22 | A | NT up | 4 |
| *rpl8* | 60S ribosomal protein L8 | A | NT up | 4 |
| *RPL36* | 60S ribosomal protein L36 | A | NT up | 4 |
| *Eloc* | Elongin-C | A | NT up | 4 |
| *tmem167a* | Protein kish-A | A | NT up | 4 |
| *Rpl17* | 60S ribosomal protein L17 | A | NT up | 4 |
| *RpL13* | 60S ribosomal protein L13 | A | NT up | 4 |
| *rpl10a* | 60S ribosomal protein L10a | A | NT up | 4 |
| *rps6* | 40S ribosomal protein S6 | A | NT up | 4 |
| *RPL10* | 60S ribosomal protein L10 | A | NT up | 4 |
| *Rpl26* | 60S ribosomal protein L26 | A | NT up | 4 |
| *Rps9* | 40S ribosomal protein S9 | A | NT up | 4 |
| *ak2* | Adenylate kinase 2, mitochondrial | A | NT up | 4 |
| *nap1l1* | Nucleosome assembly protein 1-like 1 | A | NT up | 4 |
| *--* | -- | A | NT up | 4 |
| *Rps14* | 40S ribosomal protein S14 | A | NT up | 4 |
| *Rpl38* | 60S ribosomal protein L38 | A | NT up | 4 |
| *Rack1* | Receptor of activated protein C kinase 1 | A | NT up | 4 |
| *rpl31* | 60S ribosomal protein L31 | A | NT up | 4 |
| *mcts1-a* | Malignant T-cell-amplified sequence 1-A | A | NT up | 4 |
| *Rps8* | 40S ribosomal protein S8 | A | NT up | 4 |
| *PFDN1* | Prefoldin subunit 1 | A | NT up | 4 |
| *ATP5L* | ATP synthase subunit g, mitochondrial | A | NT up | 4 |
| *eef1g-a* | Elongation factor 1-gamma-A | A | NT up | 4 |
| *RPL13A* | 60S ribosomal protein L13a | A | NT up | 4 |
| *rpl4-b* | 60S ribosomal protein L4-B | A | NT up | 4 |
| *RSL24D1* | Probable ribosome biogenesis protein RLP24 | A | NT up | 4 |
| *rps23* | 40S ribosomal protein S23 | A | NT up | 4 |
| *rps24* | 40S ribosomal protein S24 | A | NT up | 4 |
| *pno1* | RNA-binding protein PNO1 | A | NT up | 4 |
| *RPL29* | 60S ribosomal protein L29 | A | NT up | 4 |
| *ATP5C1* | ATP synthase subunit gamma, mitochondrial | A | NT up | 4 |
| *--* | -- | A | NT up | 4 |
| *eif3b* | Eukaryotic translation initiation factor 3 subunit B | A | NT up | 4 |
| *Rpl23* | 60S ribosomal protein L23 | A | NT up | 4 |
| *rpl19* | 60S ribosomal protein L19 | A | NT up | 4 |
| *Lsm2* | U6 snRNA-associated Sm-like protein LSm2 | A | NT up | 4 |
| *Prdx5* | Peroxiredoxin-5, mitochondrial | A | NT up | 4 |
| *Btf3* | Transcription factor BTF3 | A | NT up | 4 |
| *Rps7* | 40S ribosomal protein S7 | A | NT up | 4 |
| *mtap* | S-methyl-5'-thioadenosine phosphorylase | A | NT up | 4 |
| *CCDC13* | Coiled-coil domain-containing protein 13 | A | NT up | 4 |
| *Rpl27* | 60S ribosomal protein L27 | A | NT up | 4 |
| *RPL3* | 60S ribosomal protein L3 | A | NT up | 4 |
| *smyd5* | SET and MYND domain-containing protein 5 | A | NT up | 4 |
| *RPLP0* | 60S acidic ribosomal protein P0 | A | NT up | 4 |
| *RPL34* | 60S ribosomal protein L34 | A | NT up | 4 |
| *Rpl14* | 60S ribosomal protein L14 | A | NT up | 4 |
| *RPL9* | 60S ribosomal protein L9 | A | NT up | 4 |
| *Rpl6* | 60S ribosomal protein L6 | A | NT up | 4 |
| *TPT1* | Translationally-controlled tumor protein homolog | A | NT up | 4 |
| *SFT2D1* | Vesicle transport protein SFT2A | A | NT up | 4 |
| *ARPC1A* | Actin-related protein 2/3 complex subunit 1A | A | NT up | 4 |
| *RPL35* | 60S ribosomal protein L35 | A | NT up | 4 |
| *mapk14* | Mitogen-activated protein kinase 14 | A | NT up | 4 |
| *Rpl11* | 60S ribosomal protein L11 | A | NT up | 4 |
| *EIF5A2* | Eukaryotic translation initiation factor 5A-2 | A | NT up | 4 |
| *Rps19* | 40S ribosomal protein S19 | A | NT up | 4 |
| *eif2a* | Eukaryotic translation initiation factor 2A | A | NT up | 4 |
| *eif3d* | Eukaryotic translation initiation factor 3 subunit D | A | NT up | 4 |
| *eif3h* | Eukaryotic translation initiation factor 3 subunit H | A | NT up | 4 |
| *Hspe1* | 10 kDa heat shock protein, mitochondrial | A | NT up | 4 |
| *RPL21* | 60S ribosomal protein L21 | A | NT up | 4 |
| *rps27a* | Ubiquitin-40S ribosomal protein S27a | A | NT up | 4 |
| *rps3a* | 40S ribosomal protein S3a | A | NT up | 4 |
| *Rps27* | 40S ribosomal protein S27 | A | NT up | 4 |
| *RPL7* | 60S ribosomal protein L7 | A | NT up | 4 |
| *Rpl32* | 60S ribosomal protein L32 | A | NT up | 4 |
| *GFM2* | Ribosome-releasing factor 2, mitochondrial | A | NT up | 4 |
| *rpl7a* | 60S ribosomal protein L7a | A | NT up | 4 |
| *eif3l* | Eukaryotic translation initiation factor 3 subunit L | A | NT up | 4 |
| *rps2* | 40S ribosomal protein S2 | A | NT up | 4 |
| *Rps3* | 40S ribosomal protein S3 | A | NT up | 4 |
| *snrpf* | Small nuclear ribonucleoprotein F | A | NT up | 4 |
| *Rpl12* | 60S ribosomal protein L12 | A | NT up | 4 |
| *EEF1A* | Elongation factor 1-alpha 1 | B | NT up | 4 |
| *EEF2* | Elongation factor 2 | B | NT up | 4 |
| *Nlrc3* | Protein NLRC3 | C | NT up | 4 |
| *Nlrc3* | Protein NLRC3 | C | NT up | 4 |
| *Pol* | LINE-1 retrotransposable element ORF2 protein | C | NT up | 4 |
| *ART2* | Putative uncharacterized protein ART2 | C | NT up | 4 |
| *ART2* | Putative uncharacterized protein ART2 | C | NT up | 4 |
| *ART2* | Putative uncharacterized protein ART2 | C | NT up | 4 |
| *--* | -- | C | NT up | 4 |
| *--* | -- | C | NT up | 4 |
| *--* | -- | C | NT up | 4 |
| *PXN1* | Jeltraxin | D | QLD down | 5 |
| *PXN1* | Jeltraxin | D | QLD down | 5 |
| *PXN1* | Jeltraxin | D | QLD down | 5 |
| *Gp9* | Platelet glycoprotein IX | E | QLD down | 5 |
| *hacd4* | Very-long-chain (3R)-3-hydroxyacyl-CoA dehydratase | E | QLD down | 5 |
| *ITGA2B* | Integrin alpha-IIb | E | QLD down | 5 |
| *CYR61* | Protein CYR61 | E | QLD down | 5 |
| *PLEK* | Pleckstrin | E | QLD down | 5 |
| *--* | -- | E | QLD down | 5 |
| *ABCD4* | ATP-binding cassette sub-family D member 4 | F | N/A | N/A |
| *Znf318* | Zinc finger protein 318 | F | N/A | N/A |
| *--* | -- | F | N/A | N/A |
| *--* | -- | F | N/A | N/A |
| *pol* | RNA-directed DNA polymerase from mobile element jockey | F | N/A | N/A |
| *--* | -- | F | N/A | N/A |
| *tc1a* | Transposable element Tc1 transposase | F | N/A | N/A |
| *hdac8* | Histone deacetylase 8 | F | N/A | N/A |
| *ACIN1* | Apoptotic chromatin condensation inducer in the nucleus | F | N/A | N/A |
| *--* | -- | F | N/A | N/A |
| *Hnrnpl* | Heterogeneous nuclear ribonucleoprotein L | F | N/A | N/A |
| *RTase* | Probable RNA-directed DNA polymerase from transposon BS | F | N/A | N/A |
| *SRRM1* | Serine/arginine repetitive matrix protein 1 | F | N/A | N/A |
| *--* | -- | F | N/A | N/A |
| *arglu1* | Arginine and glutamate-rich protein 1 | F | N/A | N/A |
| *RBM39* | RNA-binding protein 39 | F | N/A | N/A |
| *pol* | RNA-directed DNA polymerase from mobile element jockey | F | N/A | N/A |
| *FNBP4* | Formin-binding protein 4 | F | N/A | N/A |
| *HPF1* | Haze protective factor 1 | F | N/A | N/A |
| *Znf316* | Zinc finger protein 316 | F | N/A | N/A |
| *Sfswap* | Splicing factor, suppressor of white-apricot homolog | F | N/A | N/A |
| *SRSF11* | Serine/arginine-rich splicing factor 11 | F | N/A | N/A |
| *--* | -- | F | N/A | N/A |
| *Rbm25* | RNA-binding protein 25 | F | N/A | N/A |
| *CLASRP* | CLK4-associating serine/arginine rich protein | F | N/A | N/A |
| *--* | -- | F | N/A | N/A |
| *scrib* | Protein scribble homolog | F | N/A | N/A |
| *PAXBP1* | PAX3- and PAX7-binding protein 1 | F | N/A | N/A |
| *--* | -- | F | N/A | N/A |
| *--* | -- | F | N/A | N/A |
| *rbm5* | RNA-binding protein 5 | F | N/A | N/A |
| *--* | -- | F | N/A | N/A |
| *Kmt2b* | Histone-lysine N-methyltransferase 2B | F | N/A | N/A |
| *--* | Retrovirus-related Pol polyprotein from type-1 retrotransposable element R1 | F | N/A | N/A |
| *SEC16A* | Protein transport protein Sec16A | F | N/A | N/A |
| *ABCC5* | Multidrug resistance-associated protein 5 | F | N/A | N/A |
| *--* | -- | F | N/A | N/A |
| *Zbed4* | Zinc finger BED domain-containing protein 4 | F | N/A | N/A |
| *Zbed6* | Zinc finger BED domain-containing protein 6 | F | N/A | N/A |
| *SF1* | Splicing factor 1 | F | N/A | N/A |
| *FBRS* | Probable fibrosin-1 | F | N/A | N/A |
| *--* | -- | F | N/A | N/A |
| *--* | -- | F | N/A | N/A |
| *Brms1l* | Breast cancer metastasis-suppressor 1-like protein | F | N/A | N/A |
| *--* | -- | F | N/A | N/A |
| *--* | -- | F | N/A | N/A |
| *AREL1* | Apoptosis-resistant E3 ubiquitin protein ligase 1 | F | N/A | N/A |
| *--* | -- | F | N/A | N/A |
| *--* | -- | F | N/A | N/A |
| *--* | -- | F | N/A | N/A |
| *SRCAP* | Helicase SRCAP | F | N/A | N/A |
| *SLC25A28* | Mitoferrin-2 | F | N/A | N/A |
| *mfsd11* | UNC93-like protein MFSD11 | F | N/A | N/A |
| *taok2* | Serine/threonine-protein kinase TAO2 | F | N/A | N/A |
| *Srsf5* | Serine/arginine-rich splicing factor 5 | F | N/A | N/A |
| *--* | -- | F | N/A | N/A |
| *BPTF* | Nucleosome-remodeling factor subunit BPTF | F | N/A | N/A |
| *SNRNP70* | U1 small nuclear ribonucleoprotein 70 kDa | F | N/A | N/A |
| *CTC1* | CST complex subunit CTC1 | F | N/A | N/A |
| *CLK2* | Dual specificity protein kinase CLK2 | F | N/A | N/A |
| *CCNL1* | Cyclin-L1 | F | N/A | N/A |
| *HDAC6* | Histone deacetylase 6 | F | N/A | N/A |
| *ZNF653* | Zinc finger protein 653 | F | N/A | N/A |
| *R3HDM1* | R3H domain-containing protein 1 | F | N/A | N/A |
| *SF3B3* | Splicing factor 3B subunit 3 | F | N/A | N/A |
| *ASH1L* | Histone-lysine N-methyltransferase ASH1L | F | N/A | N/A |
| *cyhr1-a* | Cysteine and histidine-rich protein 1-A | F | N/A | N/A |
| *--* | Retrovirus-related Pol polyprotein from type-1 retrotransposable element R1 | F | N/A | N/A |
| *an3* | Putative ATP-dependent RNA helicase an3 | F | N/A | N/A |
| *Son* | Protein SON | F | N/A | N/A |
| *PNISR* | Arginine/serine-rich protein PNISR | F | N/A | N/A |
| *--* | UPF0469 protein KIAA0907 homolog | F | N/A | N/A |
| *CNTRL* | Centriolin | F | N/A | N/A |
